# Concordant genomic, chemical and morphological divergence reveals two cryptic species in the paper wasp *Polistes dominula* (Hymenoptera: Vespidae)

**DOI:** 10.64898/2026.11.19.766385

**Authors:** Fabian Schweitzer, Kenny Jandausch, Malte J. B. Lehmann, Luca Mehlhorn, Hannah M. Oelkuch, Luca Drescher, Amelie Kienzle, Thomas van de Kamp, Marcus Zuber, Leander Bertsch, Aglaia M. Bouma, Alina Bräunig, Finn Brunßen, Judith Frech, Maike E. Stephan, Stefan Tischendorf, Erik van der Spek, Lucy A. Wagner, Jan J. Wieringa, Jan Buellesbach, Barbara Feldmeyer, Florian Menzel, Thomas Schmitt, Jens Runge, Volker Nehring, Oliver Niehuis

## Abstract

The paper wasp *Polistes dominula* (Christ, 1791) is one of the most widespread vespid species in Europe and has become invasive in temperate regions on other continents. In Central Europe its mitochondrial genomes fall into two distinct haplotype groups with overlapping distribution. Whether these groups are reproductively isolated, and thus two bona fide species, is an open question. We sampled *P. dominula* in southern Germany and the Netherlands, where both haplotypes occur syntopically. We found that individuals with different mitochondrial haplotypes exhibit fixed differences at more than 18,000 nuclear loci and have haplotype-specific cuticular hydrocarbon profiles. They also differ subtly but consistently in wing and body shape, and in *Wolbachia* prevalence and strain identity. *Polistes dominula* is thus a complex of two biological species. However, among more than 850 wasps, twelve had ambiguous characters, three of them showing signs of hybridisation. Both reference genomes published as *P. dominula* belong to *Polistes pectoralis* (**rev. stat.**). Our new assembly is therefore the first reference genome of *P. dominula* sensu stricto. The remarkable variation in ecological, behavioural, and phenotypic traits attributed to *P. dominula* may reflect differences between the two species.

## Introduction

The European paper wasp *Polistes dominula* (Christ, 1791) is a common and widespread, primitively eusocial member of the family Vespidae. Its colonies are often founded cooperatively by several foundresses, among which a single dominant queen monopolises reproduction while the subordinate foundresses remain as potential successors [1]. The queen’s position is under constant threat, both from subordinate foundresses and from interspecific social parasites. Colony stability is maintained through complex dominance hierarchies and intricate chemical recognition. Because of these particular features, *Polistes* paper wasps have become models for the study of social evolution [2–4], and *P. dominula* in particular for social evolution and chemical ecology. Although it lacks the individual face recognition of *P. fuscatus* [5], *P. dominula* uses its variable facial patterns as visual cues in nestmate recognition while colonies are small [6].

Native to Mediterranean Europe, *P. dominula* has not only expanded its range dramatically northwards since the 1980s [7–9] but has also established invasive populations in temperate regions worldwide, including North America, South America, South Africa, Australia, and New Zealand [10]. The status of *P. dominula* as a single species was called into question when barcoding in Germany revealed two sympatrically occurring mitochondrial haplotype groups with corresponding Barcode Index Numbers (BINs) [11]: BOLD:AAA9495 and BOLD:AAB7105 [12] — the two groups that we resolve here as separate species, *P. dominula* sensu stricto and *P. pectoralis*, respectively. In the same study, barcoding sequences of the 5′ region of the mitochondrial cytochrome *c* oxidase subunit I gene (COI-5P) were found to be paraphyletic with respect to the established species *Polistes bucharensis* Erichson, 1849. The two BINs differ in their broad-scale European distribution [12]: BOLD:AAA9495 occupies a south-western range and is genetically closest to a Moroccan lineage, whereas BOLD:AAB7105 occupies a south-eastern range documented from Azerbaijan to Italy. Both BINs occur throughout Germany. Neumeyer et al. [13], working in Switzerland where the two BINs also co-occur, proposed that the two haplotype groups reflect divergence in two distinct Pleistocene refugia. Their present-day ranges are consistent with that scenario and would place the refugium of BOLD:AAA9495 on the Iberian Peninsula and that of BOLD:AAB7105 on the Italian or the Balkan Peninsula, although the phylogeographic history of *P. dominula* mtDNA has not been reconstructed directly. The isolation of insect lineages in separate refugia has produced a range of outcomes upon secondary contact, from highly structured populations [14] and rare hybridisation due to cytoplasmic incompatibility [15] to complete reproductive isolation [16]. Neumeyer et al. [13] found no evidence of nuclear differentiation based on the Internal Transcribed Spacer 1 (ITS1) nucleotide sequence. Although ITS1 resolved two main clades in the gene tree, clade membership did not correspond one-to-one with mitochondrial haplotype group: while BOLD:AAA9495 individuals fell exclusively into one clade, BOLD:AAB7105 individuals were distributed across both clades (see Tab. S1 and Tab. S2 for assignments). This pattern was interpreted as evidence against a reproductive barrier, although incomplete lineage sorting or introgression from rare hybridisation events could also account for it. Because *P. dominula* is widely used as a model system, the question of whether the two clades are reproductively isolated from one another represents a notable knowledge gap.

Reproductive barriers that separate insect gene pools operate through a wide variety of mechanisms, acting either before fertilisation (prezygotic) or after it (postzygotic). Several postzygotic mechanisms interact in unique ways in Hymenoptera because of their haplodiploid sex-determination system. Although F_1_ hybrid females can be viable and fertile, their male offspring (F_2_ males) are often inviable because the heterozygosity that buffers Dobzhansky– Muller incompatibilities in diploids is absent in haploid males [17]. The daughters of F_1_ females, by contrast, are diploid: the genome they inherit from their father can mask recessive incompatibilities, so their fitness reduction can be less severe than their brothers’, as shown in the chalcidoid wasp genus *Nasonia* [18] and in hybridising species of the ant genus *Formica* [19]. Another factor known to affect reproduction across arthropods, hymenopterans included, is the endosymbiotic bacterium *Wolbachia*, which can disrupt reproduction between infected and uninfected individuals in various ways [20]. In *Polistes dominula*, the mechanism inferred from infection patterns is cytoplasmic incompatibility (CI) [21], and because both mitochondria and *Wolbachia* are maternally inherited, any fitness advantage that infected females have in a population can lead to the over-representation of their mitochondrial haplotype [22].

Like other eusocial hymenopterans, *Polistes* rely on cuticular hydrocarbons (CHCs), which form the primary barrier against desiccation and at the same time act as semiochemicals that mediate a range of social interactions, including nestmate recognition and courtship signalling [23–25]. Each individual biosynthesizes its own hydrocarbon blend, and because these blends differ consistently between species, CHC profiles are potentially informative for chemotaxonomic purposes. The sex- and colony-level variation they also carry is, for that purpose, within-species variance that any such approach must accommodate. *Polistes*, however, also deposit these cues into their environment: females distribute CHCs onto the nest surface by rubbing their metasoma against the nest material [26], while males mark territories at mating leks by rubbing their mandibles, legs, and metasoma against the substrate [27,28].

A prominent feature of many *Polistes* species is cooperative nest founding, which can involve multiple reproductive females that later compete for dominance [8]. In Europe, this social system is exploited by a complex set of interspecific usurpers, including obligate socioparasites of the subgenus *Sulcopolistes*, such as *Polistes sulcifer* and *Polistes atrimandibularis* [29], as well as facultative socioparasites like *Polistes nimpha* [30]. Because CHCs mediate nestmate recognition, *Sulcopolistes* usurpers are thought to suppress biosynthesis of their own species-specific CHCs and acquire the host’s profile through direct interindividual contact and contact with the nest surface [31]. Because CHC blends can be acquired from the social environment as well as biosynthesized, chemotaxonomic inference that rests on them alone can be misleading.

Resolving the status of the two *P. dominula* clades mentioned above also requires determining to which of them the name *P. dominula* applies. Christ’s (1791) original description of the species as *Vespa dominula* [32] predates the modern concept of type material. Many of Christ’s species descriptions were likely based on the collection of Johann Christian Gerning, as there is a high congruence between Christ’s descriptions and the presence of specimens in the Gerning material. The collection has remained partially intact and is accessible at the Museum Wiesbaden, Germany. The two *Polistes* species Christ described — *P. dominula* and *P. nimpha* — are each represented by two female specimens in the Gerning collection, making these specimens the strongest candidates for being syntypes of the two species.

In the present study, we address the question of whether the two *Polistes dominula* clades are reproductively isolated where they co-occur in Central Europe. To this end, we collected wasps of this taxon from two European countries and determined their COI haplotypes (Fig. S1). For one German population, we generated genome-wide Pool-Seq SNP data and established a panel of private alleles for nuclear genotyping. For most samples, we compared CHC profiles using unsupervised (PCA) and discriminant (N-CPLS-DA) methods. For selected samples, we compared soma and wing morphology as well as *Wolbachia* strain identity and prevalence. We also genotyped individuals whose COI haplotypes and CHC chemotypes were discordant, to search for evidence of hybridisation and mitochondrial introgression. We generated a new reference genome assembly for *P. dominula* sensu stricto and fixed the name to this species by designating as lectotype the one syntype at the Museum Wiesbaden that both somatic morphology and CHC chemotype assign to it. The type specimen of the oldest available subjective synonym, *Polistes pectoralis* Herrich-Schäffer, 1841 (**rev. stat.**), is presumably lost [12], so we assigned this name to the second species by depositing a designated neotype at the Leibniz Institute for the Analysis of Biodiversity Change (LIB: ZFMK-TIS-89039).

## Materials and Methods

### Sampling

We assembled a set of 212 males and 643 females of the *Polistes dominula* complex, collected from 36 sampling locations in Germany and 35 in the Netherlands between April and September 2019–2024 (see Supplementary File 1 and Fig. S1). Throughout, the BIN labels BOLD:AAA9495 and BOLD:AAB7105 refer only to mitochondrial haplotypes, the names *P. dominula* and *P. pectoralis* to the two species, and the *P. dominula* complex to both. Wasps were identified in the field on species-diagnostic characters [12] and freeze-killed at −80 °C if used for genome sequencing and at −20 °C otherwise. We recorded stylopisation by *Xenos vesparum* (Strepsiptera), treated below as a potential non-taxonomic source of chemical variation, and collected no more than one individual per nest where nests were accessible, to minimise pseudoreplication. Two of the females are museum specimens from the Gerning Collection in Wiesbaden, likely the syntypes on which the species description is based.

### Gas Chromatography–Mass Spectrometry

We extracted cuticular profiles from the sampled wasps and from both syntypes of *P. dominula*, except two males (ON_15380 and ON_15382) that we kept intact for long-read sequencing and Hi-C scaffolding, and 852 individuals yielded CHC profiles suitable for analysis. Wasps were submerged in *n*-hexane, the solvent volume matched to specimen size. Both syntypes were treated identically and needed a longer extraction, after which only one of them yielded usable data. One individual (ON_16086) was extracted three further times to test whether its unexpected chemotype was reproducible, and the four extracts were analysed separately. Extracts were analysed by Gas Chromatography–Mass Spectrometry (GC–MS) on an Agilent 7890B chromatograph coupled to a 5977B quadrupole mass spectrometer with electron ionisation at 70 eV (Details in Supplementary Methods).

### Chemometric Analysis

Raw GC–MS data in Agilent ChemStation format were processed in Julia version 1.12.6 [33] using JuChrom.jl version 0.0.1 [34], an in-house package available on GitHub (https://github.com/oniehuis/JuChrom.jl). Files were parsed using the module JuChrom.ChemStationMSLoader. For each MS scan, retention time, m/z values, and ion-count intensities were retained. M/z values were binned with JuChrom.binmzvalues(), using the default JuChrom.integer() binning rule, floor (m/z + 0.3), so that integer bin *n* contained values in the interval [n − 0.3, n + 0.7), as recommended by O’Callaghan et al. [35]. Ion counts assigned to the same integer m/z bin within a scan were summed, and only m/z bins 29–562 were retained.

The resulting chromatograms were then made comparable across runs; each step is described in full in the Supplementary Methods. Retention times were converted to retention indices (RIs) [36] using *n*-alkane standards run on the same instrument within days of the samples. Baselines were removed (arPLS) [37], and all runs were aligned on a common grid of 1,819 retention-index bins (RI 2100–3800) by 534 m/z channels. The injection volume was constant, but the amount of cuticular material it carried varied with wasp size, solvent volume and extraction temperature, so we analysed relative rather than absolute chemical composition: each chromatogram was transformed to centred log-ratios (CLR) [38], which removes multiplicative differences in total signal between runs. A measurement variance, estimated from the alkane standards, was propagated through every step, and the transformed data were whitened with it, so that each feature contributes in proportion to the precision of its measurement.

We explored these data by Principal Component Analysis (PCA) with the Julia package MultivariateStats version 0.10.4, fitted without reference to barcode identity. We then tested whether chemical profiles differ between BINs with N-way Canonical Partial Least Squares discriminant analysis (N-CPLS) [39], implemented in the in-house package NCPLS.jl version 0.0.1 [40], available on GitHub (https://github.com/oniehuis/NCPLS.jl). Female and male wasps were analysed separately. Each model was fitted to the three-way chemical data (sample × retention index × m/z) and to the barcode labels together with three covariates: sampling location, the absolute distance of the sampling date from midyear, and stylopisation status. Samples were weighted inversely to the frequency of their BIN, so that both BINs carried equal weight in the fit despite their unequal sizes. Performance was estimated by nested cross-validation [41], with five outer folds for testing and four inner folds for choosing between one and four latent variables, and is reported as class-balanced accuracy, in which both BINs count equally. Its significance was assessed against 1,000 repetitions of the complete workflow on permuted BIN labels, with the empirical *p*-value computed as recommended by Phipson and Smyth [42]. To identify individuals whose chemistry contradicts their barcode, we repeated the hold-out testing 1,000 times with the number of latent variables fixed at the cross-validated value and recorded each sample’s misassignment ratio: the proportion of its hold-out tests in which it was assigned to the other BIN. Samples lacking any of the covariates, including the syntype, could not be used to fit the models; instead, they were projected into the score space of the final models, which predicted each sample’s group membership from its chemical profile.

To see which retention indices carry the separation, we derived a discriminant coefficient profile from each final model. N-CPLS gives each BIN a regression coefficient for every retention index and m/z of the chemical data. We subtracted the coefficients for BOLD:AAB7105 from those for BOLD:AAA9495 and summed the difference over m/z (Supplementary Methods). This gives one discriminant coefficient per retention index. Its sign shows which BIN a higher relative signal there favours (positive for BOLD:AAA9495, negative for BOLD:AAB7105), and its absolute value shows how strongly. The substances at the two highest and the two lowest discriminant coefficients of each sex’s profile served as the starting point for extracting characters for species identification, outlined in the following two paragraphs.

We assessed whether the two species can be identified from individual chromatograms by the relative height of two neighbouring peaks. In each sex, one of the four substances at the extremes of the discriminant coefficient profile was paired with a neighbouring peak whose discriminant coefficient has the opposite sign, 7-MeC27 (RI 2738.1) with the 13-;11-MeC27 peak (RI 2730.1) in females and 11,15-diMeC33 (RI 3351.5) with 5,15-diMeC33 (RI 3373.2) in males. The height of each peak was taken as the signal of the baseline-corrected Total Ion Current Chromatogram (TICC) at the peak’s median apex RI over all runs, irrespective of species membership, interpolated linearly between the two neighbouring scans. The TICC signal ratio of each pair (13-;11-MeC27 to 7-MeC27 in females, 11,15-diMeC33 to 5,15diMeC33 in males) served as a predictor of species membership, with a threshold set to maximise balanced accuracy. Predictive performance was estimated by Leave-One-Out Cross-Validation (LOOCV), with the threshold set anew without the held-out run, and by the Area Under the Curve (AUC) of the ratio’s Receiver Operating Characteristic (ROC).

We also performed selective substance detection to find qualitative chemical differences between the two species and recorded the presence of each of these eight substances (bold in Tab. S3). A substance counted as present when, for each of its detection ions, the extractedion trace, normalised to the run’s total ion current, rose at the substance’s RI further above its local baseline than the same trace rises by chance at RI where the substance does not elute (Details in Supplementary Methods). Detection limits were set from all runs, irrespective of species membership.

The figures of representative total ion current chromatograms and N-CPLS discriminant coefficient profiles (Fig. S2, Fig. S3) were generated with Makie.jl version 0.24.10 [43]. The resulting vector graphics were subsequently refined in Inkscape version 1.2.2 [44]. Major substances were identified for females and males of both species, and so were the substances at the extremes of the discriminant coefficient profiles, however small their peaks (Tab. S3).

### DNA Extraction

DNA was extracted from muscle tissue of the mesosoma with the Quick-DNA Miniprep Plus Kit (Zymo Research). We followed the kit protocol, except that proteinase K digestions were carried out overnight. DNA was eluted in 50 µL of elution buffer and stored at −20 °C until further use.

### COI Barcoding

We PCR-amplified the mitochondrial COI barcoding fragment (COI-5P) with the LCO1490-JJ / HCO2198-JJ primer pair [45], and products were unidirectionally Sanger-sequenced by Macrogen Europe (Amsterdam, The Netherlands). Nucleotide sequences were extracted from chromatograms and assigned to Barcode Index Numbers (BINs) [11] through the BOLD Identification Engine API [46,47]. A sequence was assigned to a BIN only if all ten of its closest matches agreed, which was consistently the case (Details in Supplementary Methods).

### Sequencing and Assembly of the *P. dominula* Genome

Two adult males from Schallstadt, Germany, assigned to BOLD:AAA9495 by COI barcoding, were used for long read assembly and scaffolding. One provided PacBio HiFi data from ultra-low-input libraries on a Sequel II, the other Arima Hi-C libraries for scaffolding; both were sequenced at the laboratory centre of the LOEWE Centre for Translational Biodiversity Genomics (TBG) at Senckenberg. Non-mitochondrial reads [48–50] were assembled, Hi-C scaffolded and taxonomically partitioned in three rounds, keeping reads that mapped to contigs with a hymenopteran or no taxonomic assignment [51–53]. The assembly was soft-masked [54–56] and assessed for gene-space completeness, contiguity and quality [57–61] (Tab. S4); it is available through ENA (GCA_965197175.2). We aligned it to the two assemblies previously published as *P. dominula*, GCF_001465965.1 [62] and GCA_049000335.1 [63], referred to as *Pdom_r1.2(USA)* and *Pdom_raw_medaka(NZ)*, to lift the private-allele panel’s (see below) SNP coordinates over, and computed pairwise distances among available *Polistes* assemblies [64,65] (Details in Supplementary Methods).

### Genomic Pool-Seq

We focused on two Freiburg sites (Littenweiler and Eschholzpark) with roughly similar abundances of the two BINs, and selected an equal number of individuals per haplotype from each, excluding wasps parasitised by *Xenos* and samples of insufficient DNA concentration. Each site contributed 27 individuals per haplotype (108 in total; 54 per haplotype), pooled in equimolar amounts into two balanced pools. The libraries were sequenced on an Illumina NextSeq 500 (StarSEQ GmbH, Mainz, Germany) to a target of 15 Gb per pool, giving roughly 50 million paired-end reads each. Reads were mapped to the new assembly (GCA_965197175.2) and screened for SNPs [66–69]. Differentiation was assessed at each SNP by Fisher’s exact test and across sliding windows with fixation indices (F_st_). Fisher’s exact test results were corrected for false discovery rate by the Benjamini-Hochberg procedure (λ = 0), giving a 1% threshold of *p* < 0.003, and Bonferroni-corrected [70] across 2,164,621 SNPs to *p* < 2.31e-08; correction and genome-wide visualisation were carried out in R [71–74]. We identified SNPs monomorphic in one pool and fixed for the alternative allele in the other; these constitute the private-allele panel used for individual genotyping and comparison with the reference assemblies previously published as *P. dominula* (Details in Supplementary Methods).

### Individual Genome Sequencing

We selected individuals for genome skimming on discordance between chemotype and mitochondrial haplotype: females with a misassignment ratio above 50% in the outlier scans, females whose BIN membership the final N-CPLS-DA model predicted incorrectly, and males with any misassignment at all, plus the only BOLD:AAB7105 female from the Netherlands (ON_16126). DNA extracts from twelve individuals were sequenced on an Illumina NextSeq 2000 by StarSEQ GmbH (Mainz, Germany) or Macrogen Europe (Amsterdam, The Netherlands), giving roughly 50 million 150 bp reads each. Reads were processed with the Pool-Seq pipeline above and allele states called at the private allele panel [66–68,75], then classified per site as homozygous for the Pool A allele, homozygous for the Pool B allele, heterozygous for the two, or other. Local heterozygosity was summarised per individual as the proportion of heterozygous sites in a centred sliding window over the panel SNPs, ordered along the N90 scaffolds (Details in Supplementary Methods).

### Wing Morphometrics

Right forewings of 200 females (97 BOLD:AAA9495, 103 BOLD:AAB7105) were photographed with identical setup and parameters. After digital correction for lens distortion [76–78], nineteen landmarks, mostly at vein intersections, were recorded in randomised sample order [79–81] following Bustamante et al. [82], itself based on Francoy et al. [83] (Fig. S4). Shape variation was analysed by generalised Procrustes analysis, PCA and Canonical Variate Analysis (CVA), with 10,000-iteration permutation tests for differences between BINs and Leave-One-Out CrossValidation (LOOCV) on the PCs explaining 95% of cumulative variance [72,73,84] (Details in Supplementary Methods).

### 3D Somatic Morphometrics

We selected 40 wasps for 3D imaging, aiming at 20 per BIN on whether a heptacosene (C27:1), an alkene of *P. pectoralis* females, could be detected by GC–MS, and confirmed their BIN haplotype by barcoding afterwards. Both suspected syntypes of *P. dominula* were included (Fig. S5). Specimens were scanned by Synchrotron Radiation-based Micro-Computed Tomography (SR-µCT) at the IMAGE Beamline of the KIT Light Source [85,86], at an effective pixel size of 6.1 µm and in four to five height steps per wasp. Tomographic reconstructions yielded both phaseand absorption-contrast datasets [87,88], which were blended and converted into an 8-bit volume per height step and then registered and merged to generate unified 3D volumes per individual [89,90]. Each of the fifteen distance characters was measured three times per individual and averaged, following Wurdack et al. [91] and defined in Tab. S5 and Fig. S6. Isosize, a shape Principal Component Analysis (PCA) and Linear Discriminant Analysis (LDA) ratio extraction were computed [72,73,92]; the PCA projection is coloured by BIN identity and the LDA by species assignment, to account for mitochondrial introgression. We evaluated the four most BIN-discriminative length ratios by Receiver Operating Characteristic (ROC) curves and the Area Under the Curve (AUC) (Details in Supplementary Methods).

### *Wolbachia* Multi Locus Sequence Typing

We determined *Wolbachia* infection status, prevalence and strain identity by amplicon-based multilocus sequence typing (MLST) [93], amplifying the five standard markers (gatB, coxA, hcpA, ftsZ, fbpA) and wsp from the 108 Pool-Seq individuals (Littenweiler and Eschholzpark, Freiburg) with the recommended primers (Tab. S6) and Sanger-sequencing them at Macrogen Europe (Amsterdam, The Netherlands). Alleles were typed and strains assigned to sequence types (STs) through PubMLST [94], whenever unambiguously possible; individuals with at least one successfully typed marker were classified as infected.

## Results

### COI Barcoding

We obtained 853 individual COI-5P sequences, 207 belonging to BOLD:AAA9495 and 646 to BOLD:AAB7105. BOLD:AAA9495 dominated in the Netherlands (60 of 61 sequences), where BOLD:AAB7105 was almost absent. In Germany, BOLD:AAB7105 was prevalent (645 of 792 sequences) and BOLD:AAA9495 widespread but less abundant (147 of 792 sequences; Fig. S1).

### Chemometric Analysis

Principal Component Analysis (PCA) of the whitened chemical data (scaled so that each feature contributes in proportion to the precision of its measurement) comprising all available female wasps (N = 645: 167 of BOLD:AAA9495, 477 of BOLD:AAB7105, and one nonbarcoded syntype) separated the majority of females assigned to different BINs along the third principal component (PC3), although several outliers were present in both barcode groups (Fig. S7A). Of the three components, only PC3 distinguished the barcode groups (AUC = 0.97, 95% CI 0.96–0.99), whereas the first component did not (AUC = 0.52, 95% CI 0.47–0.56). Seasonal variation loaded on a different axis: PC2 correlated with day of year (*r* = −0.51), while PC3 showed no detectable association with it (*r* = −0.04, *p* = 0.912; Tab. S7). The corresponding analysis of male wasps (N = 210: 41 of BOLD:AAA9495, 169 of BOLD:AAB7105) also revealed differentiation between the two barcode groups, in this case along the second principal component (PC2), although some overlap remained when this axis was considered in isolation. Simultaneous consideration of the first and second principal components separated the two groups without overlap (Fig. S7B).

Our implementation of the N-CPLS discriminant analysis includes known non-taxonomic sources of chemical variation, namely sampling location, season, and stylopisation status, alongside the barcode assignment, so that the resulting axes describe the chemical contrast between the barcode groups rather than these potential covariates. Application of this analysis to the whitened chemical data of all females (N = 635; 158 of BOLD:AAA9495, 477 of BOLD:AAB7105), clearly separated wasps of the two barcode groups in the score space spanned by the first two latent variables (Fig. 1A), with very few outliers. Nested cross-validation selected two latent variables in four of the five outer folds and three latent variables in the remaining fold. The mean class-balanced cross-validated accuracy was 99.1%, and no permutation run reached this value (1,000 permutations; *p* < 0.001). The outlier scan based on a model with two latent variables identified eight samples with a misassignment ratio greater than 50% (Supplementary File 2). Particularly notable were samples ON_13593 and ON_15008, which were assigned to the incorrect BIN in 100% of the tests, and sample ON_13555, which was assigned to the incorrect BIN in 86% of the tests. Projection of an additional ten samples with known barcode identity (all BOLD:AAA9495) placed six samples within the score space of the corresponding barcode group, BOLD:AAA9495. The exception was one wasp, ON_16086, including its three technical replicates obtained from repeated cuticular surface extraction of the same wasp. All four chemical profiles of this wasp projected into the score space occupied by samples of barcode group BOLD:AAB7105, despite the wasp carrying the barcode of BOLD:AAA9495.

**Fig. 1:**
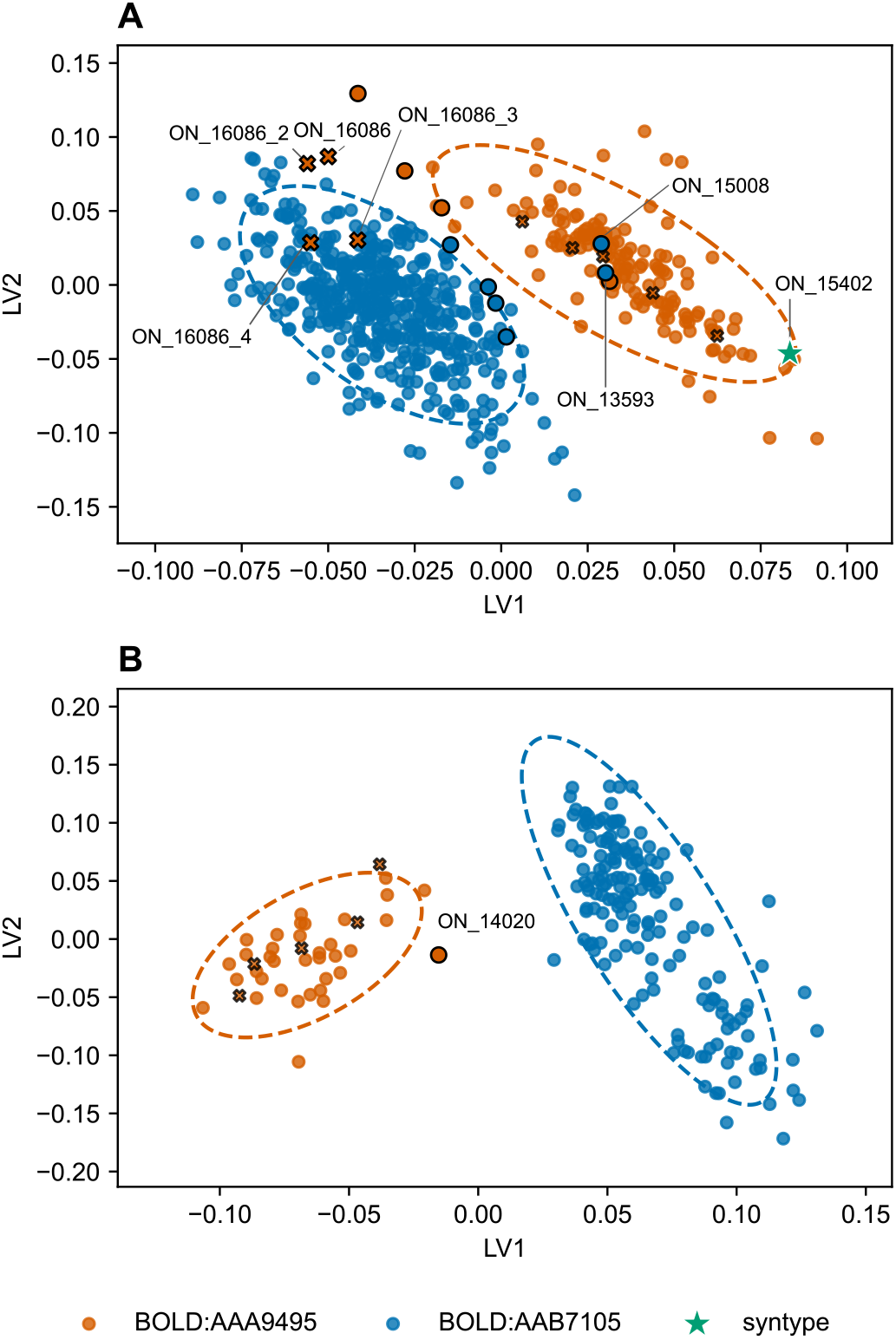
Score plots showing the separation of female (**A**: n = 645) and male (**B**: n = 210) wasps in the *Polistes dominula* species complex along the first two latent variables (LV1, LV2) of N-CPLSDA models based on their cuticular surface profiles. Points are coloured by group identity: BOLD:AAA9495 in orange, BOLD:AAB7105 in blue, and the syntype ON_15402 as a green star. Model-fitting samples are shown as filled circles and projected samples as crosses, with dashed ellipses denoting the 95% confidence region of each barcode group. Points outlined in black are also included in the individual genotyping panel (Fig. 2); individuals with noteworthy phenotype-genotype combinations are marked with their sample IDs, and labels suffixed _2 to_4 denote technical replicates of wasp ON_16086. **A**: 635 females (158 BOLD:AAA9495, 477 BOLD:AAB7105) were used for model fitting and a further ten projected, including the syntype ON_15402. **B**: 205 males (36 BOLD:AAA9495, 169 BOLD:AAB7105) were used for model fitting and five projected. ON_14020 is the only male included in the genotyping panel, where about a quarter of its private-allele states match the other species.

Application of N-CPLS discriminant analysis to the whitened chemical data of all male wasps for which all three covariates were recorded (N = 205; 36 of BOLD:AAA9495, 169 of BOLD:AAB7105), clearly separated wasps of the two barcode groups along the first latent variable (Fig. 1B), with no relevant outliers. Nested cross-validation consistently selected one latent variable in all outer folds. The mean class-balanced cross-validated accuracy was 100.0%, and no permutation run reached this value (1,000 permutations; *p* < 0.001). The outlier scan based on a model with one latent variable identified no sample with a misassignment ratio greater than 50% (Supplementary File 3). Projection of an additional five samples with known barcode identity (all BOLD:AAA9495) placed all samples within the score space of the corresponding barcode group.

In each model, the two highest discriminant coefficients, favouring BOLD:AAA9495, and the two lowest, favouring BOLD:AAB7105, each lay at the retention index of a peak (Fig. S2, Fig. S3; bold in Tab. S3). In females, the highest were those of 7-MeC27 and the 5,15-;5,17-diMeC35 peak and the lowest those of the alkenes C27:1 and C29:1; in males, the highest were those of 10,14-diMeC30 and 9,17-diMeC35 and the lowest those of C31:1 and 11,15-diMeC33.

Scored against a detection limit, none of the eight substances at the extremes of the discriminant coefficient profiles were both confined to one species and detected in all of its individuals (Tab. S8). In females, the alkenes C27:1 (RI 2691.5) and C29:1 (RI 2893.2) came closest: at least one of them was detected in 368 of 475 *P. pectoralis* but in only 1 of 166 *P. dominula* females. Both are minor peaks and can become undetectable in low-concentration extracts. 7-MeC27 and the 5,15-;5,17-diMeC35 peak, by contrast, were more abundant in *P. dominula* females but also detected in many *P. pectoralis* females. In males, each of the four substances was largely confined to one species, detected in at most one individual of the other, and 10,14-diMeC30 came closest to a species-specific substance, detected in all 41 *P. dominula* and in 1 of 169 *P. pectoralis* males.

Where single substances fell short, the relative height of two neighbouring peaks separated the species in both sexes, without a fitted model or peak integration. In female *P. pectoralis*, the 13-;11-MeC27 peak was typically much higher than the 7-MeC27 peak (median TICC signal ratio 8.62), whereas in *P. dominula* the 7-MeC27 peak was typically about three times as high (median 0.33; Fig. S8). The ratio separated the species with an AUC of 0.966, and runs with a ratio above 1.67 were predicted to be *P. pectoralis* with a LOOCV balanced accuracy of 95.2% (Fig. S9). In male *P. pectoralis*, the 11,15-diMeC33 peak was typically 37% higher than the 5,15-diMeC33 peak (median ratio 1.37), whereas in *P. dominula* it reached only about a tenth of its height (median 0.11; Fig. S10). The ratio separated the males with an AUC of 0.999, and with a threshold of 0.22 the LOOCV balanced accuracy was 99.7% (Fig. S11).

### Genomic Pool-Seq

We detected significant genome-wide differences in SNP allele frequencies between the two BINs in the Freiburg population. Of the 2,164,621 detected SNPs, 636,391 were significant at a false discovery rate of 1% (*p* < 0.003) and 215,155 remained significant under the stricter Bonferroni-corrected threshold (*p* < 2.31e-08). We established a panel of 18,026 private allele SNPs for individual genotyping (visualised in Fig. S12). The sliding window F_st_ analysis shows a median fixation index of 0.146 across all included contigs. Per-scaffold median F_st_ values indicate high genome-wide genetic divergence in sympatry (Fig. S13).

### Individual Sequencing

All individuals subjected to genotyping with the private allele panel showed strong association with either Pool A (mt-haplotype BOLD:AAA9495) or B (mt-haplotype BOLD:AAB7105). Among the twelve genotyped individuals, ten were classified as homozygous for more than 95% of allele states, while two showed considerable proportions of nuclear genetic admixture (Fig. 2). Of the latter two, the male (ON_14020) showed a roughly 3/4 to 1/4 split between Pool allele states, while the female (ON_13593) was roughly 3/4 homozygous for Pool-A-allele-states and 1/4 heterozygous for allele states in both Pools. For two individuals, we detected a mismatch between genotype and COI barcode assignment. For the female ON_13593, the distribution of allele states across the genome showed a concentration of heterozygous sites on three scaffolds (Fig. S14), consistent with recent hybridisation. The possible cross and backcross histories compatible with the observed genotype place the single hybridisation three to five generations ago. Similarly, in ON_14020 the likely-introgressed genetic information was also concentrated in few scaffolds that can be explained by a single hybridisation event three to four generations ago (Fig. S15). The two assemblies previously published as *P. dominula* [62,63] are both allied with *P. pectoralis*: at 90.0% and 89.7% of the panel loci recovered by whole-genome alignment they match Pool B, and they agree with each other at 98.2% of shared loci (Fig. S16, Tab. S9, Tab. S10; see Supplementary Results).

**Fig. 2:**
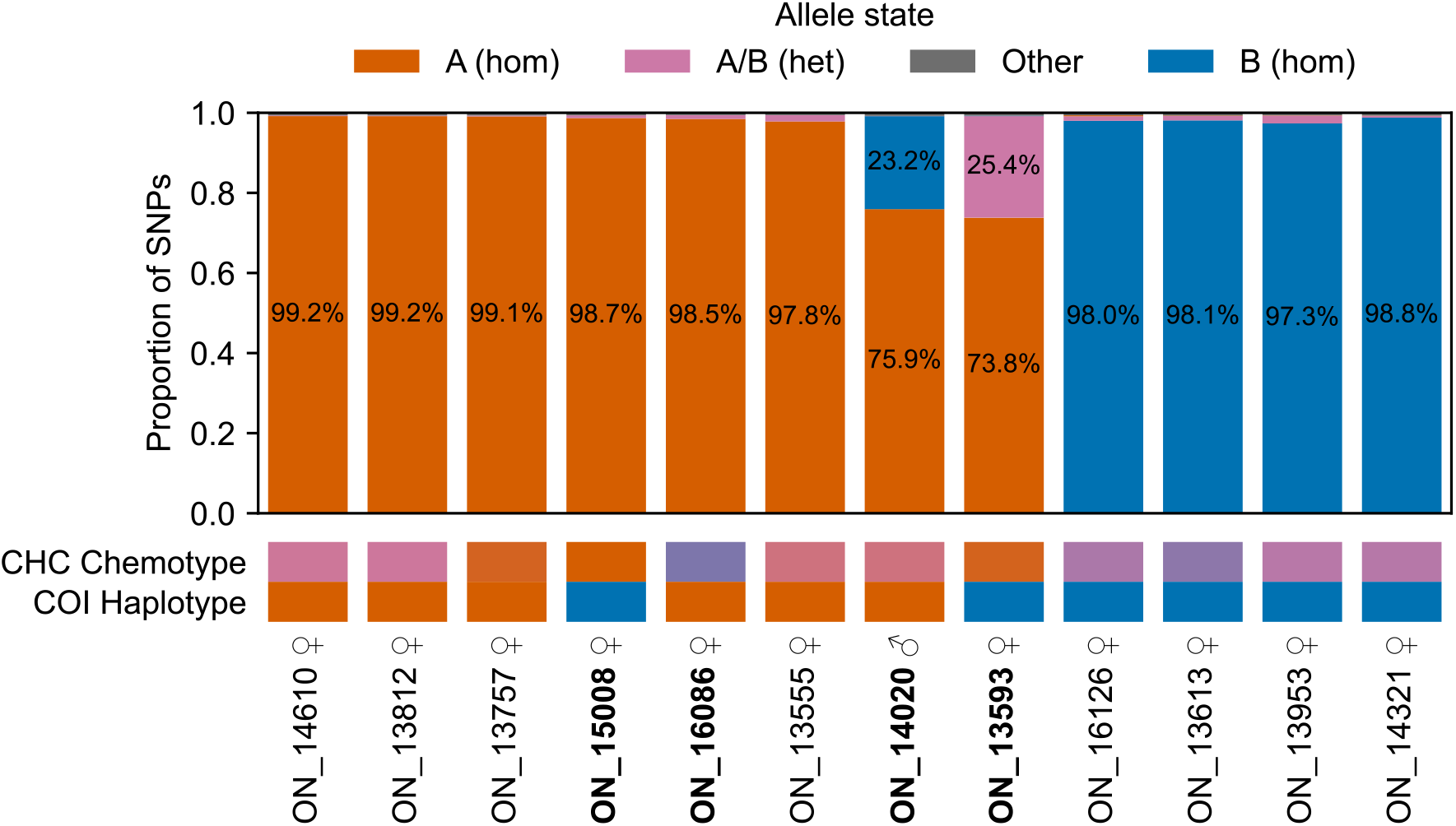
Proportions of allele states in selected individuals for a panel of 18,026 SNPs. Selected individuals showed a misassignment ratio >50% in the outlier scan of the chemical analysis. Allele counts are summarised as matching Pool A (orange), matching Pool B (blue) or matching a heterozygous state with both alleles present (magenta). Samples are ordered left to right by decreasing A-allele-state percentage. For chemotype colour representation, latent variables LV1 and LV2 from the N-CPLS analysis were projected onto the axis connecting the two group centroids (the mean score positions of BOLD:AAA9495 and BOLD:AAB7105), reducing each specimen to one value on the A-to-B chemical gradient. Colour was then mapped along this gradient from orange (A) through magenta (ambiguous) to blue (B), with the ends of the scale anchored at the median projected score of each barcode group; specimens beyond the medians were saturated to orange or blue, respectively. The mitochondrial BIN membership is shown in orange for BOLD:AAA9495 and in blue for BOLD:AAB7105. Bold sample IDs highlight individuals with noteworthy genotype-phenotype discrepancies.

### Wing Morphometrics

The geometric morphometric analysis of forewing shape revealed subtle but consistent differences between the two BIN groups. The first two principal components explained 32% of the shape variation, with no strong clustering by BIN identity (Fig. S17). Although the CVA revealed a separation between the two groups, there was notable overlap in the distributions (Fig. S18). Permutation tests indicated that between-group wing shape variance exceeded what would be expected from within-group variance alone (Goodall’s *F*: 10.01, *p* < 0.001). Approximately 63% of the multivariate shape variance was attributable to group differences (Pillai’s Trace *V* : 0.63, *p* < 0.001). Leave-One-Out Cross-Validation (LOOCV) yielded 82.0% predictive accuracy; the confusion matrix and per-group accuracies are shown in Tab. S11. Although the covariance-scaled Mahalanobis distance was large (2.62), the pure Procrustes shape distance was small (0.0081). These differences, though statistically significant, are too subtle for unambiguous individual assignment based on wing shape alone.

### 3D Somatic Morphometrics

The 3D morphometric analysis of somatic characters showed pronounced differences between the BIN groups, exceeding those in wing shape. PCA revealed a clear tendency toward separation along the first principal component, with some overlap between groups (Fig. 3A). The suspected syntypes showed no strong association with either BIN. One individual (ON_15008), assigned to BOLD:AAB7105 by COI barcoding, clustered with BOLD:AAA9495 in the PCA, consistent with the chemical profile analysis and individual sequencing and pointing to mitochondrial introgression. LDA ratio extraction identified the pronotal-collar-breadth to toruli-distance ratio as the most discriminating variable (AUC = 0.99). Plotting the two most discriminating ratios revealed clear separation between the two species with minimal overlap (Fig. 3B). Syntype ON_15401 clustered strongly with *P. pectoralis*, though no chemical profile data are available for it. Syntype ON_15402, by contrast, clustered with *P. dominula*, consistent with its chemical profile (compare Fig. 1A). The ROC curve analysis for the four most discriminating ratios is shown in Fig. S19; sensitivity and specificity are reported in Tab. S12.

**Fig. 3:**
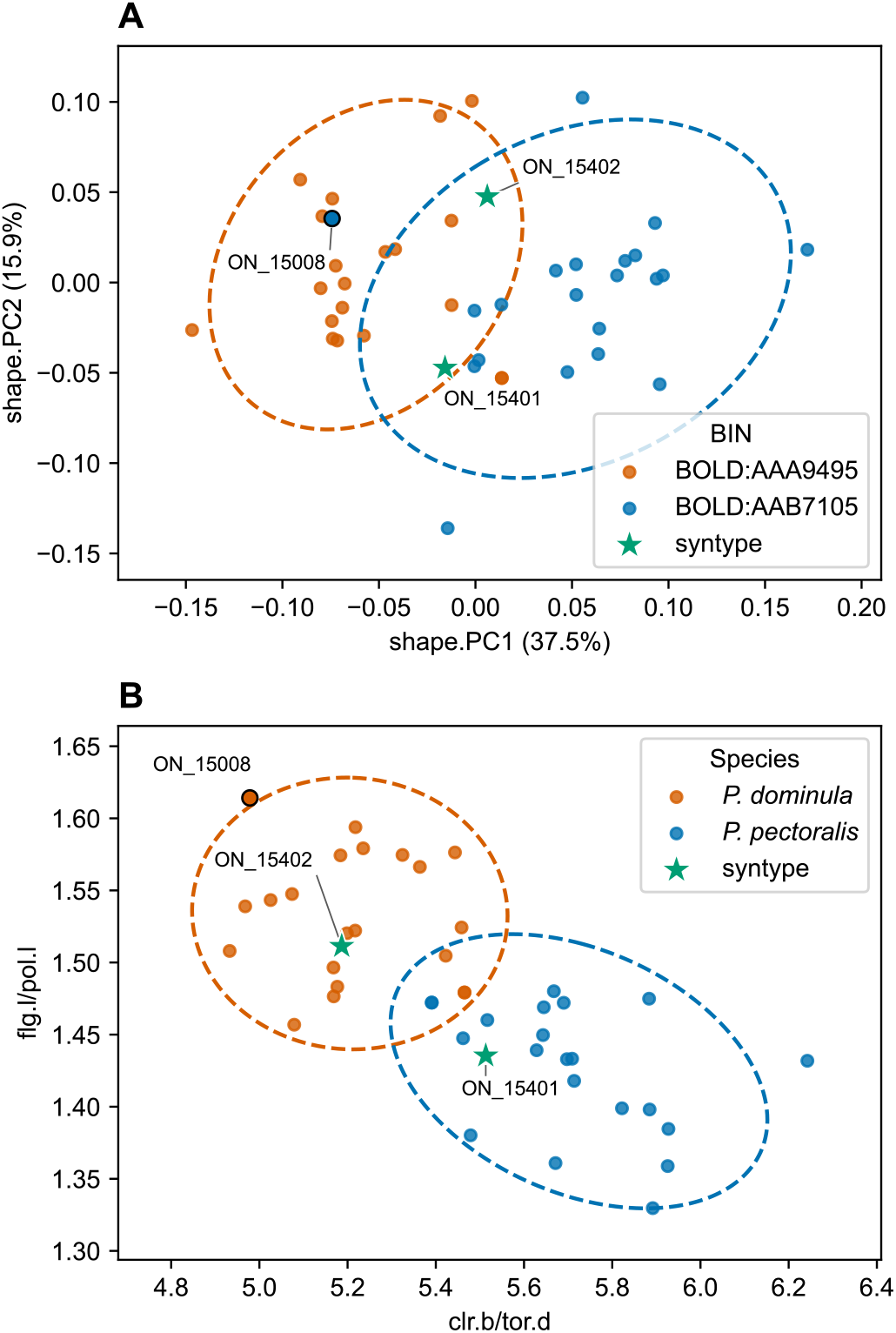
3D somatic morphometrics of the *P. dominula* species complex with each wasp represented by one point. **A:** PCA, coloured by BIN (BOLD:AAA9495 in orange, BOLD:AAB7105 in blue). **B:** The two most discriminating ratios from the LDA ratio extraction, pronotal collar breadth to toruli distance (x) and length of the first flagellomere to the distance between the lateral ocelli (y), coloured by species assignment (*P. dominula* in orange, *P. pectoralis* in blue). The syntypes (green stars) are labelled. ON_15008, outlined, is in the individual genotyping panel (Fig. 2). ON_15008 carries a BOLD:AAB7105 haplotype but is assigned to *P. dominula* on its nuclear genotype (Fig. 2) and chemical profile, pointing to mitochondrial introgression, so it changes colour between the panels.

### *Wolbachia* Multi Locus Sequence Typing

The MLST analysis revealed differences in prevalence and strain identity between the two BIN groups. BOLD:AAA9495 showed a lower *Wolbachia* infection rate of 28% and higher strain diversity (MLST sequence types ST-37, 355, 500). BOLD:AAB7105 showed a higher infection rate of 59% with slightly lower strain diversity (ST-37, 83) (Fig. S20). The strain most common in BOLD:AAB7105 was the only strain shared between the two groups.

### Taxonomic Designations

**Disclaimer:** This preprint is not issued for the purposes of zoological nomenclature (ICZN Art. 8.2), and as a preliminary version it is not a published work (Art. 9.9). The lectotype and neotype designations it describes take effect only with the publication of the peer-reviewed article.

### Polistes dominula (Christ, 1791)

**Lectotype (here designated):** ♀, Gerning Collection, Museum Wiesbaden, Germany, leg. J. I. Gerning before 1791, det. S. Jung; specimen ON_15402 (Tab. S13; Fig. S5C/D).

Christ’s description of *Vespa dominula* designated no holotype [32], and the two female *Polistes* specimens in the Gerning Collection that correspond to this taxon are likely its surviving syntypes. They likely belong to different species of the complex. ON_15402 is referred to the species of BOLD:AAA9495 by both lines of evidence (Tab. S14): its cuticular hydrocarbon profile lacks the alkenes of *P. pectoralis*, has a 13-;11-MeC27 to 7-MeC27 TICC signal ratio of 0.19 (Fig. S8, Fig. S9) and is placed with BOLD:AAA9495 by the N-CPLS model (Fig. 1A, projected), and both of its somatic ratios (**clr.b/tor.d** 5.2, **flg.l/pol.l** 1.51; Fig. 3) lie outside the range of *P. pectoralis*. The somatic morphology of ON_15401, for which no cuticular hydrocarbon profile is available, is consistent with *P. pectoralis*. As long as both remain syntypes, the name *P. dominula*, hitherto applied to both members of the complex, could be fixed to either species. We therefore designate ON_15402 as lectotype, the syntype whose identity rests on both chemical and morphological evidence. The specimen is recognisable from the data and surface renderings in Tab. S13 and Fig. S5. ON_15401 thereby becomes a paralectotype (Art. 74.1.3 of the International Code of Zoological Nomenclature [95]), a status that follows from its membership in the type series, not from its identity.

### Diagnosis

Separated from *P. pectoralis* by the cuticular hydrocarbon profiles in both sexes and, in females, by two proportions of the head and pronotum, measured as in Tab. S5 and Fig. S6 and shown on the neotype of *P. pectoralis* in Fig. S21. The COI barcode is usually BOLD:AAA9495. How these characters combine to identify a specimen, and how often each state was found in the material examined, is given in Tab. S14.

Females carry relatively more 7-methylheptacosane (7-MeC27) than those of *P. pectoralis*: their 13-;11-MeC27 to 7-MeC27 TICC signal ratio (RI 2730.1 and 2738.1; Fig. S8, Fig. S9) is typically 1.67 or below. They typically lack the two alkenes of *P. pectoralis* females, heptacosene (C27:1) and nonacosene (C29:1) (Tab. S8). A ratio of pronotal collar breadth to the distance between the antennal toruli below 5.4 (**clr.b/tor.d** < 5.4), or of the length of the first flagellomere to the distance between the lateral ocelli above 1.48 (**flg.l/pol.l** > 1.48), points to *P. dominula*. The species overlap at **clr.b/tor.d** 5.4–5.5 and **flg.l/pol.l** 1.46–1.48.

Males carry 10,14-dimethyltriacontane (10,14-diMeC30, RI 3056.7; Tab. S8) and relatively more 5,15-dimethyltritriacontane (5,15-diMeC33): their 11,15-diMeC33 to 5,15-diMeC33 TICC signal ratio (RI 3351.5 and 3373.2; Fig. S10, Fig. S11) is 0.22 or below.

*Polistes pectoralis* Herrich-Schäffer, 1841 (rev. stat.)

**Neotype (here designated):** ♀, Germany, Freiburg, Eschholzpark, 48.00° N, 7.83° E, 2022-0608, leg. et det. M. Lehmann; COI BIN BOLD:AAB7105; deposited with the ID ZFMK-TIS-89039 in the Leibniz Institute for the Analysis of Biodiversity Change, Bonn (LIB) (Tab. S13).

*P. pectoralis* was described from Italian material [96] and has since been treated as a junior synonym of *P. dominula*. We revive it for the taxon corresponding to BOLD:AAB7105 and designate a neotype with the express purpose of clarifying the taxonomic status of the name. The name must now be tied to one of two morphologically similar species, and its original name-bearing type, a male, is lost [12]. The taxon is separated from *P. dominula* by its cuticular hydrocarbon profile, its somatic proportions and, with exceptions due to mitochondrial introgression, its COI barcode (see Diagnosis below and Tab. S14). Both lines of evidence identify the neotype as *P. pectoralis*: its cuticular hydrocarbon profile contains C27:1 and C29:1 and has a 13-;11-MeC27 to 7-MeC27 TICC signal ratio of 2.42 (Fig. S8, Fig. S9), and both of its somatic ratios (**clr.b/tor.d** 5.8, **flg.l/pol.l** 1.40; Fig. 3) point to *P. pectoralis*. The specimen is recognisable from the data in Tab. S13, from the photographs in Fig. S21 and from its COI sequence, and is the property of the LIB, a research collection that preserves name-bearing types and makes them accessible for study.

The neotype comes from Freiburg, whereas the original type came from Italy. No material from that region was available to us for collecting the cuticular, morphometric and barcode data by which the two species can be separated. The taxon is continuously distributed across Europe. Freiburg, the closest of our sampling areas to Italy, is therefore as near the original type locality as was practicable. Under Art. 76.3 of the Code [95] the type locality of *P. pectoralis* becomes that of the neotype.

### Diagnosis

Separated from *P. dominula* by the characters given for that species. The COI barcode is usually BOLD:AAB7105.

Females have a 13-;11-MeC27 to 7-MeC27 TICC signal ratio above 1.67 (Fig. S8, Fig. S9), and most carry heptacosene (C27:1, RI 2691.5), nonacosene (C29:1, RI 2893.2) or both (Tab. S8). Both are minor peaks and can become undetectable in low-concentration extracts. A **clr.b/tor.d** above 5.5 or a **flg.l/pol.l** below 1.46 points to *P. pectoralis*.

Males typically lack 10,14-diMeC30 and have an 11,15-diMeC33 to 5,15-diMeC33 TICC signal ratio above 0.22 (Fig. S10, Fig. S11).

## Discussion

The concordant morphological, chemical, and nuclear genetic differences we found between the two sympatrically occurring mitochondrial haplotypes of the *Polistes dominula* complex indicate a strong, even though still incomplete, reproductive barrier. Except for wing morphology, species-specific trait differences are immediately apparent in visualisations of overall variance and become diagnostic when discriminant procedures are applied. Chemical profiles were diagnostic, with mean class-balanced cross-validated accuracies of 99.1% for females and 100.0% for males. Even individuals with the highest chemical misclassification rates are unambiguously assigned by the private allele panel. Consistent application of the biological species concept requires considering these groups as two species. We did not perform crossing experiments, and our inference therefore rests on the pattern of concordance observed in nature. This is, however, what the concept asks for, as it defines species as groups of interbreeding natural populations that are reproductively isolated from other such groups [97]. In our sampling area the two groups occur syntopically, yet remain separated by fixed differences at more than 18,000 nuclear loci and by near-diagnostic chemical profiles, with only three individuals carrying evidence of hybridisation or introgression. Crossing experiments could nevertheless resolve a distinction our data cannot yet make, namely whether prezygotic or postzygotic barriers maintain that isolation.

Examining individuals for whom prediction of the mitochondrial haplotype from their chemical profiles was difficult reveals a complex pattern of genotype associations. The two individuals with the highest chemical misclassification rates, ON_13593 and ON_15008, both show a complete mismatch between chemical profile and mitochondrial haplotype, with their chemotype and nuclear genotype agreeing against the mitochondrial genotype. In ON_15008, the nuclear genome carries little or no introgression, so the discordance is essentially confined to the mitochondrial genome, the signature of mitochondrial introgression, in which the BOLD:AAB7105 mitochondrial genome has passed into a *P. dominula* nuclear background. ON_13593, by contrast, carries considerable nuclear introgression, with heterozygous blocks concentrated on three scaffolds (Fig. S14), yet its chemical profile still falls squarely on one side rather than appearing intermediate. That a genome with such a degree of admixture still yields an unambiguous chemotype suggests that the chemical phenotype is governed by relatively few loci, so that the profile of one species can shift to that of the other through a small number of genomic changes. In both individuals carrying nuclear admixture, ON_13593 and ON_14020, the introgressed material is confined to a few scaffolds rather than dispersed across the genome (Fig. S14, Fig. S15). This blockwise distribution is, however, what recombination alone predicts given the recent backcross histories we infer, so it does not by itself localise the loci that govern the chemical phenotype. Localising them would require scoring introgressed blocks and chemotypes jointly across a mapping population of hybrids. Individual ON_16086 is an even more extreme instance of the same effect. Both the SNP panel and the mitochondrial haplotype unambiguously assign it to BOLD:AAA9495, yet despite no notable introgressed blocks (<1% AB heterozygous alleles, <0.05% B homozygous alleles) its chemotype matches *P. pectoralis*. Together these cases show that COI alone can misassign individuals in this species complex, although such cases were rare in our sample. Its reliability elsewhere is unknown as introgression rates may be higher in other contact zones, and comparative mitochondrial and nuclear data from such zones are currently lacking.

An alternative explanation for the chemotype–genotype mismatch in ON_16086 lies in the complex network of obligate and facultative interspecific usurpation observed in *Polistes* colony founding. Usurpation by obligate socioparasites such as *Polistes sulcifer* is accompanied by a shift in the usurper’s chemical profile toward that of the host, most likely through acquisition of host CHCs through interindividual contact or exchange with the nest surface [26]. Similar processes likely occur in facultative usurpation, as observed in *P. nimpha* [30]. Individual ON_16086 may simply be a potential *P. dominula* usurper congregating in a *P. pectoralis* nest, having thereby acquired the *P. pectoralis* chemotype. Most genotyping efforts in females with ambiguous chemotypes revealed no signal of hybridisation. Chemical misassignment therefore need not indicate introgression. Beyond acquisition from the social environment, the cuticular hydrocarbon layer is inherently multifunctional. Compounds that serve as semiochemicals also form the insect’s primary barrier against desiccation, so profile composition responds to ambient conditions [98] and to the physiological state of the individual as well as to genotype [99]. We conditioned the discriminant models on sampling date for this reason. Reproductive status, by contrast, was not recorded, although both warrant careful attention wherever CHC profiles are used for species inference. In our own data the two sources of variation are separable. The principal component carrying the taxonomic contrast (PC3) shows no detectable association with day of year (*r* = −0.04), whereas the seasonal signal loads on a different component (PC2, *r* = −0.51). The sequenced male (ON_14020), however, exhibits clear genetic admixture between *P. dominula* and *P. pectoralis*. Interestingly, this genetic introgression appears to be qualitatively mirrored in its chemical profile. Along the primary discriminating axis, its chemical signature is shifted away from the *P. dominula* centroid toward *P. pectoralis* to a degree that reflects its mixed ancestry. While discriminant projections must be interpreted with caution, this apparent alignment between genomic and chemical intermediacy suggests hybrid males could serve as a valuable resource for mapping the loci and genomic changes underlying chemical profile variability.

Although this study was not designed to detect nuclear genetic introgression or recent gene flow, three findings support splitting *P. dominula* sensu lato into two distinct species. First, chemotype–haplotype mismatches are detectable but rare in our sample (eight individuals with misassignment ratios above 50% among 642 analysed female wasps), even though the two barcode groups co-occur in sympatry, indicating that hybridisation is either very infrequent or that hybrids have very low fitness. Second, we found two individuals whose genotype is consistent with independent recent hybridisation events inferred to lie three to five generations back. Third, females mate with males in lek aggregations where males deposit cuticular hydrocarbons onto the substrate [27,28]. Whether females discriminate among males on the basis of these compounds has not been demonstrated, but the pronounced differences between male chemotypes would make chemically mediated assortative mating a plausible contributor to the reproductive barrier. Furthermore, although the female N-CPLS-DA model was highly accurate overall (99.1% mean class-balanced cross-validated accuracy), eight females were nonetheless misassigned in more than half of the hold-out tests in which they appeared. The male model, by contrast, reached 100.0% accuracy, and no male showed a comparable misassignment ratio. This is consistent with stronger incompatibility effects in the haploid sex, whether of nuclear, mitonuclear, or cytoplasmic origin. An elevated lethality of F_2_ males alone could lead to an absence of intermediate chemical phenotypes in the population. Additionally, male chemical profiles can potentially sharply diverge in the presence of reinforcement selection [100], as the expression of male chemical phenotypes is not held back by heterozygosity. The observation of unidirectional mitochondrial introgression from *P. pectoralis* into *P. dominula*, despite both species sharing the same *Wolbachia* strain at vastly different prevalences, can be explained by genetic hitchhiking. When infected hybrid females carrying the BOLD:AAB7105 mitochondrial haplotype backcross into the largely uninfected *P. dominula* population, they possess a distinct reproductive advantage: they can successfully mate with both infected and uninfected males. Because the bacteria and mitochondria are maternally co-inherited, the BOLD:AAB7105 haplotype might have been able to penetrate into the *P. dominula* gene pool by simply hitchhiking on the expanding *Wolbachia* infection. The divergent *Wolbachia* strains and infection rates of the two species point to differences in reproductive biology and are themselves likely to contribute to the reproductive barrier between them, since cytoplasmic incompatibility reduces the fitness of crosses between differently infected individuals.

Our designation of the syntype ON_15402, for which both chemical and morphological data are available, as lectotype of *P. dominula* should anchor a stable taxonomy for the species complex. The glacial refugial origin hypothesis for the two species remains viable and the geographic origin of *P. pectoralis* may be the Italian Peninsula. The oldest valid synonymised name for *Polistes dominula* is *Polistes pectoralis* Herrich-Schäffer, 1841 [96], which is coincidentally also based on material of Italian origin, but its male type specimen has been declared lost [12]. The species taxon broadly corresponding to BOLD:AAB7105 should therefore be named *Polistes pectoralis*. This has the unfortunate consequence that most literature on the invasive characteristics of *P. dominula* would need to be reinterpreted as describing the invasive characteristics of *P. pectoralis*, since the latter species appears in barcoding data outside its native range (see Supplementary Results) and is represented by the previously published reference genome assemblies from the United States and New Zealand. The clear, but incomplete association of these two assemblies with *Polistes pectoralis*, when only considering private alleles detected in one German population, suggests that more extensive sampling in Europe is needed for both species to detect truly species-specific SNP alleles and the large overlap in SNP allele states hints at a historical connection between the invasions of the United States and New Zealand by *P. pectoralis*.

With the integrative methods we applied and evaluated, it should be possible to identify both species with reasonable confidence in laboratory settings from live or museal specimens. As even rare hybridisation events between *P. dominula* and *P. pectoralis* lead to mitochondrial introgression and therefore COI-based misidentification, this study underscores the need for cautious use of mitochondrial barcoding in species identification and delimitation, and supports the adoption of nuclear markers in taxonomic studies [101]. A specimen whose lines of evidence conflict (Tab. S14) may be of mixed ancestry and needs such markers, for instance the private allele panel used here (Fig. 2), to be identified. While standardised tools for routine chemical identification of insect species are not yet available, Gas Chromatography–Mass Spectrometry (GC–MS) analysis of CHC extracts is relatively quick and inexpensive and can also be applied to museum specimens [102]. In our study, CHC profile analysis distinguished *P. dominula* from *P. pectoralis* with very high accuracy. However, it should ideally be complemented by COI barcoding within an integrative identification framework. That accuracy rests on the whole profile and a fitted model. Simpler chemical characters (see Diagnosis) leave a minority of specimens undetermined but can be detected from a single chromatogram without detailed quantitative analyses. The specific alkenes that are mostly present in *P. pectoralis* but lacking in *P. dominula* are minor compounds and can become undetectable in dilute extracts. Because chemical differences may be more pronounced in sympatry as a result of reinforcement, we cannot rule out the possibility that some of the observed differences are specific to the contact zones investigated in our study. Morphological data, on the other hand, appeared less diagnostic for the two cryptic species. While wing venation patterns are successfully used in honey bees to distinguish subspecies [83] and even different populations [103], it performs with only about 80% accuracy in the *Polistes dominula* species complex. The use of somatic characters for species identification seems more promising, but whether the precision of SR-µCT-based imaging is needed, or the measurement of the discriminating ratios can be achieved with a light microscope, remains to be tested. As the suggested classification thresholds are based on 20 females per species, values close to the classification threshold should be used with caution.

Our chosen approach is best applied to taxonomic questions in sympatry. The third prominent mitochondrial haplotype in the *Polistes dominula* species complex, BOLD:ACR3974, was recorded in Morocco, as well as the isles of Sardinia and Corsica. It is currently unknown whether this haplotype’s distribution also overlaps with the two species studied here, which could open the door to more discoveries of diversity and taxonomic complexity in what was known as *Polistes dominula*.

## Acknowledgments

Parts of this work were funded by the Deutsche Forschungsgemeinschaft (DFG, German Research Foundation) within the Priority Programme SPP 2349 (‘Genomic Basis of Evolutionary Innovations,’ GEvol; project number 458908270). In particular, the project ‘Genomic and Gene Regulatory Basis of Rapid Evolutionary Diversification of a Multifunctional Trait’ (project number 503307636) provided support to BF, FM, JB, ON, TS, and VN. We thank Wolfgang Haberer, Elvira Hafner, and Bettina Dobrescu for technical support. We thank Swane Jung for the discovery of the *P. dominula* syntypes in the Gerning collection and Fritz Geller-Grimm for providing access to them. We thank Theo Peeters, Jan Smit and Wim Klein for providing *P. dominula* samples from various localities in the Netherlands and Gerd Reder for providing us a sample of *P. pectoralis* from Rhineland Palatinate. We thank Fabian Staubach and Jonathan Henshaw for the helpful methodological discussions. We thank Angelica Cecilia for assistance at the IMAGE beamline and Tomáš Faragó for tomographic reconstruction of the SR-µCT data. Furthermore, we want to thank Maike Faller and Thomas Pauli for support in combined sampling efforts. The authors acknowledge support by the High Performance and Cloud Computing Group at the Zentrum für Datenverarbeitung of the University of Tübingen, the bwForCluster ‘Helix’ and ‘BinAC2’ initiatives, the state of Baden-Württemberg through bwHPC, and the German Research Foundation (DFG) through grants no INST 37/935-1 FUGG and INST 35/1597-1 FUGG. We acknowledge the KIT Light Source for provision of instruments at their beamlines and we would like to thank the Institute for Beam Physics and Technology (IBPT) for the operation of the storage ring, the Karlsruhe Research Accelerator (KARA).

## AI Use Declaration

Claude (Anthropic; Claude Opus 4.8 to 5.5 and Sonnet 5.5, through Claude Code) was used to edit and shorten the text, to check it against the data, and to write and revise code to render figures, tables and manuscript documents, as well as selected analysis steps. ChatGPT (OpenAI; various models through Codex) was used to write and revise code for JuChrom.jl and NCPLS.jl. The authors reviewed all AI-assisted text and code and take full responsibility for the content.

## Author Contribution Statement

F.S.: conceptualization, data curation, formal analysis, investigation, methodology, resources, software, validation, visualization, writing—original draft; K.J.: conceptualization, methodology, validation; M.J.B.L.: formal analysis, investigation, methodology, visualization, writing—original draft; L.M.: formal analysis, investigation, visualization, writing—original draft; H.M.O.: formal analysis, investigation; L.D.: software, validation; A.K., L.B., J.F.: investigation, validation; T.v.d.K., M.Z.: investigation, resources, visualization; A.M.B., A.B., F.B., M.E.S., S.T., E.v.d.S., J.J.W.: investigation; L.A.W.: visualization; J.B., F.M., T.S.: funding acquisition, investigation, project administration; B.F.: funding acquisition, methodology, project administration, resources, validation; J.R.: investigation, supervision; V.N.: funding acquisition, methodology, project administration, supervision, validation; O.N.: conceptualization, data curation, formal analysis, funding acquisition, investigation, methodology, project administration, resources, software, supervision, validation, visualization, writing—original draft; all authors: writing—review and editing.

## Conflict of Interest Declaration

We declare we have no competing interests.

## Data Accessibility

On publication, a single FreiData record will link to every item below and become the entry point to cite.

**Genome assembly.** The near chromosome-level *P. dominula* assembly is available through ENA under accession GCA_965197175.2.

**Raw sequence data.** Pool-Seq and individual-genome raw reads are deposited at ENA under BioProject PRJEB115294 (secondary accession ERP195513) and will be released on publication.

**Barcode data.** COI traces and barcode sequences, including that of the neotype of *P. pectoralis*, are deposited in BOLD and will be made public on publication.

**Morphological data.** Wing images, µCT scan data and the full-resolution photographs of the neotype of *P. pectoralis* (Fig. S21) will be made available at FreiData on publication.

**Chemical data.** Raw GC–MS chromatograms will be made available at FreiData on publication.

**Software and workflows.** Chromatogram processing and the N-CPLS implementation are at https://github.com/oniehuis/JuChrom.jl and https://github.com/oniehuis/NCPLS.jl. The Pool-Seq workflow is at https://github.com/git-hobo/popoolation2/tree/master/workflow, with its analysis container as docker::fabianschweitzer/popoolation2:v1.0.0. The SNP liftover and individual genotyping workflows are at https://github.com/git-hobo/pdom_snp_allele_states (snp_panel_liftover.smk and snp_panel_typing.smk) and will be released on publication. Hi-C scaffolding used the Arima mapping pipeline (https://github.com/arimagenomics/mapping_pipeline).

**Supplementary material.** Supplementary Files 1–4 — sample collection data, the female and male outlier-scan tables, and the external BOLD records — are available online, together with the Supplementary Methods, Figures and Tables.

## Supplementary Materials

### Supplementary Methods

#### Gas Chromatography–Mass Spectrometry

We extracted cuticular chemical profiles from the sampled wasps of the *P. dominula* complex and from both syntypes of *P. dominula*, with the exception of two males (ON_15380 and ON_15382) that were reserved for long-read genome sequencing and Hi-C scaffolding and therefore had to remain intact. Extraction yielded profiles suitable for analysis for 852 individuals. Wasps were extracted by submerging them in *n*-hexane at room temperature for 10 minutes. Extractions were carried out in 2 mL screw-top vials (Agilent Technologies), with the solvent volume adjusted to specimen size so that each wasp was covered to the wing tips. The resulting differences in extract concentration are addressed by the compositional treatment of the chromatograms described below. The two syntypes were instead extracted for 30 minutes, because a 10-minute extraction yielded no usable signal for one of them and only a weak signal for the other, and we treated both identically. The longer extraction did not alter this pattern. One syntype yielded chemical data suitable for analysis, the other did not. After its initial extraction, one individual (ON_16086) was stored in absolute ethanol and extracted three further times to test whether its unexpected chemotype was reproducible. Its four extracts were analysed separately. Extracts of all individuals were analysed using a 7890B gas chromatograph coupled to a 5977B quadrupole mass spectrometer (GC–MS; Agilent Technologies, Santa Clara, California, United States), with electron ionisation at 70 eV. All extracts were injected in pulsed splitless mode into a preheated inlet (300 °C) connected to a 30 m × 250 µm × 0.25 µm HP-5ms Ultra Inert (19091S-433UI) fused silica column (Agilent Technologies), with a helium carrier gas flow of 1 mL min^−1^. The oven temperature was initially set to 50 °C, increased by 5 °C min^−1^ to a maximum of 300 °C over 50 minutes, and then held at 300 °C for 15 minutes.

#### Chemometric Analysis

To map retention times (RTs) of sample runs to Kováts retention indices (RIs) [36], we annotated standards composed of 31 *n*-alkanes, ranging from *n*-octane to *n*-octatriacontane. These standards were run on the same GC–MS instrument under identical conditions within a 12-day period of a given sample run, with a median interval of less than one day. The standard runs were processed using the same integer m/z binning rule and the same accepted m/z range, 29–562, as the sample runs, followed by dwell normalisation, assuming homogeneous dwell times. Alkane peaks were detected with JuChrom.findalkanes(), using baselinezerothreshold = 1.0 and apexmzscanorder = :descending. The annotations of the ladder steps were visually verified and then used to fit a strictly increasing, smooth cubic B-spline mapping from RT to RI with JuChrom.fitmap() using default settings. The dwell-normalised *n*-alkane standards were also used to estimate a linear intensity-variance model and the lag-1 residual autocorrelation, ρ, with JuChrom.fitalkanevariancemodel().

For each sample run, the fitted RT-to-RI mapping was first used with JuChrom.invmap() to identify the RT interval corresponding to RI 2100–3800. The m/z-binned mass scan series was then trimmed to this interval with JuChrom.retentiontrim!() and converted to a mass scan matrix with JuChrom.mscanmatrix(). Intensities were dwell-normalised with JuChrom.dwellnormalize(), assuming homogeneous dwell times.

For each dwell-normalised intensity value, an approximate variance was predicted from the fitted variance model with JuChrom.varpred(). Retention times were then converted to retention indices with JuChrom.applymap(). During this transformation, intensities were divided by the local Jacobian, *J* = dRI/dRT, which preserves peak area under the change of retention coordinate. The corresponding per-cell variances were propagated by the same fixed-scaling approximation, that is, each variance was divided by *J*^2^. We next estimated baselines for every m/z trace of each RI-transformed, m/z-binned chromatogram with JuChrom.arpls(), JuChrom’s implementation of asymmetrically reweighted penalised least-squares smoothing (arPLS) [37]. The function was called with default parameters except that baseline stiffness was increased by setting λ = 10^7^. Because the input objects were variance mass scan matrices, the previously estimated per-cell variances were used internally as fitting weights during baseline estimation. The fitted baselines were subtracted with JuChrom.subtractbaseline(). No clipping or censoring was applied at this stage. Negative baseline-corrected values were therefore retained, and the original variance matrices were carried forward without adding uncertainty from the baseline estimates. This avoided imposing an artificial intensity floor before retention binning.

To align retention coordinates among runs, we identified a common regular RI grid with JuChrom.densestgrid(), using all 855 chromatograms selected for chemometric analysis. This function searches the common retention overlap for the densest regular grid for which every run contributes at least one scan to every bin. In this dataset, this produced 1,819 RI bins with a bin width of ca. 0.9 RI units, spanning the common overlap from RI 2100.6 to 3799.7. Intensities were then mapped onto this grid with JuChrom.binretentions(), using the lag-1 residual autocorrelation, ρ, estimated from the corresponding *n*-alkane standard. For each m/z channel and RI bin, binretentions() computes an inverse-variance-weighted mean intensity from the scans falling into that bin. The corresponding variance is propagated as the variance of this weighted mean and inflated by a variance inflation factor based on ρ to account for serial correlation among adjacent scans. The resulting binned matrices contained 1,819 RI bins by 534 m/z channels.

After retention binning, non-positive and very small (see below) positive intensities were replaced with JuChrom.replacecensored() before applying the centred log-ratio transformation (CLR) [38], which requires strictly positive values. The variance-dependent detection-limit term was disabled (*k* = 0), so censoring was based only on a run-specific empirical global floor, defined as the 1% quantile of positive intensities in that run. Intensities at or below this floor were replaced by half the floor. For these censored values, variances smaller than floor^2^/12 were raised to floor^2^/12, whereas larger propagated variances were retained. The value floor^2^/12 corresponds to a uniform uncertainty model for an unknown true intensity between zero and the censoring floor.

The censored matrices were then transformed with JuChrom.clr(). This was done because absolute ion intensities were not interpreted as comparable abundances among individuals. The amount of extractable cuticular material can vary with wasp size, extraction efficiency, injection amount, and instrument response, and no biologically defensible normalisation to a common total amount was available. We therefore analysed relative chemical composition rather than absolute peak height. The CLR transformation represents each chromatogram by log-ratios among its RI × m/z features and removes arbitrary multiplicative differences in total signal between runs. The propagated CLR variances were then used for whitening with JuChrom.whiten(), using a standard-deviation floor of 10^−8^ to avoid division by nearzero variances. The resulting tensor contained chromatographic measurements for 855 chromatograms, 1,819 RI bins, and 534 m/z channels.

Before applying supervised N-CPLS discriminant modelling, we first examined the chemical data using Principal Component Analysis (PCA) as an unsupervised exploratory method to visualise the dominant patterns of variation in the chemical profiles without using Barcode Index Number (BIN) assignments or other response variables during model fitting. BIN labels were used only for post-hoc interpretation of the PCA scores, allowing us to assess whether the major axes of chemical variation were associated with BIN membership. PCA also allowed us to screen for unusual samples and broad clustering patterns before fitting the supervised N-CPLS models. PCA was performed on the centred and whitened chemical-profile matrix using the Julia package MultivariateStats version 0.10.4.

We used N-way Canonical Partial Least Squares (N-CPLS) [39] to test whether chemical profiles differed between BINs and to predict the taxonomic identities of selected samples, including a syntype, from their chemical profiles. N-CPLS was performed in Julia using NCPLS.jl version 0.0.1 [40], an in-house package written by the last author of this study and available on GitHub (https://github.com/oniehuis/NCPLS.jl). We specified a multilinear N-CPLS discriminant model and fitted it to the tensor-valued predictors, the categorical class labels, and an additional response matrix containing the geographic latitude and longitude of the sampling location, the absolute distance of the sampling date from midyear (|day of year − 183|), and a binary variable indicating whether the sampled individual was stylopised. These additional variables were encoded as floating-point values. The predictor tensor and the primary response block — that is, the internally one-hot-encoded categorical class labels — were centred. Class imbalance was accounted for by weighting samples inversely to their class frequencies. In the analysis of female samples, the predictor tensor contained 635 samples, 158 of which belonged to BOLD:AAA9495 and 477 to BOLD:AAB7105. In the analysis of male samples, the predictor tensor contained 205 samples, 36 of which belonged to BOLD:AAA9495 and 169 to BOLD:AAB7105.

Predictive performance was evaluated using the N-CPLS discriminant-analysis cross-validation utilities, with a nested cross-validation scheme as suggested by Smit et al. [41]. We applied a stratified cross-validation scheme with five disjoint outer folds and four disjoint inner folds (NCPLS.cvda()). The outer folds were used only to estimate predictive performance, whereas the inner folds were used for model selection. In each outer cross-validation split, one of the five outer folds was held out as the test set, and the remaining four outer folds were used as the outer training set. This outer training set was then further partitioned into four stratified inner folds. Candidate N-CPLS-DA models with one to four latent variables were fitted within the inner cross-validation loop, and the best-performing number of latent variables was selected for each inner validation fold. For each outer split, the final number of latent variables was defined as the median of the inner-fold selections, rounded down to an integer. With the default selector, ties were resolved in favour of the smaller component count. A final model was then fitted to the full outer training set using the selected number of latent variables and applied to the held-out outer test set.

Performance was quantified as one minus the weighted Normalised Misclassification Cost (NMC). The weighted NMC assigns each test sample an inverse-frequency class weight, normalised so that the weights sum to one, thereby reducing the influence of the more frequent BIN. The reported cross-validation accuracy therefore estimates the predictive performance of the complete modelling workflow, including latent-variable selection, rather than the performance of a model with a single fixed number of latent variables.

To assess the statistical significance of the model’s predictive accuracy, we used NCPLS.permda() to generate 1,000 null cross-validation scores by randomly permuting BIN labels across samples and repeating the same nested cross-validation workflow, including selection among one to four latent variables. The empirical one-sided p-value was calculated with NCPLS.pvalue() as (b + 1)/(B + 1), where B is the number of permutations and b is the number of permuted accuracies greater than or equal to the observed accuracy [42].

Putative outliers were screened using NCPLS.outlierscan(), which repeatedly performs stratified outer-fold testing and records how often each sample is misclassified when held out. We express this per sample as its misassignment ratio: the number of hold-out tests in which the sample was assigned to the other BIN, divided by the number of tests in which that sample was held out. The outlier scan used the same settings as NCPLS.cvda(), except that the number of latent variables was fixed at the value suggested by cross-validation: two for the female analysis and one for the male analysis. The number of outer-fold repeats was set to 1,000.

For subsequent projection and class prediction of samples not included in model fitting because they lacked mandatory metadata, we fitted final N-CPLS-DA models using the number of latent variables suggested by the nested cross-validation results. This resulted in two latent variables for the female analysis and one latent variable for the male analysis. These final models were used to project the chemical profiles of excluded samples into the corresponding N-CPLS score spaces with NCPLS.project() and to predict their BIN assignments with NCPLS.predictclasses().

Discriminant coefficient profiles were computed from the same final models. For the first *A* latent variables, NCPLS.coef() returns the regression coefficients *B* = *RQ*^T^, where *Q* holds the response loadings and *R* = *W* (*P*^T^*W*)^−1^ the weights that give the scores directly from the data (*T* = *XR*), computed from the multilinear weights *W* and the predictor loadings *P*, both unfolded to one row per retention index × m/z cell. *B* thus holds one coefficient per retention index, m/z channel and BIN response, on the scale of the whitened CLR data. With *A* = 2 for females and *A* = 1 for males, we formed the contrast *B*_AAA9495_ − *B*_AAB7105_ and summed it over the 534 m/z channels, so that all ions of a substance add up to one discriminant coefficient at its retention index.

The presence of single substances was scored in Julia version 1.12.6 [33] on each run’s chromatogram on its retention-index axis. For each substance we chose one or more detection ions specific to it within a few RI units. The molecular ion for the alkenes (C27:1, m/z 378; C29:1, m/z 406; C31:1, m/z 434) and branch fragments for the methyl-branched alkanes (7-MeC27, m/z 309; 5,15-;5,17-diMeC35, m/z 463; 10,14-diMeC30, m/z 323 and 225; 9,17-diMeC35, m/z 407 and 267; 11,15-diMeC33, m/z 351 and 239). Co-eluting internal isomers share the fragments of a common branch, so where two ions are given, one per branch, both were required. Each ion trace was smoothed over three scans and divided by the run’s total ion current, so that runs of different concentration compare. The height of a peak at a given RI was the highest local maximum within ±1.5 RI units, measured above a straight baseline drawn between the trace minima on either side within 6 RI units, as in manual integration. The detection limit of each ion was set from chance intensities. The same height was measured at every whole RI 10 to 40 units on either side of the substance, in every run. At each of these positions the 99.5th percentile over runs was taken as the largest chance intensity, and the median over positions as the limit, so that positions carrying another peak, or no signal at all, do not decide it. Because the annotated RI comes from one representative run, each substance was first re-centred on the median apex of the runs that clearly carried its first detection ion. A substance was scored as present in a run when every one of its detection ions exceeded its limit at that centre. The species labels entered none of these steps.

#### COI Barcoding

We PCR-amplified the mitochondrial cytochrome c oxidase I (COI) barcoding fragment (COI5P) using the oligonucleotide primers LCO1490-JJ and HCO2198-JJ [45; supplied by metabion international, Freising, Germany]. The PCR protocol included an initial denaturation step at 94 °C for 3 min, followed by 35 cycles of denaturation at 94.0 °C for 45 s, annealing at 54.0 °C for 30 s, and extension at 72.0 °C for 90 s, with a final extension at 72 °C for 10 min. PCR products were purified using the ExoProStar kit (GE Healthcare, Chicago, Illinois, United States) and submitted to Macrogen Europe (Amsterdam, The Netherlands) for unidirectional Sanger nucleotide sequencing. Chromatogram files were processed using Unipro UGENE version 38.1 [46]. Nucleotide sequences were quality-trimmed at both ends using a Phred score threshold of 57. Sequences shorter than 200 nucleotides were discarded and the remaining nucleotide sequences were assigned to Barcode Index Numbers (BINs) [11] using BOLDiggercline version 2.1.0 [47] for the BOLD Identification Engine API. A nucleotide sequence was assigned to a BIN if all ten of its most similar matches corresponded to that BIN, which was consistently the case.

#### Sequencing and Assembly of the *P. dominula* Genome

We collected two adult males from Schallstadt, Germany, assigned to BOLD:AAA9495 by COI barcoding (from a right hindleg, as described above), and froze them alive at −80 °C. One was used for PacBio HiFi sequencing with ultra-low-input libraries on the Sequel II platform, the other was used for assembly scaffolding with Arima Hi-C libraries and protocols. Both sequencing steps were performed by the laboratory centre of the LOEWE Centre for Translational Biodiversity Genomics (TBG) at Senckenberg. Read calling of the PacBio HiFi subreads was performed with DeepConsensus version 1.2.0 [48]. Adapter sequences from PCR amplification steps were removed with Cutadapt version 5.1 [49]. We removed mitochondrial reads by assembling the mitogenome with MitoHiFi version 3.2.1 [50] and excluding all reads involved in that assembly. The remaining reads underwent three rounds of assembly with Hifiasm version 0.25.0-r726 [51], scaffolding with the Arima mapping pipeline (available via https://github.com/arimagenomics/mapping_pipeline) and YaHS version 1.2.2 [52], and three rounds of taxonomic partitioning with the snakemake version 9.23.1 implementation of BlobToolKit version 4.5.0 [53] to remove reads mapping to contigs assigned to non-hymenopteran taxa, retaining those mapping to contigs with a hymenopteran or no taxonomic assignment. Soft-masking was performed with two rounds of RepeatMasker version 4.1.9 [54] using RepBase version 27.03 [55] and RepeatModeler2 version 2.0.6 [56]. The final assembly was assessed for gene-space completeness, contiguity, and assembly quality with BUSCO version 5.8.2 [57], compleasm version 0.2.8 [58], QUAST version 5.0.2 [59], backmap version 0.6 [60], and Merqury version 1.3 [61]. The assembly is available through ENA (NCBI GenBank Accession: GCA_965197175.2). We compared our assembly to the reference genome assemblies previously published as *P. dominula*, GCF_001465965.1 [62] and GCA_049000335.1 [63], referred to as *Pdom_r1.2(USA)* and *Pdom_raw_medaka(NZ)*, by aligning them with minimap2 version 2.30-r1287 [64] and defined a workflow for lifting over SNP coordinates of the private allele panel from the Pool-Seq data to the other reference assemblies. We identified allele states for each assembly and compared them to the allele states determined in the Pool-Seq experiment, including only loci with unique mapping results. The workflow can be found at https://github.com/git-hobo/pdom_snp_allele_states as snp_panel_liftover.smk. We also compared genetic distances between available assemblies in the genus *Polistes* by computing pairwise distances with Mash version 2.3 [65].

#### Genomic Pool-Seq

We focused on two sampling locations (Littenweiler, Freiburg and Eschholzpark, Freiburg) with roughly similar abundances of the two BINs. To ensure balanced representation, we selected an equal number of individuals per haplotype from each location, excluding those parasitised by *Xenos*. Prior to sample selection, DNA concentrations and modal fragment sizes were assessed using a fragment analyzer (Agilent 5200 Fragment Analyzer System, DNF-488 High Sensitivity Genomic DNA Kit), and samples with concentrations below 1 ng/µL were excluded. Because wasps were collected without regard to haplotype, one haplotype was typically caught in greater numbers than the other at a given site. To equalise the two, we downsampled the more common haplotype, pairing each individual of the rarer haplotype with one individual of the more common haplotype of similar modal fragment size. This yielded each of the two sampling sites contributing 27 individuals per haplotype (108 individuals total, 54 per haplotype). Individual DNA samples were then pooled in equimolar amounts to ensure equal representation of each individual in the final Pool-Seq libraries, yielding two balanced pools of 54 individuals each. The two Pool-Seq libraries were sequenced on an Illumina NextSeq™ 500 platform by StarSEQ GmbH (Mainz, Germany), with 150 bp read length and a target sequencing depth of 15 Gb per pool, yielding approximately 50 million paired-end reads per pool. Adapter trimming and quality filtering were performed using Trimmomatic version 0.39 [66]. Reads were mapped to the *Polistes dominula* reference genome (NCBI GenBank accession: GCA_965197175.2) using BWA-MEM version 0.7.18 [67]. Aligned reads with a mapping quality below 20 were excluded using SAMtools version 1.21 [68]. SNPs were identified, and genetic differentiation assessed at each SNP by Fisher’s exact test and by a sliding window computation of fixation indices (F_st_) with a window size and step width of 500, using PoPoolation2 version 1.201 [69]. Only SNPs with coverage between 40 and 200 and a minor allele count ≥ 3 (min-coverage: 40, max-coverage: 200, min-count: 3) were retained for downstream analysis. We built a Docker analysis container (available as docker::fabianschweitzer/popoolation2:v1.0.0) for it, and a Snakemake workflow (available at https://github.com/git-hobo/popoolation2/tree/master/work-flow) for use with Illumina sequencing datasets.

False Discovery Rate (FDR) correction was applied to Fisher’s exact test results using the qvalue version 2.42.0 [71] library in R version 4.3.3 [72] via RStudio version 2026.04.0+526 [73]. The FDR estimation followed the Benjamini-Hochberg procedure (λ = 0), resulting in a significance threshold of *p* < 0.003 to control FDR at 1%. We also applied a Bonferroni correction [70] for multiple testing (2,164,621 SNPs), defining a stricter threshold of *p* < 2.31e-08. Genome-wide SNP allele frequency differences were visualised using qqman version 0.1.9 [74]. We used custom Python scripts to identify SNPs that were monomorphic in one pool but completely fixed for an alternative allele in the other. These SNPs constitute the private allele panel used for individual genotyping and comparison to other reference assemblies.

#### Individual Genome Sequencing

We selected females for genome-skimming WGS based on discordance between chemotype and mitochondrial haplotype group membership. Females included in the final N-CPLS-DA model were selected, when they showed a misassignment ratio above 50% in the outlier scans. Females excluded from the final model were selected, when BIN membership was predicted incorrectly by the final N-CPLS-DA model. Males were selected when they showed a misassignment ratio other than 0%. We additionally included the only BOLD:AAB7105 female from the Netherlands (ON_16126). DNA extracts from twelve individuals were sequenced individually on an Illumina NextSeq 2000 platform by StarSEQ GmbH (Mainz, Germany) or Macrogen Europe (Amsterdam, The Netherlands), generating approximately 50 million 150 bp reads per individual. Reads were processed using the same trimming, mapping, and filtering pipeline described for Pool-Seq above. Allele states for the private allele panel were detected with the variant calling subroutine implemented in BCFtools version 1.23.1 [75]. We then examined allele states at positions corresponding to Pool-Seq private alleles and classified each as homozygous for the Pool A allele, homozygous for the Pool B allele, or heterozygous for the alleles separating the pools. All remaining variant states were assigned to another category. The workflow as implemented in Snakemake can be found at https://github.com/git-hobo/pdom_snp_allele_states as snp_panel_typing.smk. Heterozygosity was likewise visualised from the individual Variant Call Format (VCF) files. Panel sites were restricted to N90-scaffolds and ordered by scaffold and physical position. Scaffolds were concatenated end-to-end into a single genome-ordered coordinate, separated by 1-Mb gaps, to give each site a continuous plotting position. Local heterozygosity was then computed per individual as a centred sliding window over the ordered panel SNPs. The window spanned 50 SNPs rather than a fixed number of base pairs, so its physical width varies with local marker density. Within each window the rolling AB proportion is the number of sites classified as AB divided by the number of non-missing (called) sites. Windows containing fewer than ten called sites were left undefined. The resulting trace was plotted for each individual as a separate track along the genome-ordered axis (proportion 0–1).

#### Wing Morphometrics

Right forewings of 200 females (97 of BOLD:AAA9495, 103 of BOLD:AAB7105) were detached at the base using scissors, spread under alcohol (70%), and mounted between two glass slides (1 mm thickness). Wings were photographed individually using a Canon EOS RP camera mounted on an Olympus SZX16 stereomicroscope, illuminated with a Schott EasyLED Ring Light. Each wing was imaged with identical camera settings and a 1 cm scale bar, maintaining consistent orientation across samples. Image acquisition was controlled via Helicon Remote version 4.0.6W [76], and photographs were imported into Adobe Lightroom version 10.4 [77] for automatic colour correction.

To correct for lens distortion, geometric distortion was estimated from photographs of perpendicular glass slides, and a correction value of –1 was applied in Adobe Photoshop version 23.2.2 [78]. TPS files were generated with tpsUtil32 version 1.83 [79], and image order was randomised before landmark placement. Landmarks were digitised in tpsDig2 version 2.32 [80]. The TPS series Software is described by Rohlf [81].

Nineteen landmarks were defined following the protocol of Bustamante et al. [82], based on Francoy et al. [83]. As shown in Fig. S4, most landmarks were placed at vein intersections. Landmarks 5, 6, and 10 were placed at the apex or concave corner of the corresponding cells to improve consistency, following Bustamante et al. [82], who modified placement at these positions because of variable radius vein thickness.

Geometric morphometric analyses were performed in MorphoJ version 1.07a [84] using a generalised Procrustes analysis. Forewing shape variation was analysed through Principal Component Analysis (PCA) and Canonical Variate Analysis (CVA). Permutation tests with 10,000 iterations were conducted to assess the statistical significance of shape differences between BINs. A Leave-One-Out Cross-Validation (LOOCV) was then performed on the PC scores explaining 95% of cumulative variance, using R version 4.3.3 [72] and RStudio version 2026.04.0+526 [73], to evaluate the predictive accuracy of canonical variate-based discriminant modelling.

#### 3D Somatic Morphometrics

Among the 87 wasps collected specifically for morphological analysis, we selected 40 individuals for 3D morphometric analysis, aiming for 20 per BIN, based on preliminarily apparent chemical characteristics (detectability of C27:1, an alkene of *P. pectoralis* females) to avoid destructive DNA extraction before µCT scanning. BIN haplotypes were assigned after scanning was complete as described above. The two suspected syntypes of *P. dominula* were also included in the 3D morphometric analysis (Fig. S5). Selected individuals were scanned using highthroughput Synchrotron Radiation-based Micro-Computed Tomography (SR-µCT) [85] at the IMAGE Beamline [86] of the KIT Light Source. All individuals were kept in plastic vials and remained submerged in absolute alcohol during scanning. Scans were conducted with an X-ray beam diffracted by a Double Multilayer Monochromator (DMM) at a magnetic field of the wiggler of 2.7 T, yielding a maximum flux density at 16 keV with an energy bandwidth of 2%. To reduce the heat load on the DMM, the beam was pre-filtered with 3 mm pyrolytic graphite. A fast indirect detector system was used consisting of a 200 µm LuAG:Ce scintillator, a diffraction-limited optical microscope, and a 12-bit pco.dimax high-speed camera with 2,016 × 2,016 pixel resolution. An optical magnification of 2× resulted in an effective pixel size of 6.1 µm. For each scan, we took 200 dark field images, 200 flat field images, and 3,000 equiangularly spaced radiographic projections in a range of 180° with an exposure time of 6.25 milliseconds per frame, resulting in scan durations of 21 seconds. Each wasp was scanned in four to five height steps to accommodate the scanning window. Radiographic projections were cropped to the illuminated part of the field of view. concert version 0.31.0 [87] was used for automated data acquisition. Tomographic reconstruction was performed with tofu version 0.12.0 [88] to generate both phase contrast and absorption contrast datasets. Subsequently, these were blended and converted into 8-bit volumes, each comprising 2,016 × 2,016 × 700 voxels. The volumes representing the individual height steps for each specimen were registered and merged into single 3D volumes using Amira version 2022.2 [89]. The 8-bit greyscale voxel data were processed using Dragonfly 3D World version 2022.1.0 [90]. Measurements were taken three times per individual per character and then averaged. Measurements followed Wurdack et al. [91] and are defined in Tab. S5 and illustrated in Fig. S6. We used the Multivariate Ratio Analysis version 1.05 [92] scripts to compute isosize and perform a shape PCA and LDA ratio extraction with R version 4.3.3 [72] and RStudio version 2026.04.0+526 [73]. The shape PCA was coloured by BIN identity, and the LDA ratio extraction was based on and coloured by species assignment to account for mitochondrial introgression. We computed a Receiver Operating Characteristic (ROC) curve and the Area Under the Curve (AUC) metric to evaluate the performance of the four most BIN-discriminative character length ratios.

### Supplementary Results

#### Relations to the Previously Published Reference Genome Assemblies

Assembly-to-assembly alignment of the reference genomes previously published as *P. dominula* to our new assembly (GCA_965197175) recovered a majority of loci from the SNP panel. Of the 18,026 panel loci, we recovered 17,143 from the *Pdom_r1.2(USA)* assembly [62] and 17,359 from the *Pdom_raw_medaka(NZ)* assembly [63]. Both assemblies are closely allied with *P. pectoralis*, sharing 90.0% and 89.7% of private alleles, respectively (Fig. S16). As these percentages are nearly identical, we cross-checked allele states in both assemblies against our pool data (Tab. S9). The *Pdom_r1.2(USA)* and *Pdom_raw_medaka(NZ)* assemblies agree in their allele states at 98.2% of the shared 17,260 loci. This affinity with *P. pectoralis* is consistent with mitochondrial haplotype distribution data from the United States (source of the *Pdom_r1.2(USA)* assembly) and New Zealand (source of the *Pdom_raw_medaka(NZ)* assembly). Among the 43 records identified as *P. dominula* from the United States in the BOLD database [104], 42 are assigned to BOLD:AAB7105 and only 1 to BOLD:AAA9495. That singular BOLD:AAA9495 record (GenBank accession AB284742) is doubtful: the sequence is deposited under *Polistes metricus*, is documented only as originating from Ohio, and neither the database entry nor the publication in which the sequence appeared [105] records a collection date, collector, or determination. In New Zealand, both records identified as *P. dominula* are likewise assigned to BOLD:AAB7105. All 43 US and 2 New Zealand records were retrieved on 14 September 2026 and are listed in Supplementary File 4. The mash distance of our assembly to the two previously published assemblies is 6.7‰ for *Pdom_r1.2(USA)* and 6.5‰ for *Pdom_raw_medaka(NZ)*, while the latter two have a mash distance of 5.0‰ to each other. For comparison, the smallest mash distance between two recognised *Polistes* species in this set is 8.8‰ (*P. metricus* / *P. fuscatus*). The pairwise mash distances among all available *Polistes* assemblies are summarised in Tab. S10.

## Supplementary Figures

**Fig. S1:**
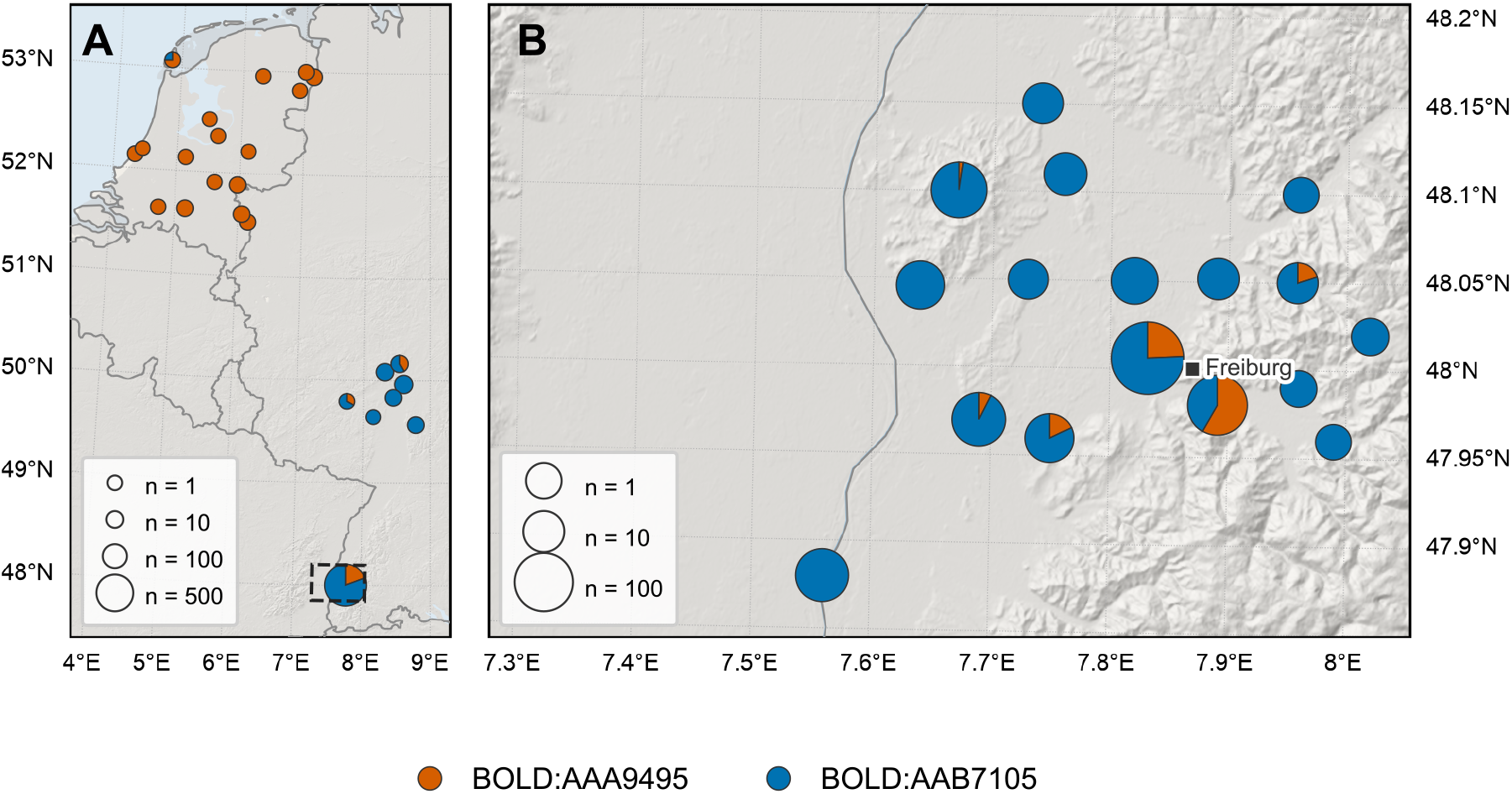
Sampling localities and their barcode composition. Specimens from the 71 localities with recorded coordinates (36 in Germany, 35 in the Netherlands) are pooled into charts by distance, within 12 km in (**A**) and 3 km in (**B**). Pie-chart radii grow with the square root of the number of specimens pooled into each, from a minimum size (see legend). (**A**): All sample sites, with the Freiburg area pooled into one chart. (**B**): The Freiburg area, outlined in (**A**), which accounts for 85% of all specimens and is drawn at its own scale. Both barcode groups were recorded together at 15 of the 71 localities. BOLD:AAA9495 in orange, BOLD:AAB7105 in blue. Relief from the Terrain Tiles on AWS Open Data: EU-DEM, produced using Copernicus data and information funded by the European Union; SRTM and GMTED2010 courtesy of the U.S. Geological Survey; ETOPO1, U.S. National Oceanic and Atmospheric Administration. Base map data from Natural Earth.

**Fig. S2:**
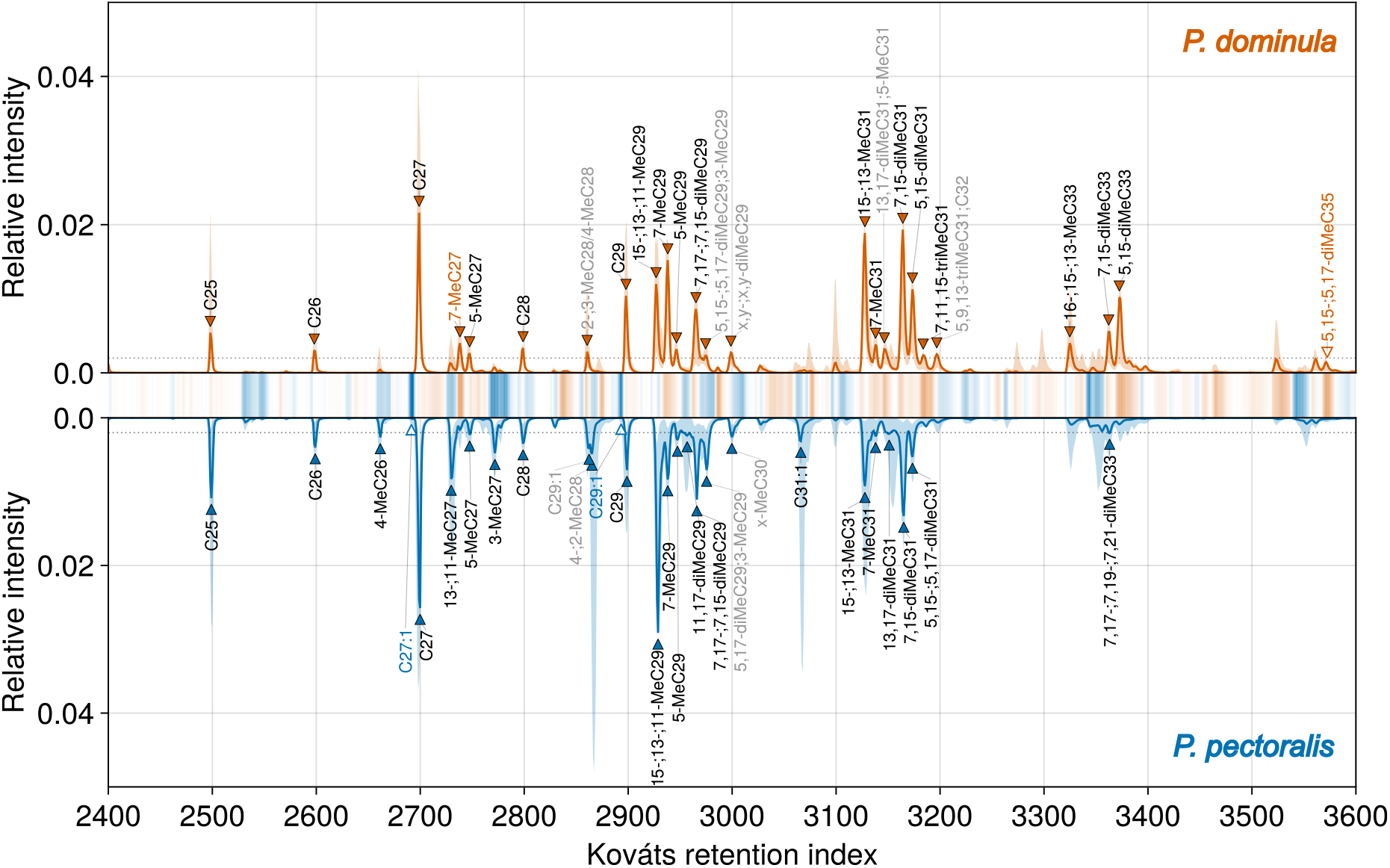
Female representative Total Ion Current (TIC) chromatograms and N-CPLS discriminant coefficient profile. The upper panel shows BOLD:AAA9495 (orange) and the lower panel BOLD:AAB7105 (blue). Solid lines in the two panels show the TIC of score-representative samples selected from the model-fitting dataset, defined as the samples nearest to the class median in the two-dimensional N-CPLS score space spanned by the first two latent variables (ON_13788 for BOLD:AAA9495 and ON_14805 for BOLD:AAB7105). Shaded ribbons show the 5–95% within-group TIC interval. Filled triangles mark peaks detected in the representative sample, open triangles substances of interest without a detected peak there, drawn at their retention index. Labels in a group’s colour mark the two substances whose discriminant coefficients favour that group most strongly (bold in Tab. S3), and faded labels tentative identifications. The central heat map shows the discriminant coefficient profile from the first two latent variables (LV1–LV2), summed over m/z at each Kováts retention index. Colour indicates which group is favoured by the sign of the discriminant coefficient: orange supports BOLD:AAA9495 and blue supports BOLD:AAB7105. Opacity is scaled to the absolute value of the discriminant coefficient, with the largest shown at full opacity. Substances were identified from their mass spectra and Kováts retention indices (Tab. S3).

**Fig. S3:**
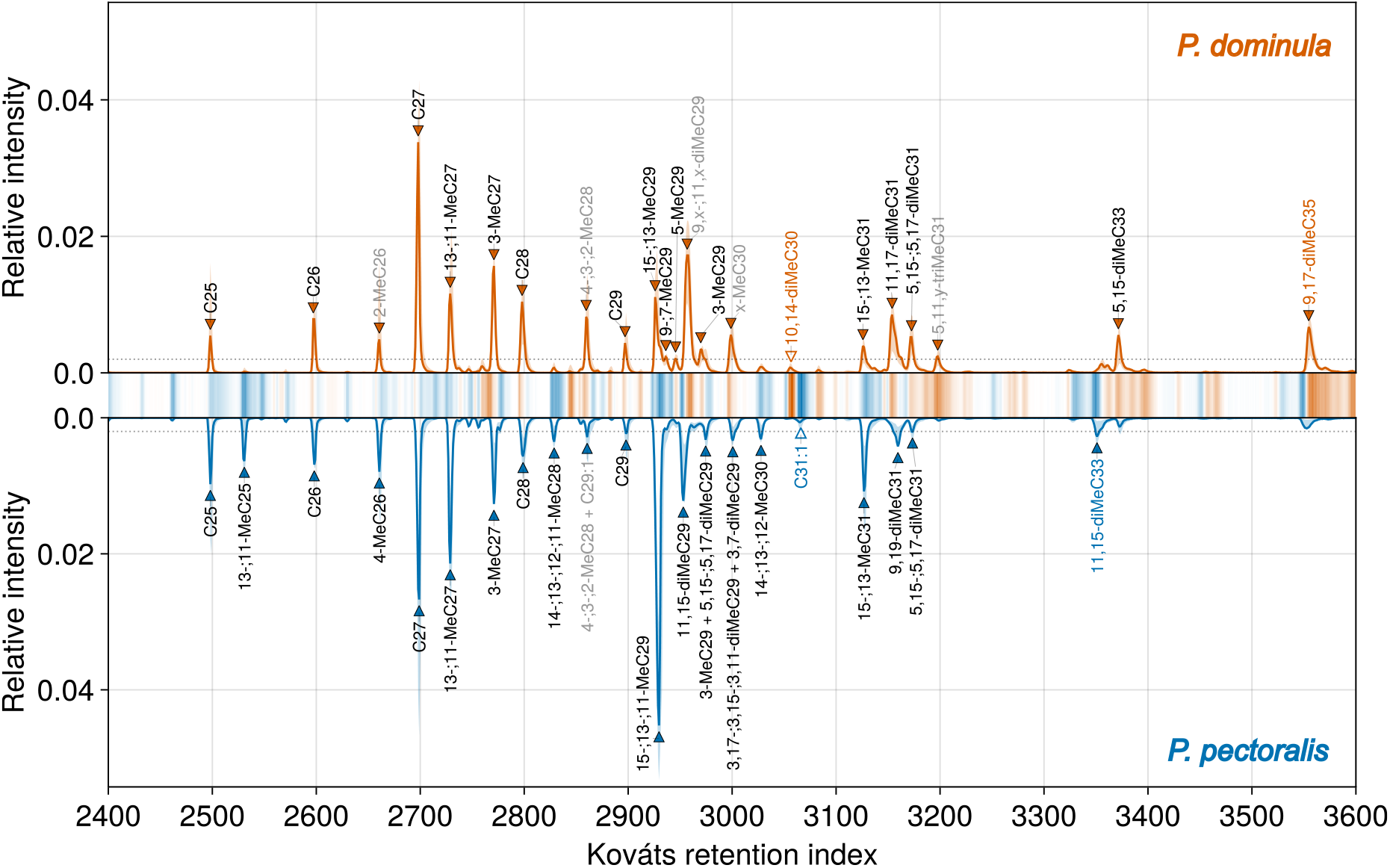
Male representative Total Ion Current (TIC) chromatograms and N-CPLS discriminant coefficient profile. The upper panel shows BOLD:AAA9495 (orange) and the lower panel BOLD:AAB7105 (blue). Solid lines in the two panels show the TIC of score-representative samples selected from the model-fitting dataset, defined as the samples nearest to the class median in the one-dimensional N-CPLS first-latent-variable score space (ON_16123 for BOLD:AAA9495 and ON_14016 for BOLD:AAB7105). Shaded ribbons show the 5–95% within-group TIC interval. Filled triangles mark peaks detected in the representative sample, open triangles substances of interest without a detected peak there, drawn at their retention index. Labels in a group’s colour mark the two substances whose discriminant coefficients favour that group most strongly (bold in Tab. S3), and faded labels tentative identifications. The central heat map shows the discriminant coefficient profile from the first latent variable (LV1), summed over m/z at each Kováts retention index. Colour indicates which group is favoured by the sign of the discriminant coefficient: orange supports BOLD:AAA9495 and blue supports BOLD:AAB7105. Opacity is scaled to the absolute value of the discriminant coefficient, with the largest shown at full opacity. Substances were identified from their mass spectra and Kováts retention indices (Tab. S3).

**Fig. S4:**
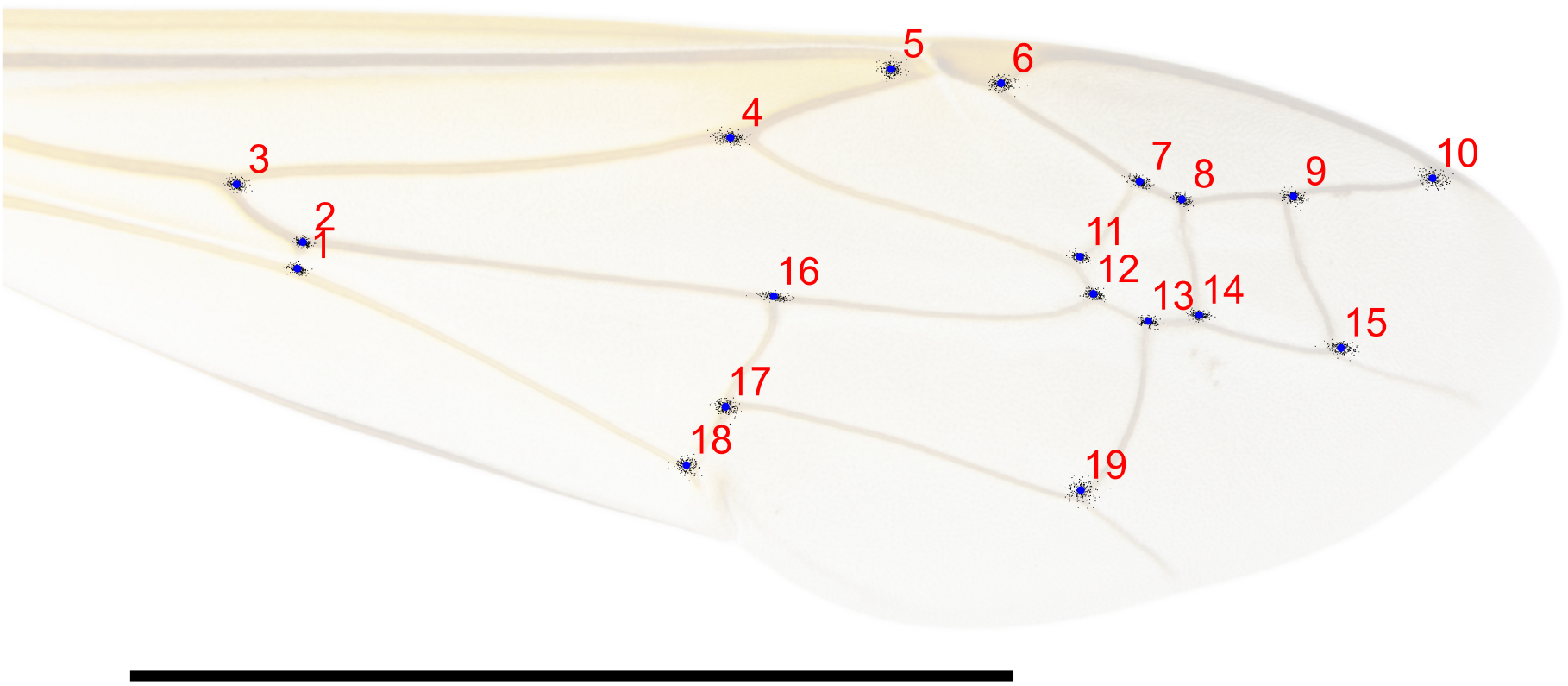
Landmark placement for geometric morphometric analysis of forewing shape in the *P. dominula* species complex, shown on the forewing of ON_13556. Blue: consensus position of landmarks 1–19 after Procrustes superimposition, numbered in red. Black dots: their positions in each of the 200 wings. Landmarks were placed at vein intersections, except for landmarks 5, 6, and 10, which were positioned at the apex or concave corner of the corresponding cells to improve consistency. Scale bar 5 mm.

**Fig. S5:**
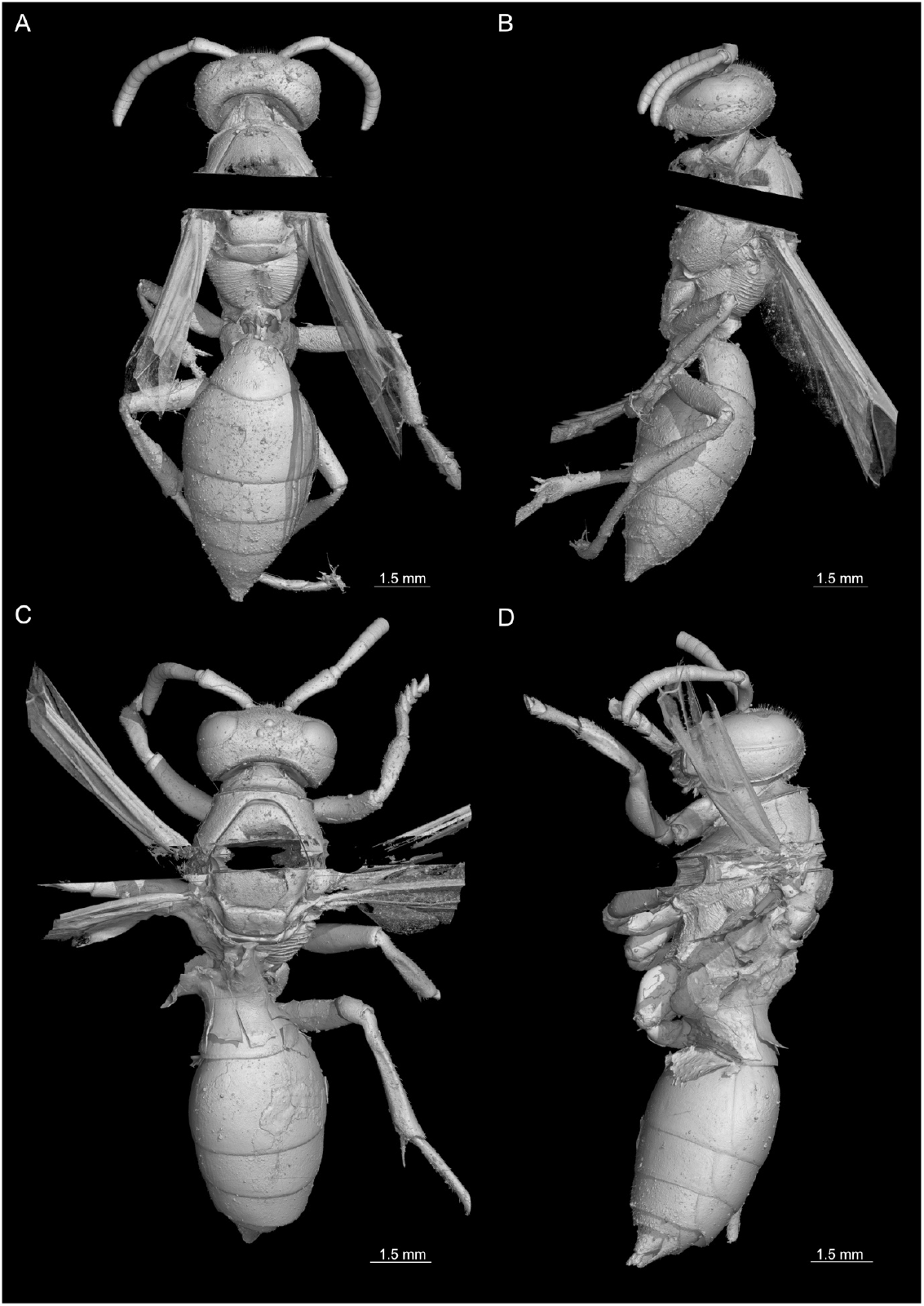
Surface renderings of µCT scans for the two suspected syntype specimens from the Gerning Collection. **A/B:** sample ON_15401, whose somatic morphology is consistent with *P. pectoralis*. **C/D**: sample ON_15402, designated here as the lectotype of *P. dominula* (Tab. S13). Structures missing from the renderings are the shadow of the steel mounting needle, which is opaque to the synchrotron beam

**Fig. S6:**
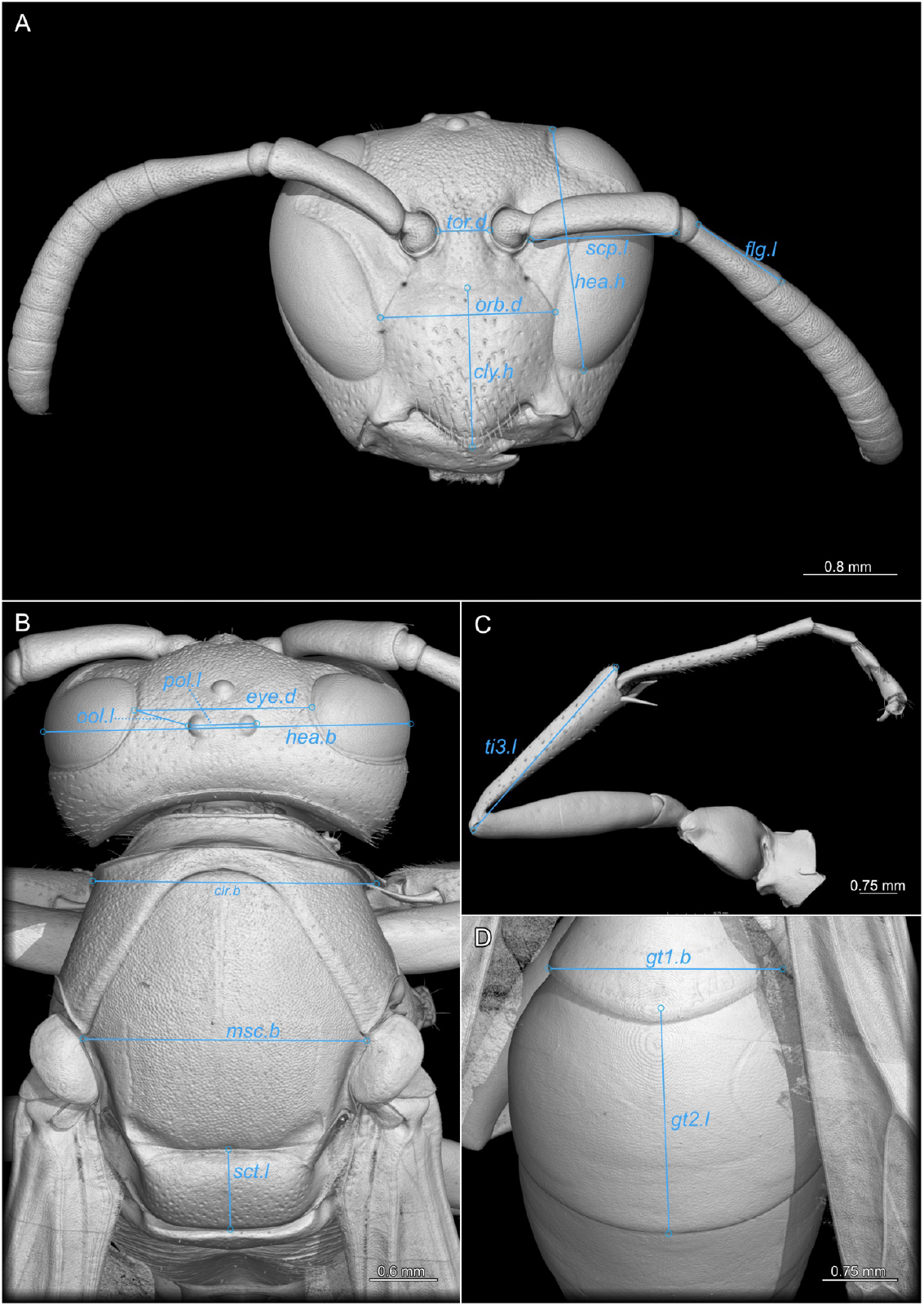
Length measurements for morphometric analysis shown on a surface rendering of a µCT scan for a female of the *P. dominula* complex. The individual measurements are defined in Tab. S5.

**Fig. S7:**
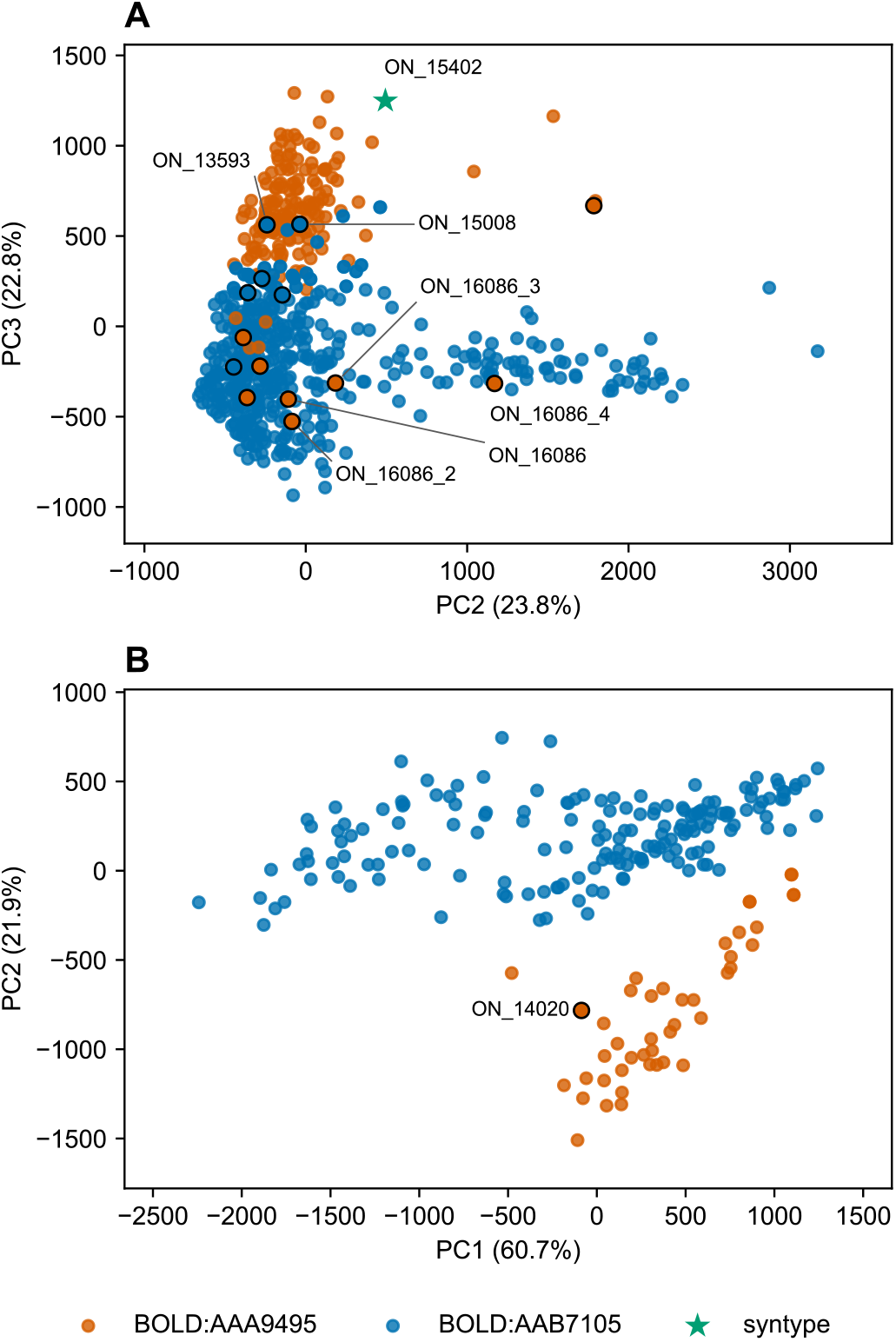
PCA score plot based on cuticular surface profiles for available wasps of the *Polistes dominula* species complex. Points are coloured according to group identity: BOLD:AAA9495 in orange, BOLD:AAB7105 in blue, and the syntype ON_15402 as a green star. **A:** Females (N = 645: 167 assigned to BOLD:AAA9495, 477 assigned to BOLD:AAB7105, and one non-barcoded syntype). Sample scores are plotted for the second and third principal components. PC1 (41.4%) did not separate the two barcode groups (AUC = 0.52, 95% CI 0.47–0.56). Points outlined in black are the eleven females included in the individual genotyping panel (Fig. 2). Individuals with noteworthy phenotype-genotype combinations are labelled, and labels suffixed_2 to _4 denote technical replicates obtained from repeated cuticular surface extraction of wasp ON_16086. **B:** Males (N = 210: 41 assigned to BOLD:AAA9495, 169 assigned to BOLD:AAB7105). Points are coloured according to group identity: BOLD:AAA9495 in orange and BOLD:AAB7105 in blue. Male ON_14020 is outlined in black and labelled because it is the only male included in the individual genotyping panel (Fig. 2), where roughly one quarter of its private-allele states match the other species.

**Fig. S8:**
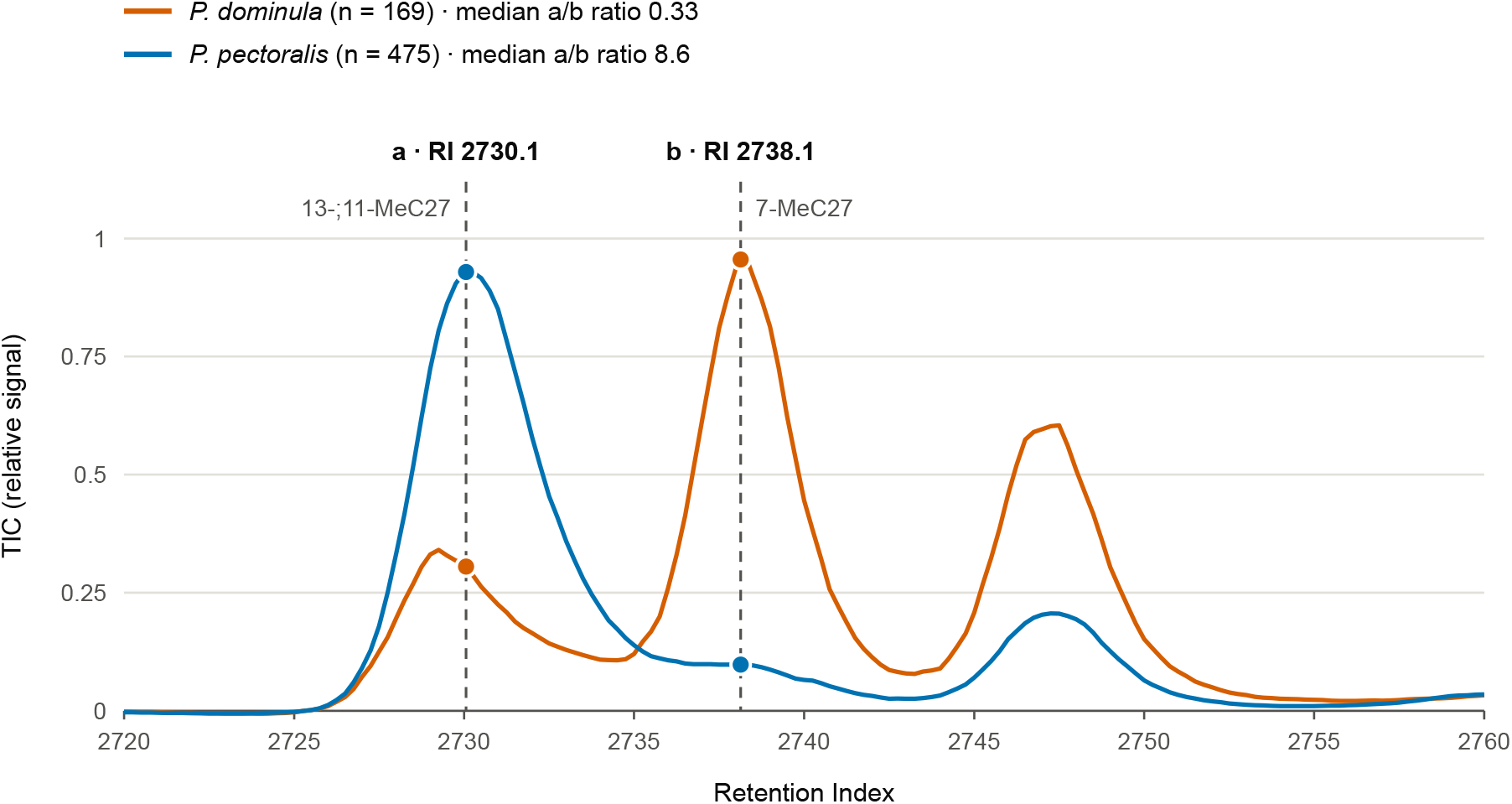
Positions of the female TICC signal ratio. Median Total Ion Current (TIC) of *P. dominula* (n = 169 runs) and *P. pectoralis* (n = 475 runs) females, in relative units, between RI 2720 and 2760. The dashed lines mark the two TICC signals, (**a**) at RI 2730.1 (co-elution cluster of 13-;11-MeC27) and (**b**) at RI 2738.1 (median apex of 7-MeC27).

**Fig. S9:**
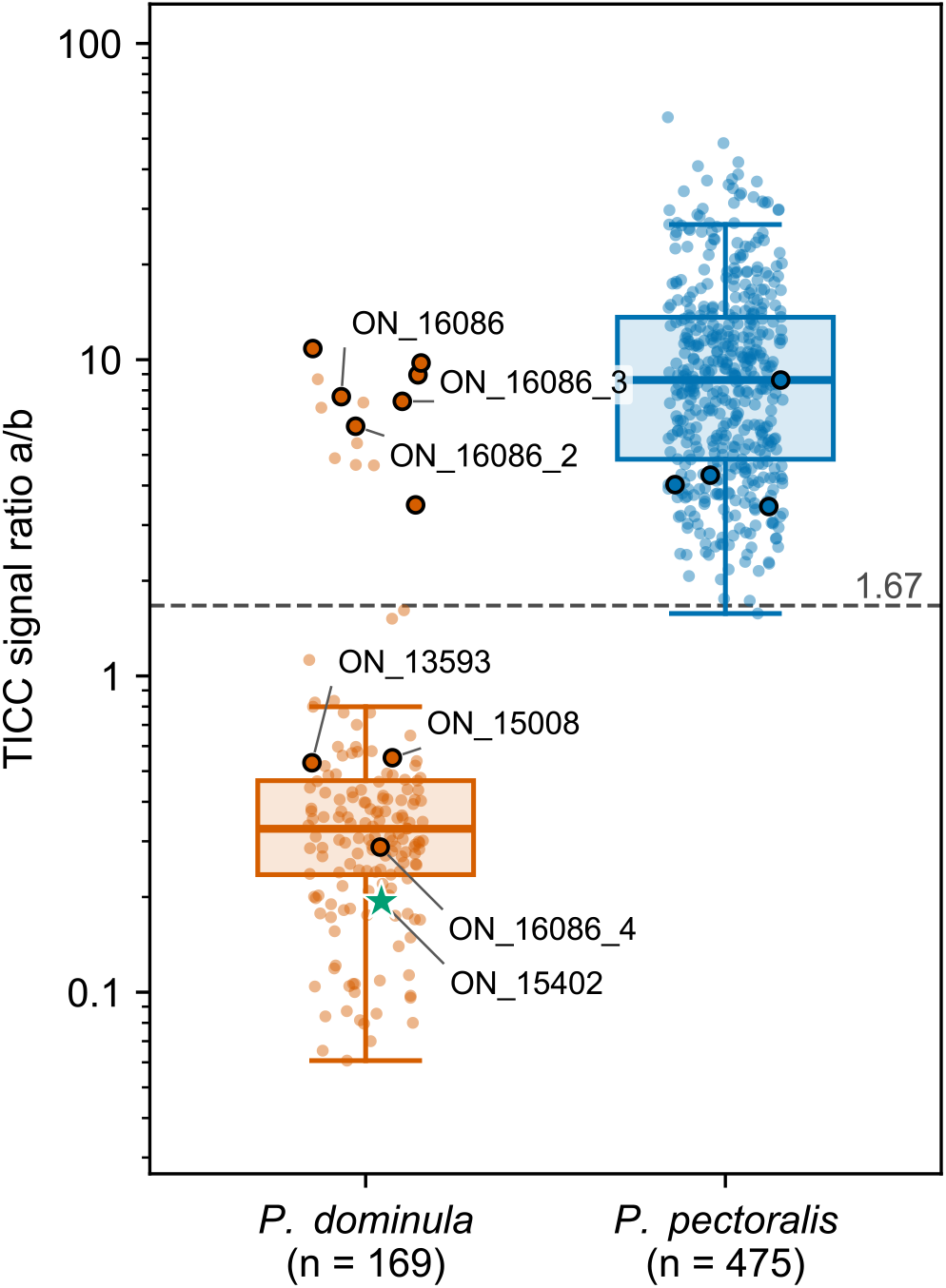
Female TICC signal ratio **a/b** of 13-;11-MeC27 to 7-MeC27 (positions in Fig. S8) by species assignment, one point per run, on a log scale with Tukey-convention box plots. Above the dashed threshold of 1.67 a run is predicted to be *P. pectoralis* (AUC = 0.966, LOOCV balanced accuracy 95.2%). Points outlined in black are the eleven females of the individual genotyping panel (Fig. 2); labels suffixed _2 to _4 denote technical replicates of ON_16086. The syntype ON_15402, lectotype of *P. dominula*, is shown as a green star among the *P. dominula* runs and was not included in the threshold-setting procedure.

**Fig. S10:**
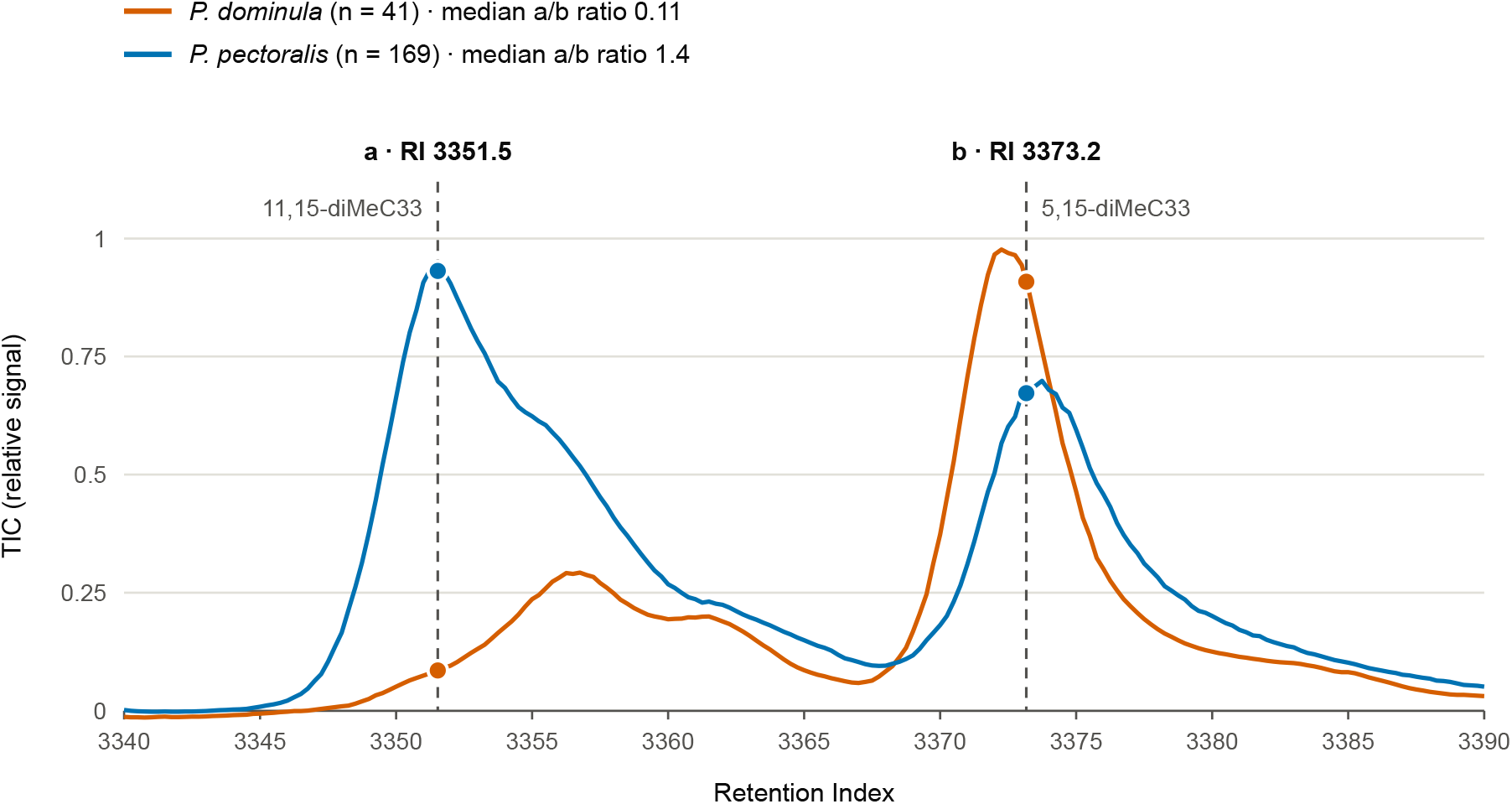
Positions of the male TICC signal ratio. Median Total Ion Current (TIC) of *P. dominula* (n= 41 runs) and *P. pectoralis* (n = 169 runs) males, in relative units, between RI 3340 and 3390. The dashed lines mark the two TICC signals, (**a**) at RI 3351.5 (overall median apex of 11,15-diMeC33) and (**b**) at RI 3373.2 (overall median apex of 5,15-diMeC33).

**Fig. S11:**
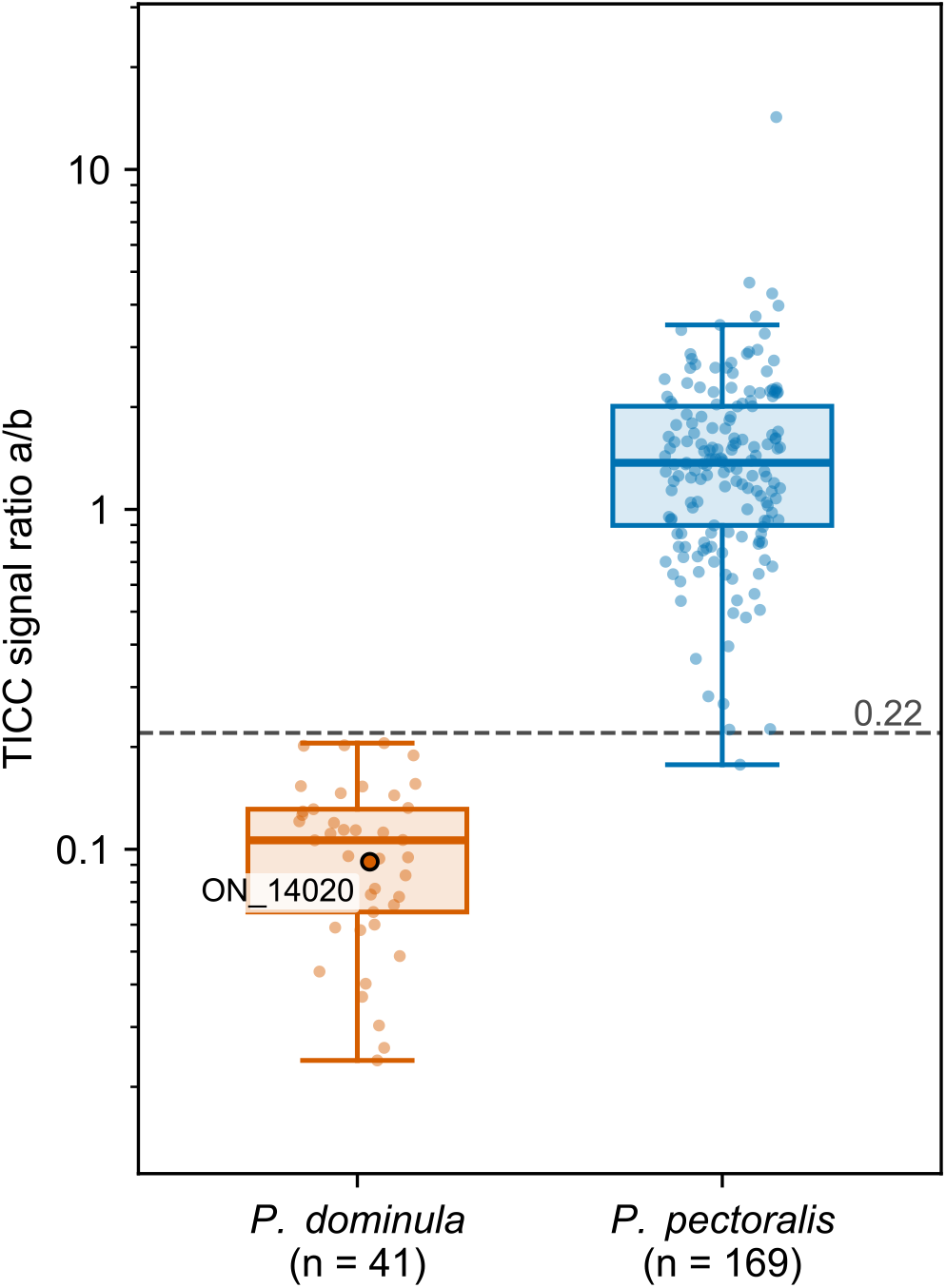
Male TICC signal ratio **a/b** of 11,15-diMeC33 to 5,15-diMeC33 (positions in Fig. S10) by species assignment, one point per run, on a log scale. Boxes as in Fig. S9. Above the dashed threshold of 0.22 a run is predicted to be *P. pectoralis* (AUC = 0.999, LOOCV balanced accuracy 99.7%). The point outlined in black is ON_14020, of the individual genotyping panel (Fig. 2).

**Fig. S12:**
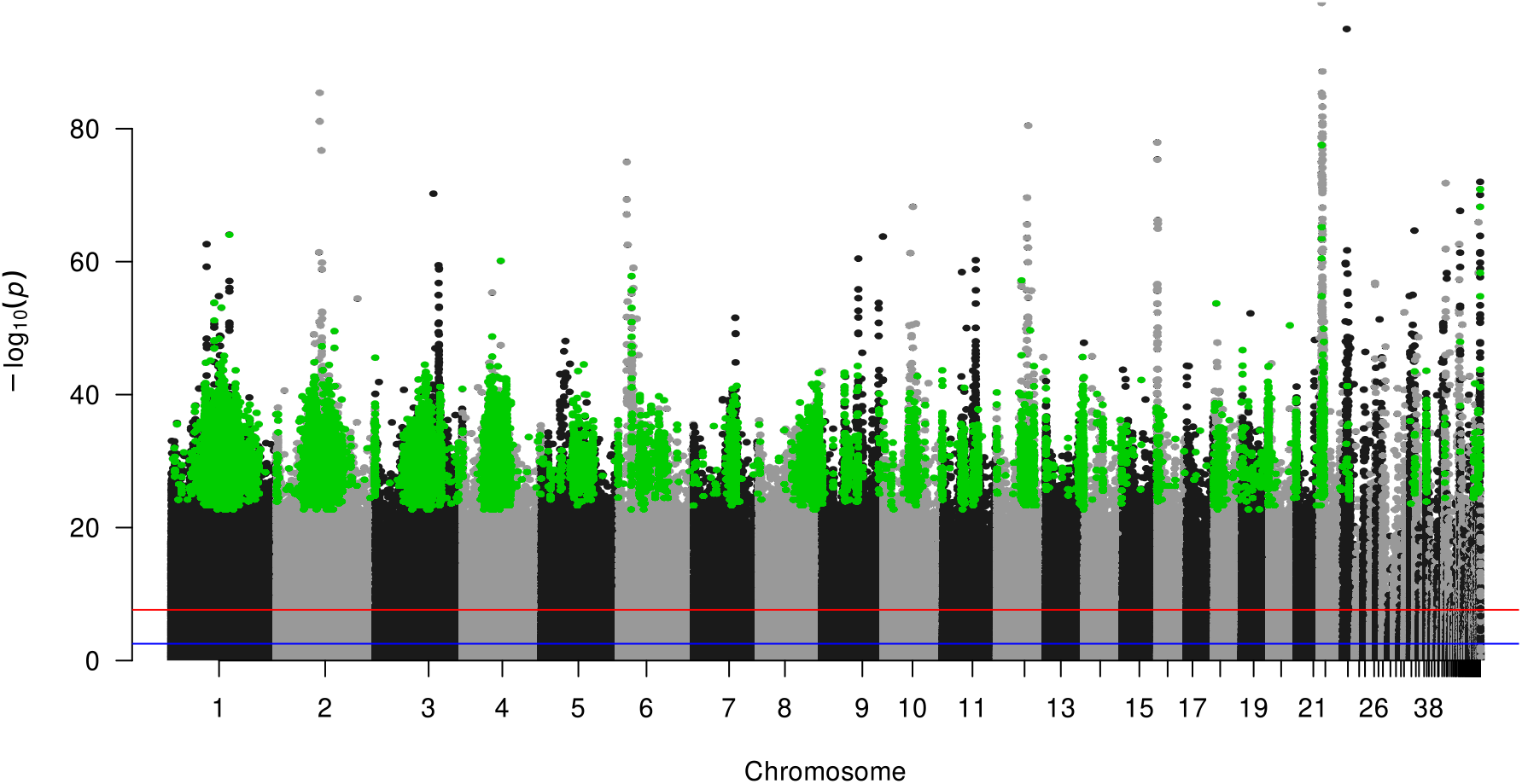
Allele frequency differences across the genome for biallelic SNPs. The x-axis represents the genomic position of SNPs projected into genome space and the y-axis shows the result for Fisher’s exact test for allele counts on a negative log scale. Private alleles are shown in green. The red line shows the FDR corrected significance threshold (*p* < 0.003), the blue line shows the Bonferroni corrected significance threshold (*p* < 2.31e-08).

**Fig. S13:**
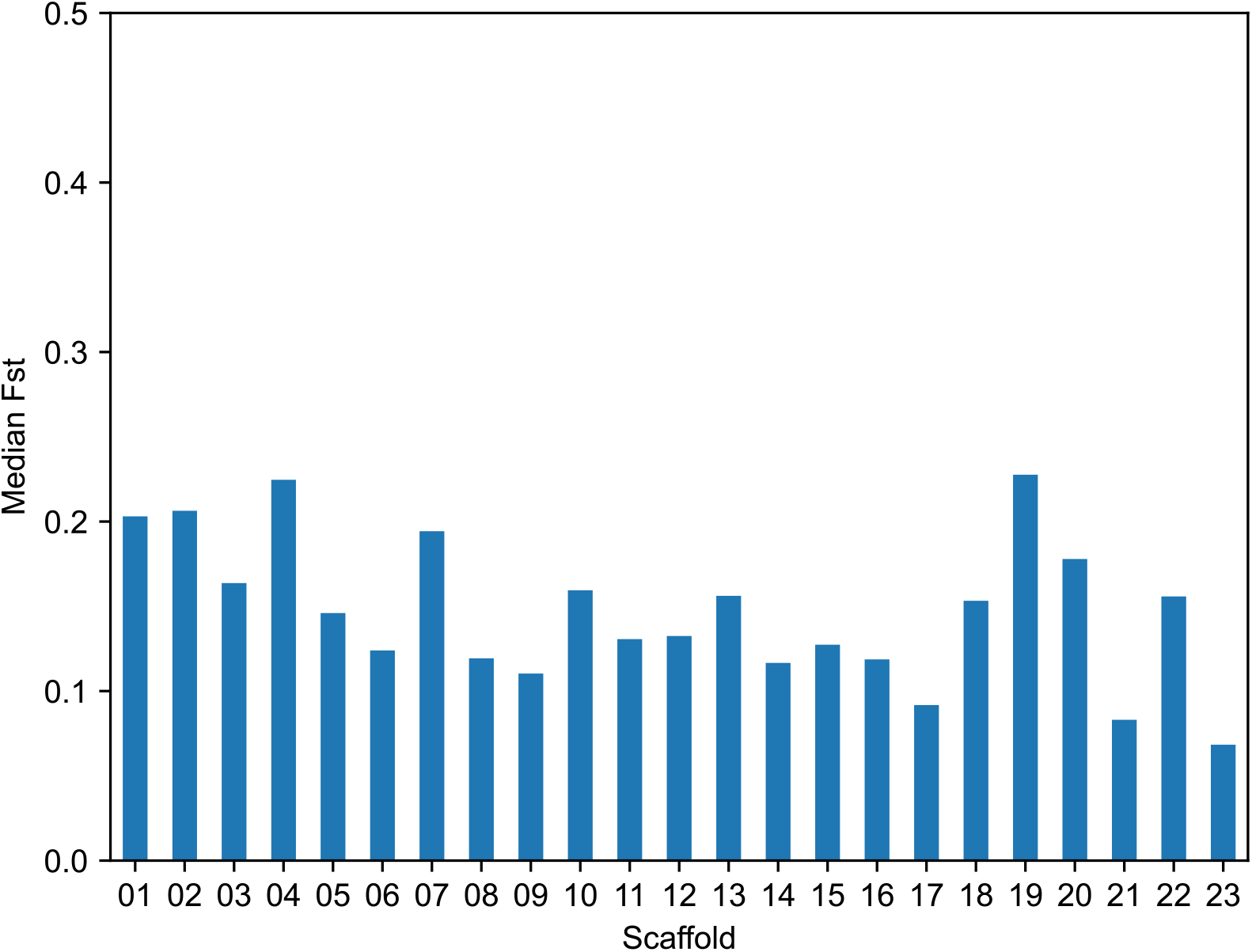
Median F_st_ per N90-scaffold for all detected SNPs excluding indels, based on genome-wide Pool-Seq data in the *Polistes dominula* species complex. Individuals were pooled based on BIN mitochondrial haplotype identity (54 BOLD:AAA9495 females and 54 BOLD:AAB7105 females)

**Fig. S14:**
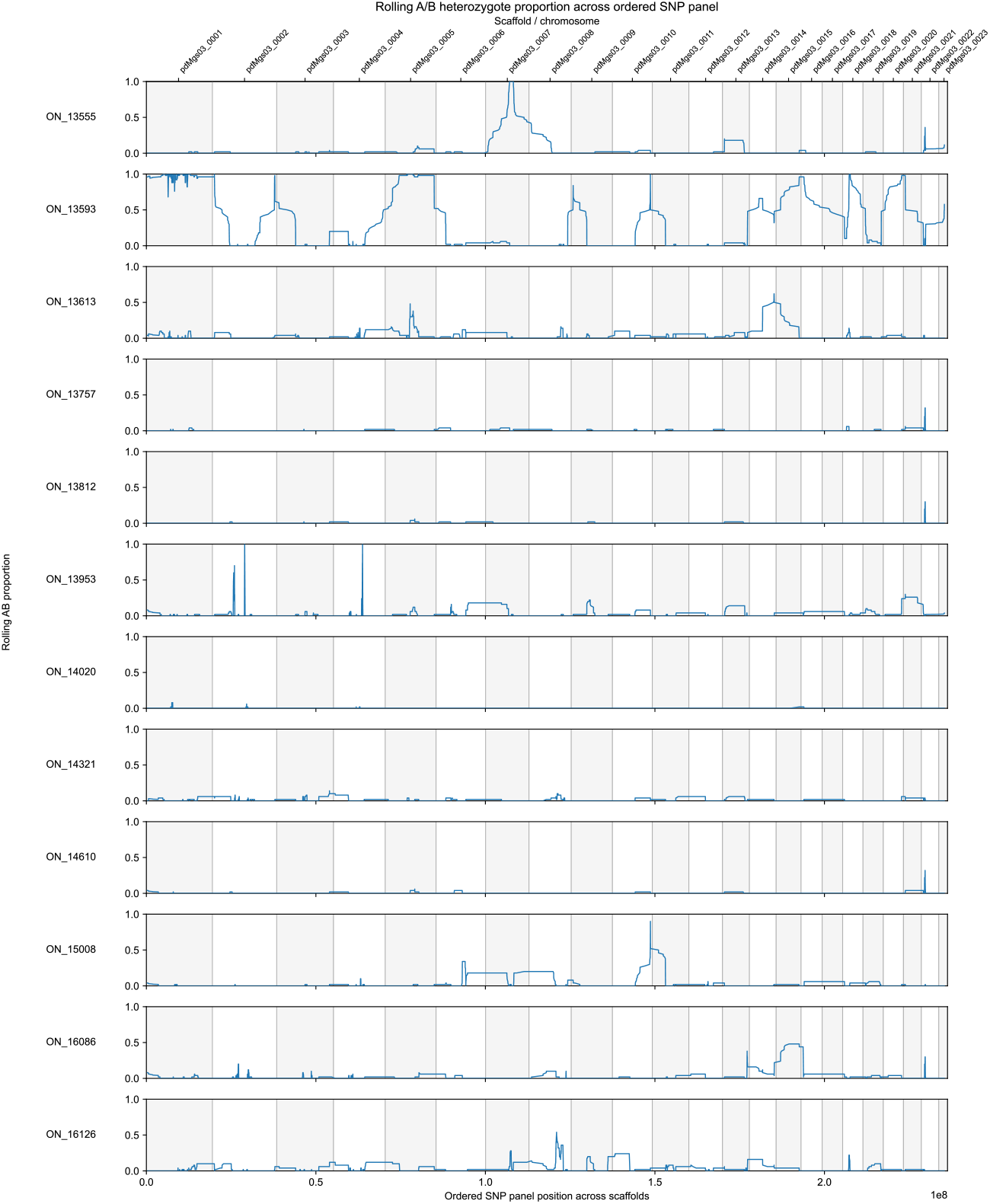
Sliding window heterozygosity analysis for private allele panel SNPs in individuals. The x-axis represents the genomic position of SNPs projected into genome space and the y-axis represents the proportion of heterozygous SNPs within each sliding window (50 SNPs window size). Note the highly elevated heterozygosity on scaffolds 1 and 5 for individual ON_13593 and that individual ON_14020 is the only male.

**Fig. S15:**
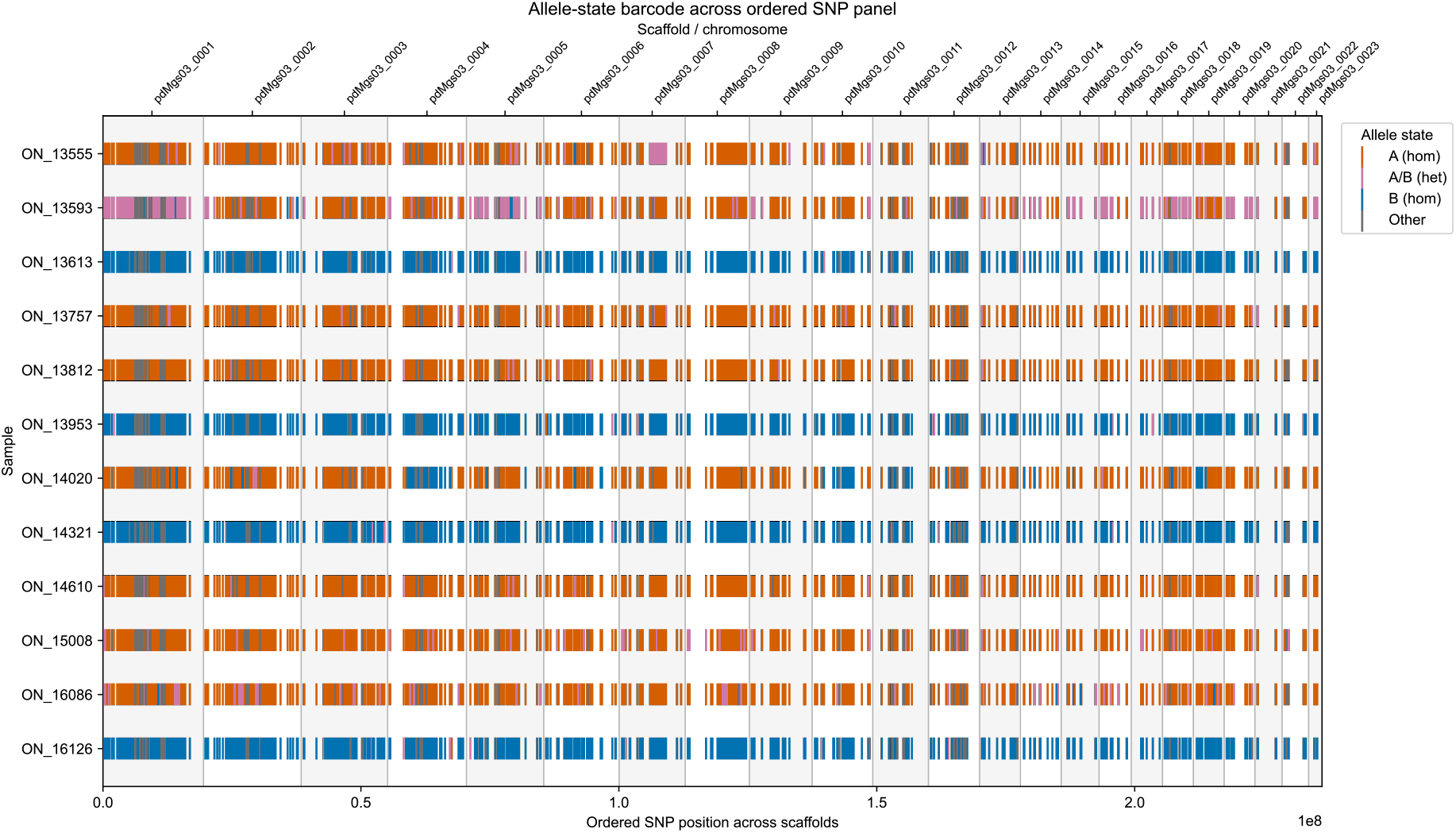
Allele states for private allele panel SNPs in slected individuals across the genome. Detected SNP alleles were coloured as matching Pool A (orange), matching Pool B (blue), matching a heterozygous state with both alleles present (magenta), or matching any other allele combination (grey).

**Fig. S16:**
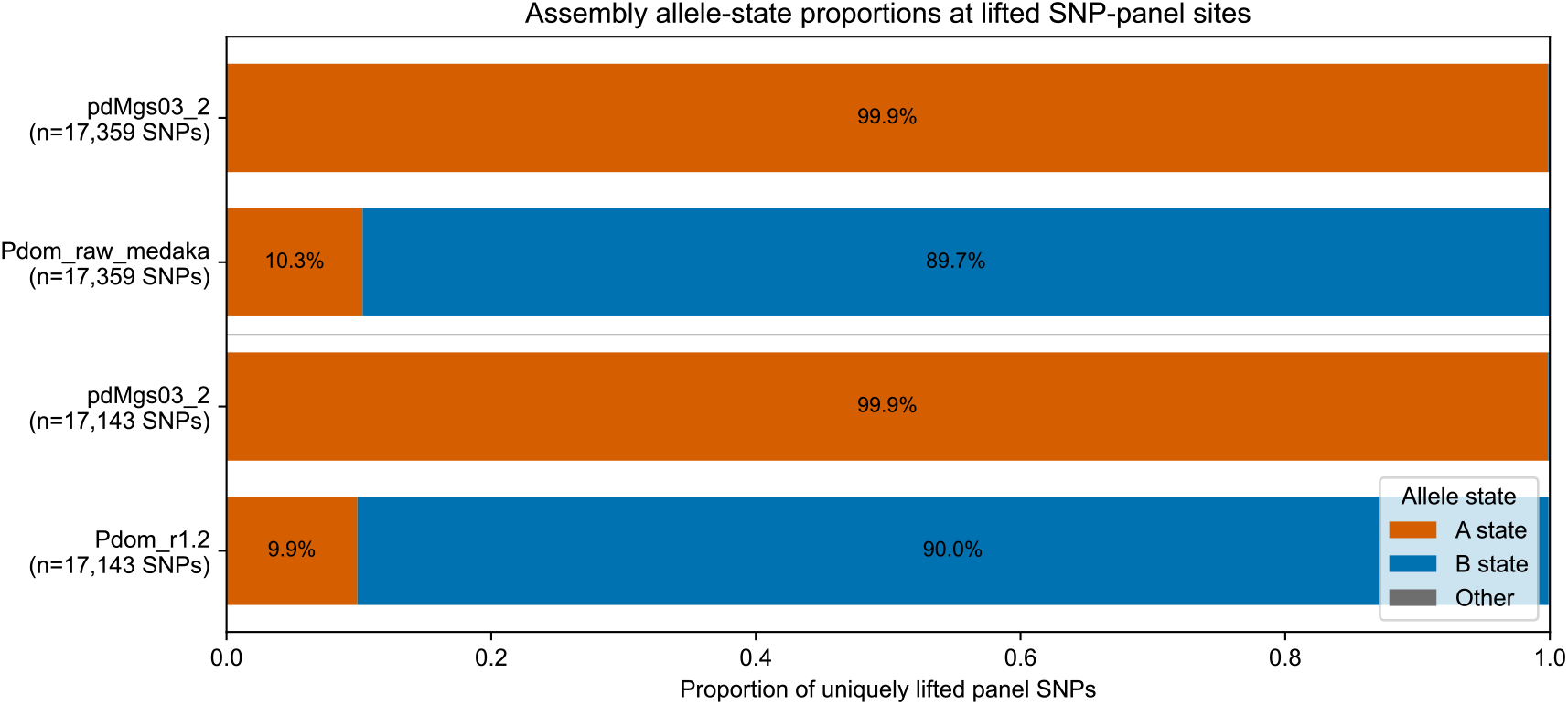
Comparison of allele states across different reference genome assemblies for a panel of 18,026 SNPs. SNP coordinates were lifted over by whole genome alignment and considered, where unique mapping was possible. Allele counts are summarised as matching Pool A (orange), matching Pool B (blue) or matching a different nucleotide (grey).

**Fig. S17:**
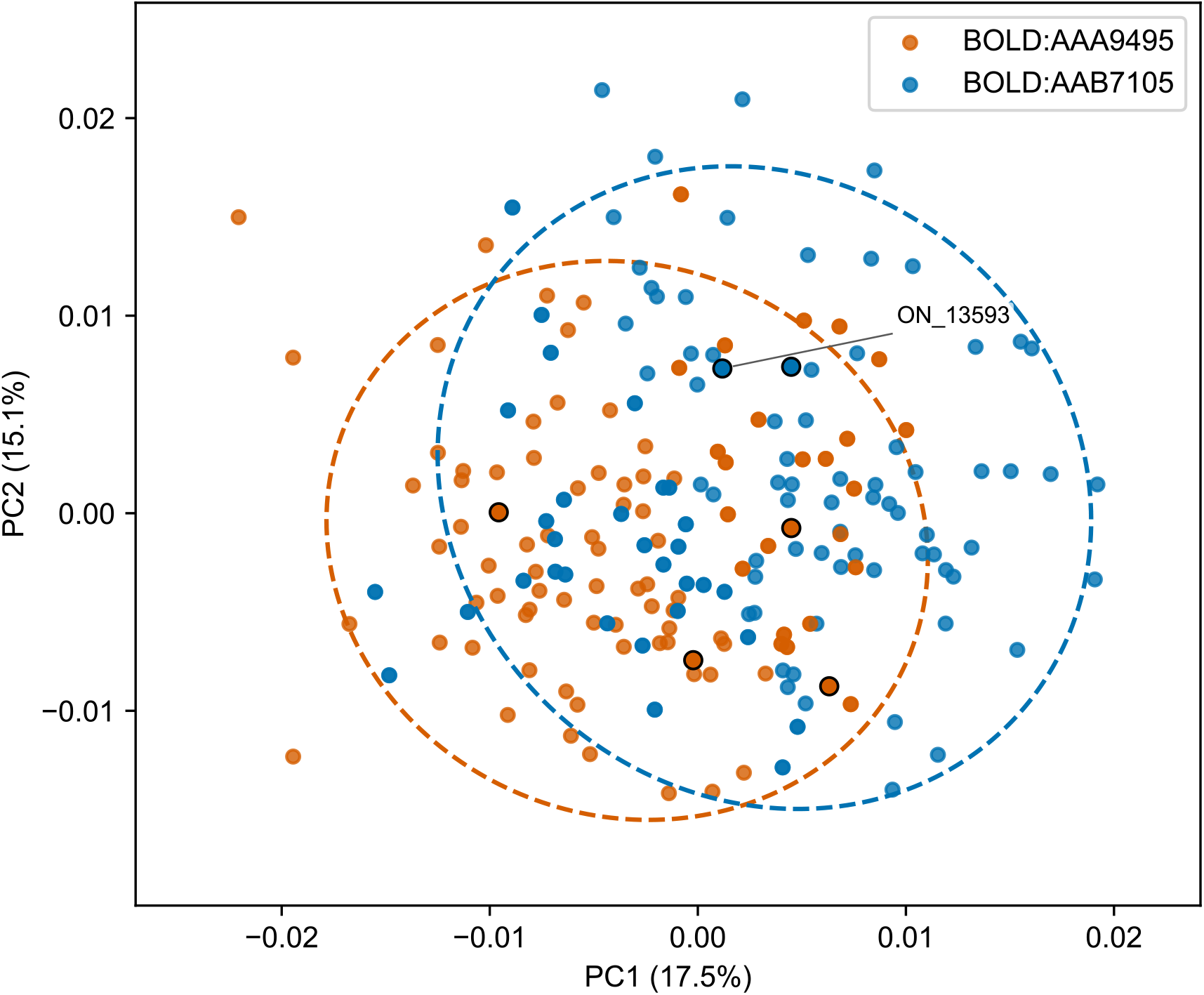
PCA of forewing shape variation in the *P. dominula* species complex for the first and second PC. Each point represents the forewing shape of an individual wasp, coloured according to its BIN identity (BOLD:AAA9495 in orange, BOLD:AAB7105 in blue).

**Fig. S18:**
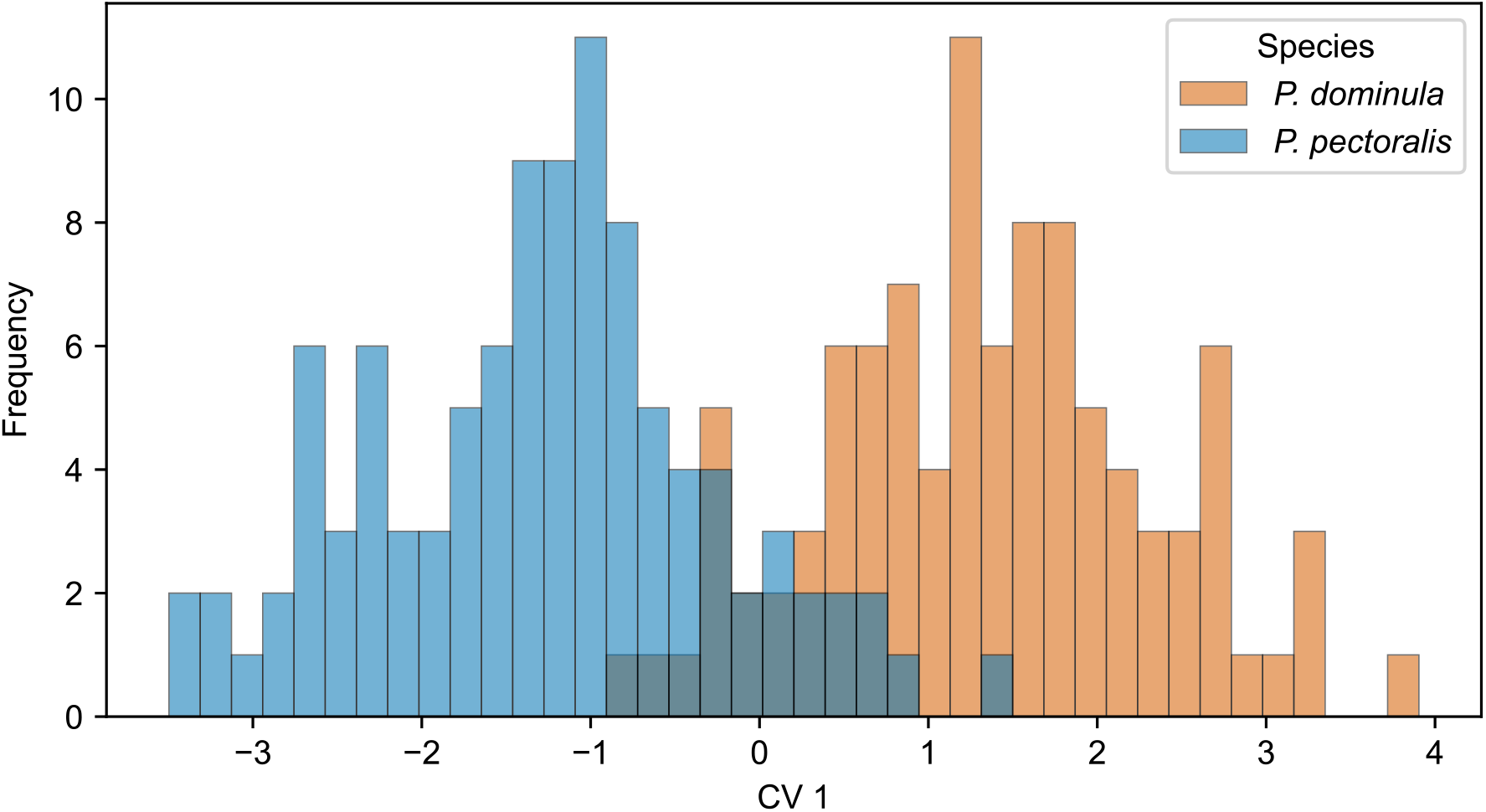
Distribution of discriminating wing shape variation in the *P. dominula* species complex. Counts were binned into 40 bins along the first canonical variate (CV1) and coloured according to species assignment (*P. dominula* in orange, *P. pectoralis* in blue). Labelled samples are included in the individual genotyping panel (Fig. 2)

**Fig. S19:**
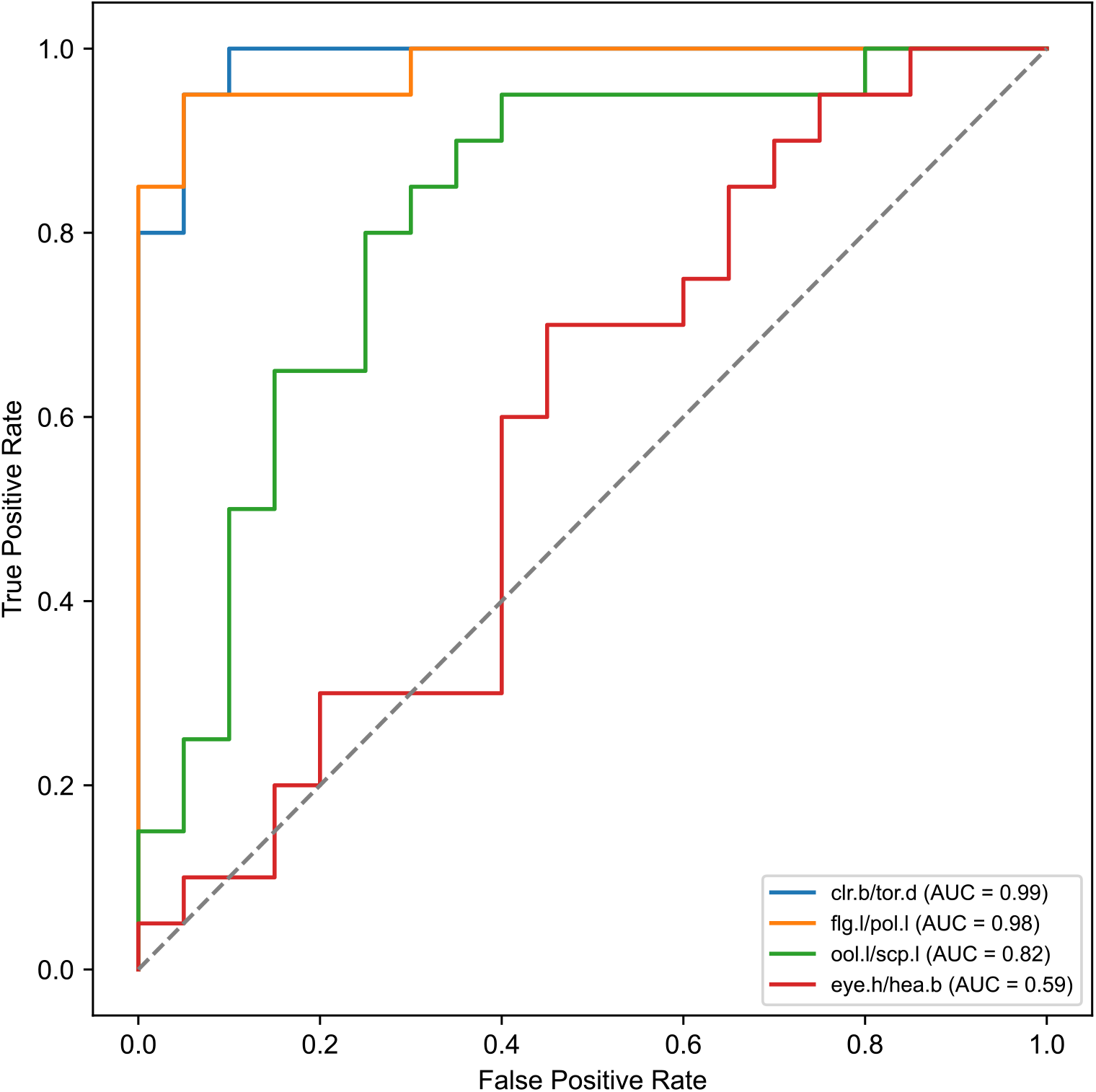
Receiver Operating Characteristic (ROC) curve analysis for the four most discriminating character length ratios from the LDA ratio extraction of 3D somatic morphometric data (n = 40). The legend shows the Area Under the Curve (AUC) metric for each ratio. For a given classification threshold between the minimum and maximum value present in the dataset, the false positive rate is shown on the x-axis and the true positive rate on the y-axis. The **clr.b/tor.d** and **flg.l/pol.l** ratios perform well in species discrimination, whereas the **ool.l/scp.l** ratio is less effective and the **eye.h/hea.b** ratio performs barely above chance.

**Fig. S20:**
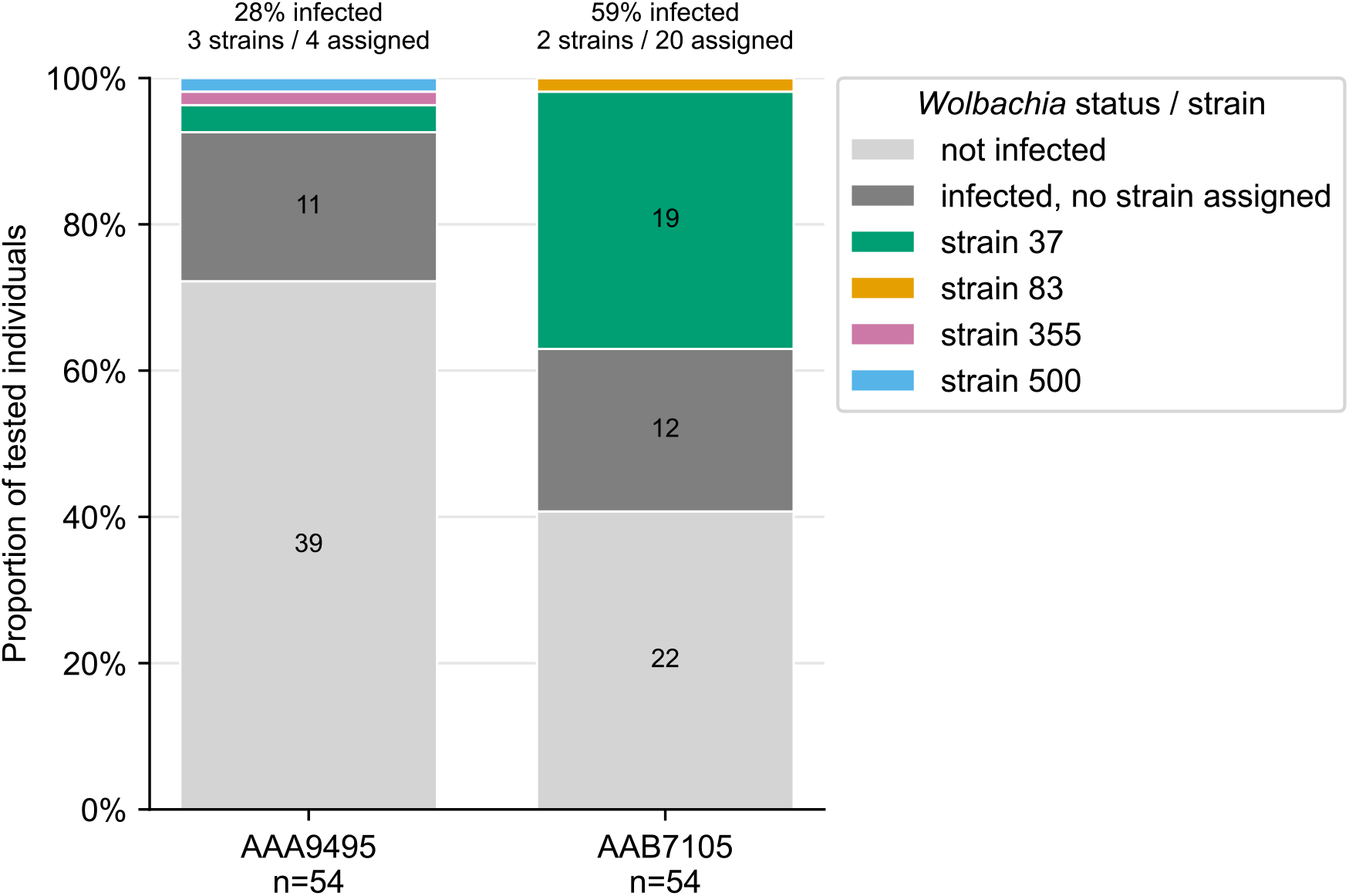
Prevalence and detected strain types of *Wolbachia* in sympatrically occurring individuals of the *P. dominula* complex belonging to different mitochondrial haplotypes (BOLD:AAA9495 left bar, BOLD:AAB7105 right bar)

**Fig. S21:**
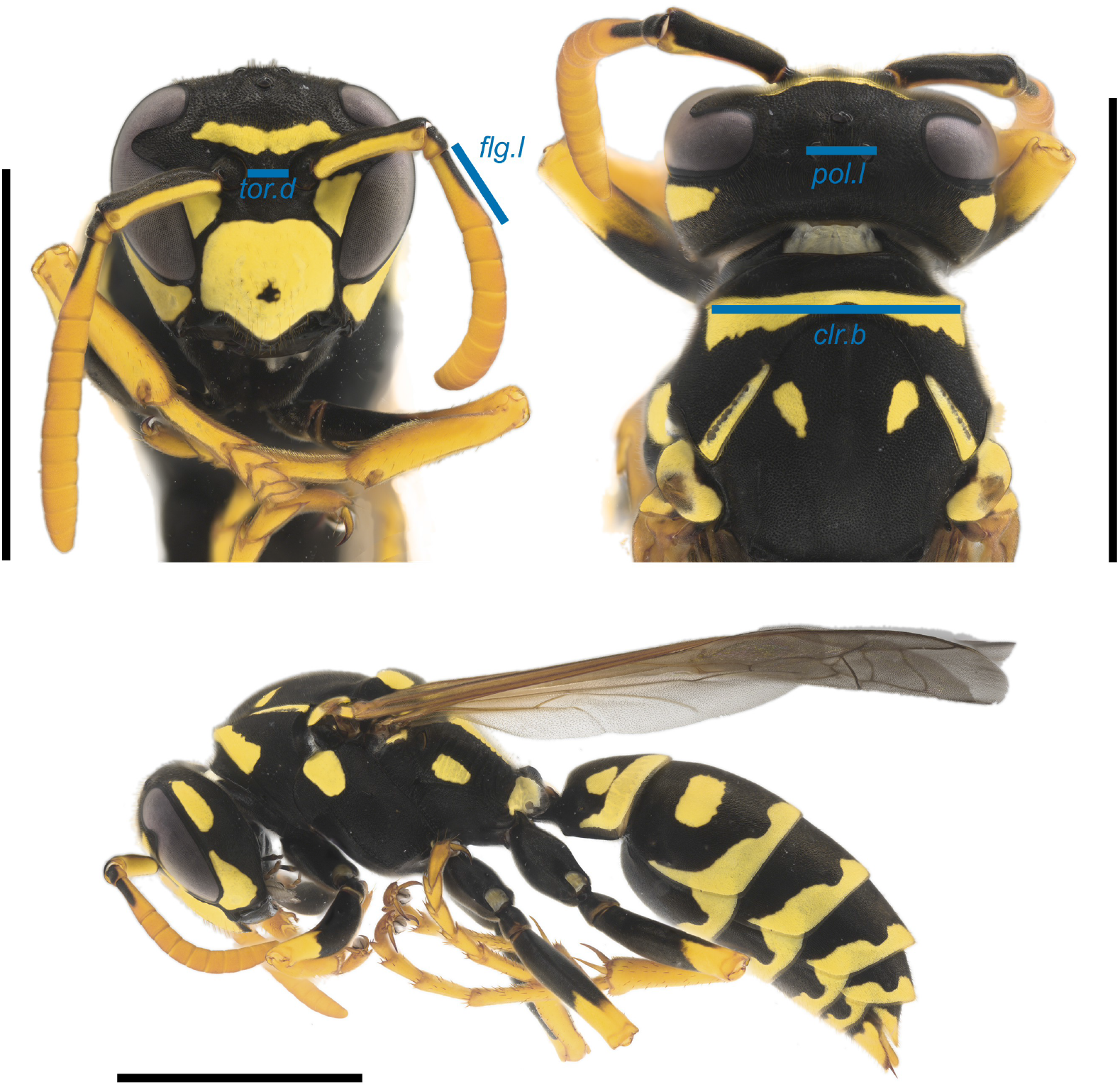
Neotype of *Polistes pectoralis* (♀, ON_15015 and ZFMK-TIS-89039 at Museum Koenig). Focus-stacked image of the head in frontal view (top left), head and pronotum in dorsal view (top right) and habitus in lateral view (bottom). Blue bars mark the four distances of the diagnostic ratios **clr.b/tor.d** and **flg.l/pol.l** (Tab. S5). *clr.b*: breadth of the pronotal collar. *tor.d*: distance between the antennal toruli. *flg.l*: length of the first flagellomere. *pol.l*: distance between the outer edges of the lateral ocelli. Colours were matched to a colour-calibrated photograph of a live *P. dominula* s.l. Black scale bars are 5 mm.

## Supplementary Tables

**Tab. S1:** Contingency table of COI BOLD BIN assignment and ITS1 tree clade assignment for specimens of the *Polistes dominula* complex in Neumeyer et al. [13]. ITS1 clades were inferred from the published ITS1 maximum-likelihood tree topology. The truncated/not placed category refers to ITS1 sequences not visible in the provided tree screenshot.

| COI BOLD<br>BIN | ITS1 upper clade | ITS1 lower clade | ITS1 missing | Total |
|---|---|---|---|---|
| BOLD:AAA9495 | 0 | 6 | 0 | 6 |
| BOLD:AAB7105 | 7 | 4 | 1 | 12 |
| **Total** | **7** | **10** | **1** | **18** |

**Tab. S2:**
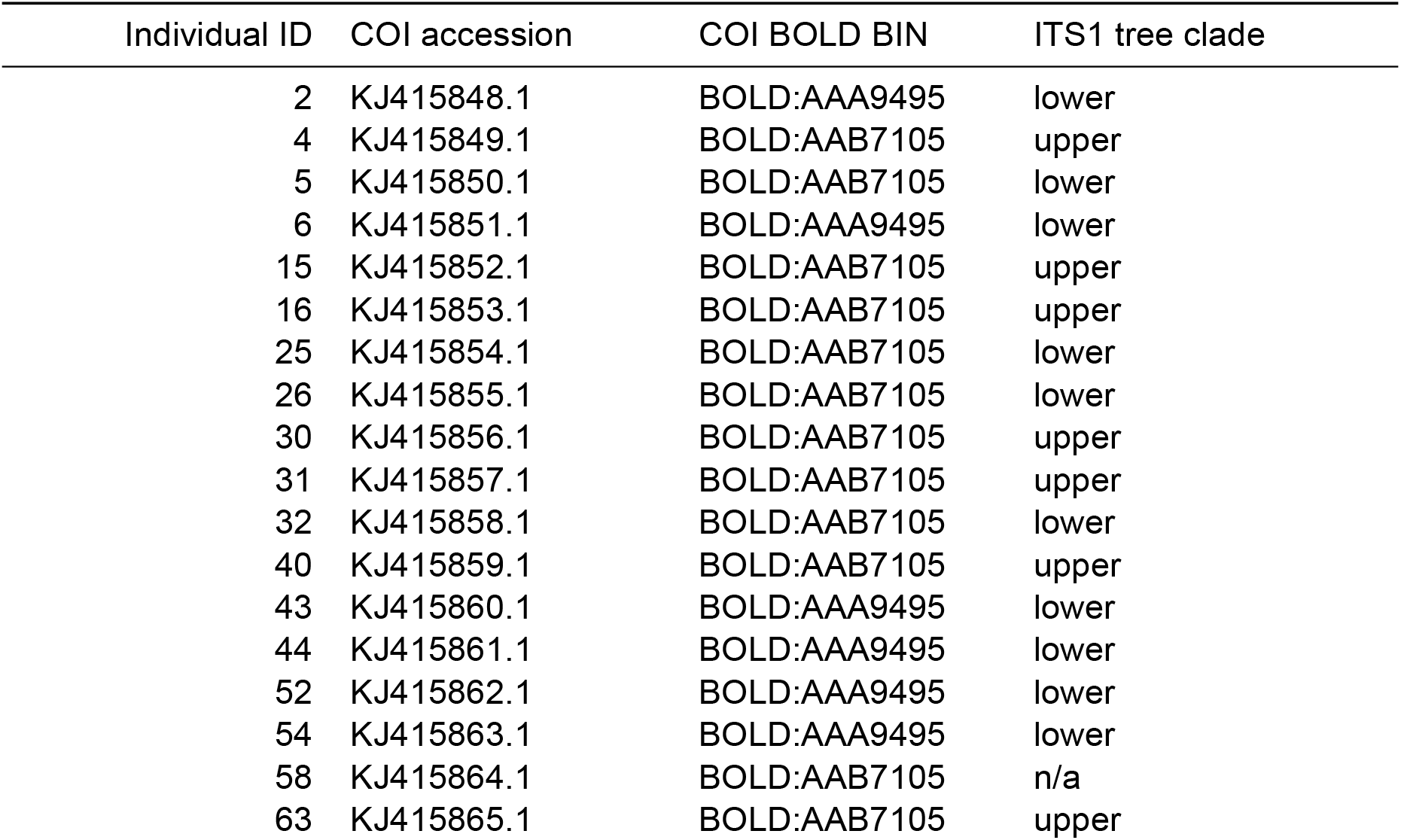
Individual-level mapping between COI accession, COI BOLD BIN assignment, and ITS1 tree clade assignment. ITS1 clade assignment follows the visible topology of the published ITS1 maximum-likelihood tree. Individual 58 has no associated ITS1 sequence.

| Individual ID | COI accession | COI BOLD BIN | ITS1 tree clade |
|---|---|---|---|
| 2 | KJ415848.1 | BOLD:AAA9495 | lower |
| 4 | KJ415849.1 | BOLD:AAB7105 | upper |
| 5 | KJ415850.1 | BOLD:AAB7105 | lower |
| 6 | KJ415851.1 | BOLD:AAA9495 | lower |
| 15 | KJ415852.1 | BOLD:AAB7105 | upper |
| 16 | KJ415853.1 | BOLD:AAB7105 | upper |
| 25 | KJ415854.1 | BOLD:AAB7105 | lower |
| 26 | KJ415855.1 | BOLD:AAB7105 | lower |
| 30 | KJ415856.1 | BOLD:AAB7105 | upper |
| 31 | KJ415857.1 | BOLD:AAB7105 | upper |
| 32 | KJ415858.1 | BOLD:AAB7105 | lower |
| 40 | KJ415859.1 | BOLD:AAB7105 | upper |
| 43 | KJ415860.1 | BOLD:AAA9495 | lower |
| 44 | KJ415861.1 | BOLD:AAA9495 | lower |
| 52 | KJ415862.1 | BOLD:AAA9495 | lower |
| 54 | KJ415863.1 | BOLD:AAA9495 | lower |
| 58 | KJ415864.1 | BOLD:AAB7105 | n/a |
| 63 | KJ415865.1 | BOLD:AAB7105 | upper |

**Tab. S3:**
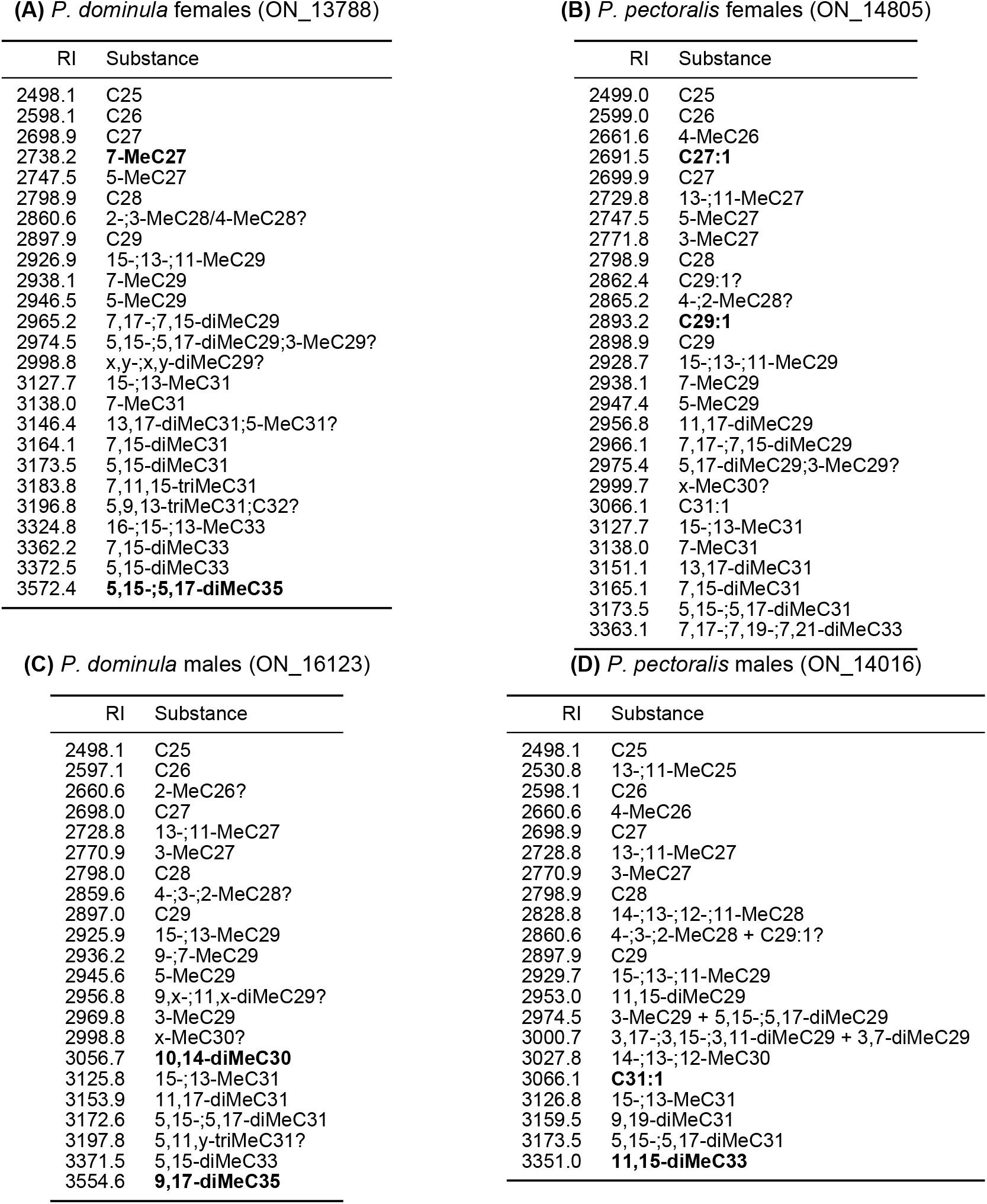
Substances identified in the representative total ion current chromatograms (Fig. S2, Fig. S3) of the females (**A**, **B**) and males (**C**, **D**) of both species. The sample each subtable annotates is given in parentheses. RI: measured Kováts retention index. Substance labels are analogous to the figures. C*N* denotes *n*-alkanes, Me, diMe and triMe one, two or three methyl branches at the numbered positions (x and y where a position is unknown), and C*N*:1 an alkene. Semicolons and plus signs separate substances that co-elute in one peak, a slash separates alternative identifications, and a question mark follows a tentative identification. Bold labels mark, for each sex and species, the two peaks whose N-CPLS discriminant coefficients favour that species most strongly. Detection frequencies for the individuals of either species are given in Tab. S8.

| **(A)** *P. dominula* females (ON_13788) |  | **(B)** *P. pectoralis* females (ON_14805) |  | **(C)** *P. dominula* males (ON_16123) |  | **(D)** *P. pectoralis* males (ON_14016) |  |
|---|---|---|---|---|---|---|---|
| RI | Substance | RI | Substance | RI | Substance | RI | Substance |
| 2498.1 | C25 | 2499.0 | C25 | 2498.1 | C25 | 2498.1 | C25 |
| 2598.1 | C26 | 2599.0 | C26 | 2597.1 | C26 | 2530.8 | 13-;11-MeC25 |
| 2698.9 | C27 | 2661.6 | 4-MeC26 | 2660.6 | 2-MeC26? | 2598.1 | C26 |
| 2738.2 | **7-MeC27** | 2691.5 | **C27:1** | 2698.0 | C27 | 2660.6 | 4-MeC26 |
| 2747.5 | 5-MeC27 | 2699.9 | C27 | 2728.8 | 13-;11-MeC27 | 2698.9 | C27 |
| 2798.9 | C28 | 2729.8 | 13-;11-MeC27 | 2770.9 | 3-MeC27 | 2728.8 | 13-;11-MeC27 |
| 2860.6 | 2-;3-MeC28/4-MeC28? | 2747.5 | 5-MeC27 | 2798.0 | C28 | 2770.9 | 3-MeC27 |
| 2897.9 | C29 | 2771.8 | 3-MeC27 | 2859.6 | 4-;3-;2-MeC28? | 2798.9 | C28 |
| 2926.9 | 15-;13-;11-MeC29 | 2798.9 | C28 | 2897.0 | C29 | 2828.8 | 14-;13-;12-;11-MeC28 |
| 2938.1 | 7-MeC29 | 2862.4 | C29:1? | 2925.9 | 15-;13-MeC29 | 2860.6 | 4-;3-;2-MeC28 + C29:1? |
| 2946.5 | 5-MeC29 | 2865.2 | 4-;2-MeC28? | 2936.2 | 9-;7-MeC29 | 2897.9 | C29 |
| 2965.2 | 7,17-;7,15-diMeC29 | 2893.2 | **C29:1** | 2945.6 | 5-MeC29 | 2929.7 | 15-;13-;11-MeC29 |
| 2974.5 | 5,15-;5,17-diMeC29;3-MeC29? | 2898.9 | C29 | 2956.8 | 9,x-;11,x-diMeC29? | 2953.0 | 11,15-diMeC29 |
| 2998.8 | x,y-;x,y-diMeC29? | 2928.7 | 15-;13-;11-MeC29 | 2969.8 | 3-MeC29 | 2974.5 | 3-MeC29 + 5,15-;5,17-diMeC29 |
| 3127.7 | 15-;13-MeC31 | 2938.1 | 7-MeC29 | 2998.8 | x-MeC30? | 3000.7 | 3,17-;3,15-;3,11-diMeC29 + 3,7-diMeC29 |
| 3138.0 | 7-MeC31 | 2947.4 | 5-MeC29 | 3056.7 | **10,14-diMeC30** | 3027.8 | 14-;13-;12-MeC30 |
| 3146.4 | 13,17-diMeC31;5-MeC31? | 2956.8 | 11,17-diMeC29 | 3125.8 | 15-;13-MeC31 | 3066.1 | **C31:1** |
| 3164.1 | 7,15-diMeC31 | 2966.1 | 7,17-;7,15-diMeC29 | 3153.9 | 11,17-diMeC31 | 3126.8 | 15-;13-MeC31 |
| 3173.5 | 5,15-diMeC31 | 2975.4 | 5,17-diMeC29;3-MeC29? | 3172.6 | 5,15-;5,17-diMeC31 | 3159.5 | 9,19-diMeC31 |
| 3183.8 | 7,11,15-triMeC31 | 2999.7 | x-MeC30? | 3197.8 | 5,11,y-triMeC31? | 3173.5 | 5,15-;5,17-diMeC31 |
| 3196.8 | 5,9,13-triMeC31;C32? | 3066.1 | C31:1 | 3371.5 | 5,15-diMeC33 | 3351.0 | **11,15-diMeC33** |
| 3324.8 | 16-;15-;13-MeC33 | 3127.7 | 15-;13-MeC31 | 3554.6 | **9,17-diMeC35** |  |  |
| 3362.2 | 7,15-diMeC33 | 3138.0 | 7-MeC31 |  |  |  |  |
| 3372.5 | 5,15-diMeC33 | 3151.1 | 13,17-diMeC31 |  |  |  |  |
| 3572.4 | **5,15-;5,17-diMeC35** | 3165.1 | 7,15-diMeC31 |  |  |  |  |
|  |  | 3173.5 | 5,15-;5,17-diMeC31 |  |  |  |  |
|  |  | 3363.1 | 7,17-;7,19-;7,21-diMeC33 |  |  |  |  |

**Tab. S4:** Quality statistics of the near chromosome-level *P. dominula* sensu stricto reference assembly. Contig statistics, gap content and gene-space completeness are from BUSCO version 5.8.2 [57], run against the Hymenoptera lineage dataset; scaffold L50, N90 and L90, GC content, read coverage and taxonomic assignment from BlobToolKit version 4.5.0 [53]; consensus quality and *k*-mer completeness from Merqury version 1.3 [61]. N50 and N90 are the lengths at or above which 50% and 90% of the assembly span is contained, and L50 and L90 the numbers of such scaffolds; contigs are the scaffold segments between runs of ten or more Ns. Completeness categories: C, complete; S, complete and single-copy; D, complete and duplicated; F, fragmented; M, missing. *k*-mer completeness is the share of solid read *k*-mers found in the assembly. Taxonomic assignment is at phylum rank, and the last two rows partition the assembly.

| Statistic | Value |
|---|---|
| Accession | GCA_965197175.2 |
| Sequencing | PacBio HiFi from one haploid male, 111× mean read coverage; Hi-C from a second male |
| Assembly span | 237.7 Mb |
| Scaffolds | 200 |
| Scaffold N50 (L50) | 11.1 Mb (9) |
| Scaffold N90 (L90) | 2.1 Mb (23) |
| Contigs | 1,064 |
| Contig N50 | 511.9 kb |
| Gap (N) content | 0.036% |
| GC content | 30.1% |
| BUSCO completeness, hymenoptera_odb12 (*n* = 5,920) | C: 97.1% [S: 96.8%, D: 0.3%], F: 1.0%, M: 1.9% |
| Consensus Quality Value (QV) | 58.8 (error rate 1.3 × 10<sup>−6</sup>) |
| *k*-mer completeness | 99.2% |
| Assigned to Arthropoda | 213.7 Mb, 89.9% (32 scaffolds) |
| No taxonomic hit | 24.0 Mb, 10.1% (168 scaffolds) |

**Tab. S5:**
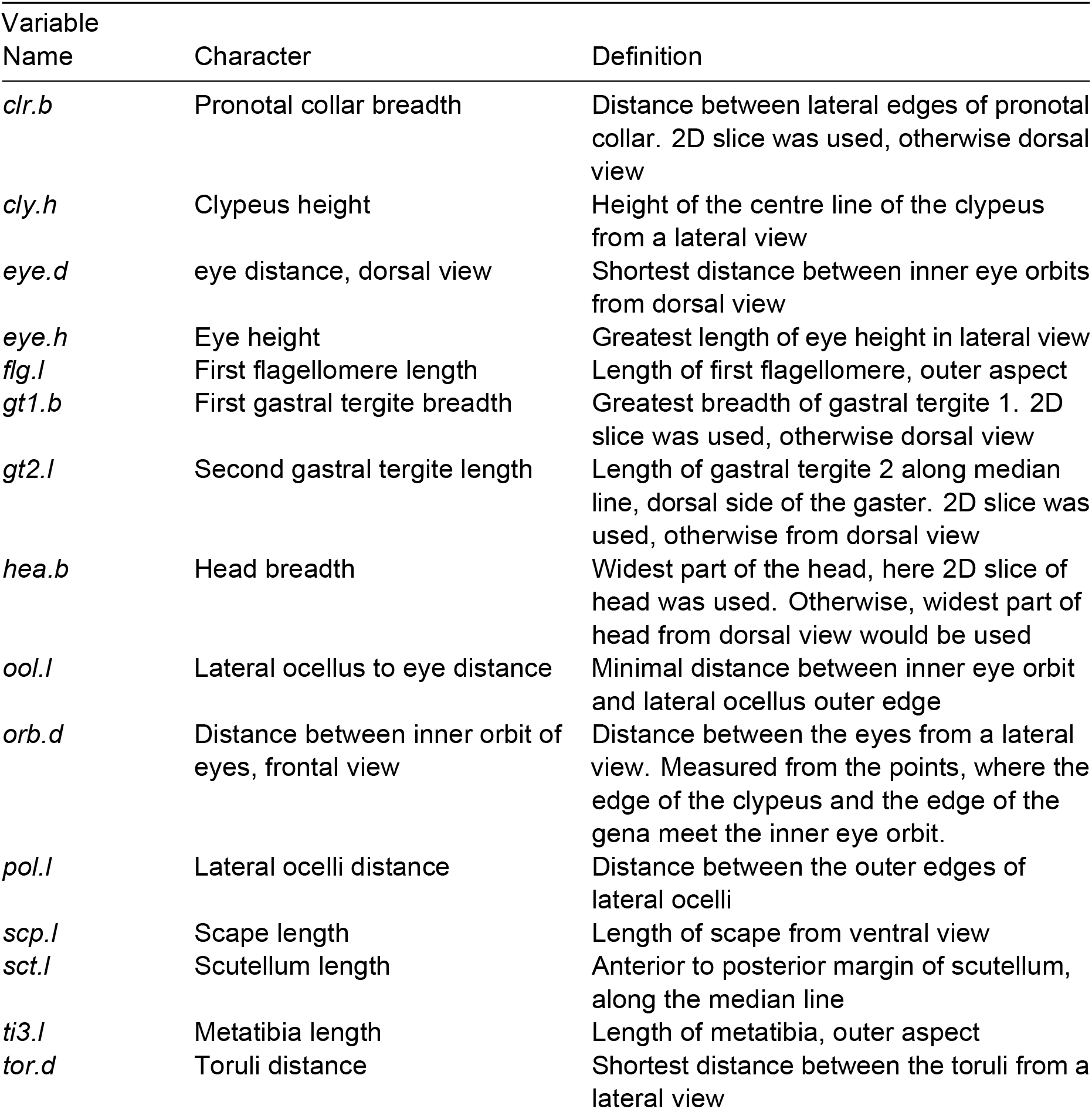
Linear distances (n=15) included as characters for somatic morphometrics of female individuals. The measurements are illustrated on a specimen in Fig. S6.

| Variable Name | Character | Definition |
|---|---|---|
| *clr.b* | Pronotal collar breadth | Distance between lateral edges of pronotal collar. 2D slice was used, otherwise dorsal view |
| *cly.h* | Clypeus height | Height of the centre line of the clypeus from a lateral view |
| *eye.d* | eye distance, dorsal view | Shortest distance between inner eye orbits from dorsal view |
| *eye.h* | Eye height | Greatest length of eye height in lateral view |
| *flg.l* | First flagellomere length | Length of first flagellomere, outer aspect |
| *gt1.b* | First gastral tergite breadth | Greatest breadth of gastral tergite 1. 2D slice was used, otherwise dorsal view |
| *gt2.l* | Second gastral tergite length | Length of gastral tergite 2 along median line, dorsal side of the gaster. 2D slice was used, otherwise from dorsal view |
| *hea.b* | Head breadth | Widest part of the head, here 2D slice of head was used. Otherwise, widest part of head from dorsal view would be used |
| *ool.l* | Lateral ocellus to eye distance | Minimal distance between inner eye orbit and lateral ocellus outer edge |
| *orb.d* | Distance between inner orbit of eyes, frontal view | Distance between the eyes from a lateral view. Measured from the points, where the edge of the clypeus and the edge of the gena meet the inner eye orbit. |
| *pol.l* | Lateral ocelli distance | Distance between the outer edges of lateral ocelli |
| *scp.l* | Scape length | Length of scape from ventral view |
| *sct.l* | Scutellum length | Anterior to posterior margin of scutellum, along the median line |
| *ti3.l* | Metatibia length | Length of metatibia, outer aspect |
| *tor.d* | Toruli distance | Shortest distance between the toruli from a lateral view |

**Tab. S6:** Oligonucleotide primers used for *Wolbachia* MLST and wsp gene amplification.

| Oligo Name | Sequence (5’ → 3’) |
|---|---|
| gatB_F1 | GAK TTA AAY CGY GCA GGB GTT |
| gatB_R1 | TGG YAA YTC RGG YAA AGA TGA |
| coxA_F1 | TTT GRG CRA TYA ACT TTA TAG |
| coxA_R1 | CTA AAG ACT TTK ACR CCA GT |
| hcpA_F1 | GAA ATA RCA GTT GCT GCA AA |
| hcpA_R1 | GAA AGT YRA GCA AGY TCT G |
| ftsZ_F1 | ATY ATG GAR CAT ATA AAR GAT AG |
| ftsZ_R1 | TCR AGY AAT GGA TTR GAT AT |
| fbpA_F1 | GCT GCT CCR CTT GGY WTG AT |
| fbpA_R1 | CCR CCA GAR AAA AYY ACT ATT C |
| wsp_F1 | GTC CAA TAR STG ATG ARG AAA C |
| wsp_R1 | CYG CAC CAA YAG YRC TRT AAA |

**Tab. S7:** Separation of the two barcode groups and association with day of year for each principal component of the female cuticular profile data. AUC is the probability that a randomly chosen BOLD:AAA9495 female scores above a randomly chosen BOLD:AAB7105 female, with 0.5 denoting no separation. Intervals are stratified bootstrap percentiles. Correlations are Pearson coefficients against day of year. Repeating the test against absolute distance from midyear, the covariate used in the N-CPLS models, likewise showed no association for PC3 (*r* = −0.06, *p* = 0.436). Technical replicates were excluded and all *p*-values are Holm-corrected across every test in the underlying analysis.

| Component | AUC (95% CI) | *p* | *r* (day of year) | *p* |
|---|---|---|---|---|
| PC1 | 0.52 (0.47–0.56) | 1.000 | −0.19 | < 0.001 |
| PC2 | 0.61 (0.57–0.65) | < 0.001 | −0.51 | < 0.001 |
| PC3 | 0.97 (0.96–0.99) | < 0.001 | −0.04 | 0.912 |

**Tab. S8:** Detection of the substances marked in bold in Tab. S3, in the cuticular profiles of all females (166 *P. dominula*, 475 *P. pectoralis*) and males (41 and 169), one run per individual. A substance counted as present when each of its detection ions exceeded a detection limit set from chance intensities, irrespective of species membership (Supplementary Methods). RI, Kováts retention index of the annotated peak. Substances in bold are those used to identify specimens in the Taxonomic Designations.

| Sex | Substance | RI | *P. dominula* | *P. pectoralis* |
|---|---|---|---|---|
| ♀ | 7-methylheptacosane (7-MeC27) | 2738.2 | 164 of 166 | 387 of 475 |
| ♀ | 5,15-; 5,17-dimethylpentatriacontane (5,15-;5,17-diMeC35) | 3572.4 | 154 of 166 | 106 of 475 |
| ♀ | **heptacosene (C27:1)** | 2691.5 | 1 of 166 | 350 of 475 |
| ♀ | **nonacosene (C29:1)** | 2893.2 | 0 of 166 | 265 of 475 |
| ♂ | **10,14-dimethyltriacontane (10,14-diMeC30)** | 3056.7 | 41 of 41 | 1 of 169 |
| ♂ | 9,17-dimethylpentatriacontane (9,17-diMeC35) | 3554.6 | 32 of 41 | 0 of 169 |
| ♂ | hentriacontene (C31:1) | 3066.1 | 1 of 41 | 136 of 169 |
| ♂ | 11,15-dimethyltritriacontane (11,15-diMeC33) | 3351.0 | 1 of 41 | 160 of 169 |

**Tab. S9:** Comparison of allele states for private allele panel SNPs in available reference genome assemblies (17,260 shared loci recovered from a panel of 18,026). The table shows the counts of SNPs where the assemblies both show the same allele as Pool A or B, are split between Pool A and B, or show a different allele than either pool.

| GCA_049000335 VS GCF_001465965 | A | B | Other |
|---|---|---|---|
| A | 1641 | 159 | 7 |
| B | 122 | 15312 | 13 |
| Other | 2 | 1 | 3 |

**Tab. S10:** Pairwise mash distances among all available *Polistes* genome assemblies, in per mille (‰; lower triangle shown; values rounded to one decimal place; row and column indices correspond). The first three assemblies belong to the *P. dominula* species complex. Our new assembly (GCA_965197175) and the two previously published assemblies *Pdom_r1.2(USA)* (GCA_001465965) and *Pdom_raw_medaka(NZ)* (GCA_049000335), the latter two both belonging to *P. pectoralis*. Mash distance is a k-mer–based estimate of genome-wide divergence (approximately one minus the average nucleotide identity).

| # | Assembly (accession) | 1 | 2 | 3 | 4 | 5 | 6 | 7 | 8 | 9 |
|---|---|---|---|---|---|---|---|---|---|---|
| 1 | *P. dominula* (GCA_965197175) |  |  |  |  |  |  |  |  |  |
| 2 | *P. pectoralis* (GCA_001465965) | 6.7 |  |  |  |  |  |  |  |  |
| 3 | *P. pectoralis* (GCA_049000335) | 6.5 | 5.0 |  |  |  |  |  |  |  |
| 4 | *P. exclamans* (GCA_023678865) | 81.1 | 81.6 | 81.5 |  |  |  |  |  |  |
| 5 | *P. canadensis* (GCA_001313835) | 81.9 | 82.2 | 82.2 | 16.1 |  |  |  |  |  |
| 6 | *P. metricus* (GCA_010416925) | 80.4 | 81.1 | 80.5 | 49.5 | 50.0 |  |  |  |  |
| 7 | *P. fuscatus* (GCA_010416935) | 80.9 | 81.2 | 80.8 | 50.0 | 50.2 | 8.8 |  |  |  |
| 8 | *P. dorsalis* (GCA_010416905) | 81.0 | 81.5 | 81.0 | 50.1 | 50.5 | 16.3 | 17.2 |  |  |
| 9 | *P. chinensis* (GCA_049000355) | 14.1 | 13.9 | 13.9 | 81.4 | 82.4 | 80.7 | 81.4 | 81.3 |  |
| 10 | *P. biglumis* (GCA_984563175) | 25.2 | 25.3 | 25.4 | 80.7 | 81.4 | 81.2 | 81.5 | 81.8 | 26.2 |

**Tab. S11:** Confusion matrix and per-group accuracies for the Leave-One-Out Cross-Validation (LOOCV) of the Canonical Variate Analysis (CVA) based on the first 19 PC scores. The table shows the number of individuals from each BIN group correctly or incorrectly classified based on wing morphology, along with the overall predictive accuracy of CVA-based predictions.

| True group | Predicted BOLD:AAA9495 | Predicted BOLD:AAB7105 | Correct | Total | Per-group accuracy |
|---|---|---|---|---|---|
| BOLD:AAA9495 | 79 | 18 | 79 | 97 | 81.44% |
| BOLD:AAB7105 | 18 | 85 | 85 | 103 | 82.52% |
| **Total** | **97** | **103** | **164** | **200** | **82.00%** |

**Tab. S12:** Receiver Operating Characteristic (ROC) curve analysis for the four most discriminating character length ratios from the LDA ratio extraction of 3D somatic morphometric data (n = 40). The table shows the Area Under the Curve (AUC), the direction of the ratio for classification, the optimal classification threshold, and the sensitivity and specificity at that threshold. Thresholds are given to about 2% of the ratio’s value, roughly what a measuring eyepiece resolves, and were chosen at that precision.

| Character<br>Ratio | AUC | Direction for *P. dominula* | Threshold | Sensitivity | Specificity |
|---|---|---|---|---|---|
| clr.b/tor.d | 0.99 | lower | 5.5 | 1.00 | 0.85 |
| flg.l/pol.l | 0.98 | higher | 1.48 | 0.85 | 0.95 |
| ool.l/scp.l | 0.82 | higher | 0.42 | 0.75 | 0.75 |
| eye.h/hea.b | 0.59 | higher | 0.64 | 0.90 | 0.25 |

**Tab. S13:** Type specimens designated in this study. The lectotype is one of two suspected syntypes held in the Gerning Collection at the Museum Wiesbaden, both recognised as *P. dominula* by S. Jung using modern identification keys. The second syntype, ON_15401, is not designated and is shown alongside the lectotype in Fig. S5. Collection dates for the Gerning material were not recorded, though 1791 is the year of the original description. The COI sequence of the neotype has been submitted to BOLD. CHC-based identification follows the rule for females (Tab. S14), from the alkenes C27:1 and C29:1 (Tab. S8) and the TICC signal ratio **a/b** of 13-;11-MeC27 to 7-MeC27 (Fig. S8, Fig. S9).

|  | **ON_15402** | **ON_15015** |
|---|---|---|
| Nomenclatural status | **Lectotype** of *P. dominula* (Christ, 1791) | **Neotype** of *P. pectoralis* Herrich-Schäffer, 1841 |
| Sex | ♀ | ♀ |
| Locality | Gerning Collection; original locality not recorded | Germany, Freiburg, Eschholzpark |
| Coordinates | — | 48.000833° N, 7.831457° E |
| Collection date | before 1791 | 2022-06-08 |
| Collector | J. I. Gerning | M. Lehmann |
| Identified by | S. Jung | M. Lehmann |
| Depository | Museum Wiesbaden, Germany | Leibniz Institute for the Analysis of Biodiversity Change, Bonn (LIB) |
| Despository accession | — | ZFMK-TIS-89039 |
| COI BIN | not barcoded | BOLD:AAB7105 |
| COI accession | — | *[pending release]* |
| Species assignment, and basis | *P. dominula*: CHC profile and somatic morphology | *P. pectoralis*: CHC profile, somatic morphology, and COI barcode |
| CHC profile | Fig. 1A (projected) | Fig. 1A (model-fitting) |
| CHC identification | neither alkene, TICC signal ratio **a/b** 0.19: *P. dominula* | C27:1 and C29:1, TICC signal ratio **a/b** 2.42: *P. pectoralis* |
| µCT surface rendering | Fig. S5C/D | scanned, not figured |
| Somatic morphometrics | **clr.b/tor.d** 5.2, **flg.l/pol.l** 1.51 (Fig. 3) | **clr.b/tor.d** 5.8, **flg.l/pol.l** 1.40 (Fig. 3) |

**Tab. S14:** Characters and rules for identifying *P. dominula* and *P. pectoralis*. No single character separates the two species without exception, so specimens are identified by agreement. Each line of evidence gives *P. dominula*, *P. pectoralis* or undetermined, and a specimen is identified when at least one line decides and none points to the other species. Counts are individuals with the state among all individuals of that species with data for the line; specimens identified as the other species by any line: none. Species membership corresponds to BIN, except for rare *P. dominula* carrying BOLD:AAB7105 through mitochondrial introgression (Fig. 2). Substances count as present above a detection limit (Tab. S8) and both alkenes are minor peaks and can fall below it in dilute extracts. **TICC signal ratios**: Fig. S8, Fig. S10. **Somatic ratios**: Fig. S6. The females with both lines had been selected on whether C27:1 could be detected (Materials and Methods).

| Line of evidence and character | *P. dominula* | *P. pectoralis* |
|---|---|---|
| **Cuticular hydrocarbons, females** |  |  |
| C27:1 (RI 2691.5) or C29:1 (RI 2893.2) | neither (165 of 166) | at least one (368 of 475) |
| 13-;11-MeC27 to 7-MeC27 TICC signal ratio | ≤ 1.67 (153 of 166) | > 1.67 (474 of 475) |
| *Identified when both agree* | 152 of 166 | 367 of 475 |
| **Cuticular hydrocarbons, males** |  |  |
| 10,14-diMeC30 (RI 3056.7) | present (41 of 41) | absent (168 of 169) |
| 11,15-diMeC33 to 5,15-diMeC33 TICC signal ratio | ≤ 0.22 (41 of 41) | > 0.22 (168 of 169) |
| *Identified when both agree* | 41 of 41 | 167 of 169 |
| **Somatic proportions, females** |  |  |
| **clr.b/tor.d** | < 5.4 (15 of 20) | > 5.5 (16 of 20) |
| **flg.l/pol.l** | > 1.48 (16 of 20) | < 1.46 (14 of 20) |
| *Identified when at least one points to the species and neither to the other* | 19 of 20 | 18 of 20 |
| **COI barcode** |  |  |
| Barcode Index Number | likely BOLD:AAA9495 | BOLD:AAB7105 |
| **Both lines, females with cuticular and somatic data** |  |  |
| *Identified* | 20 of 20 | 20 of 20 |

## Supplementary Files

**Supplementary File 1: Sample collection data.** This file contains available sourcing and collection data for all samples used in this study. The inclusion of a sample in a specific experiment is listed with boolean flags for each of the following experiments: reference genome sequencing and assembly, COI barcoding, CHC chemometrics, somatic morphometrics, wing morphometrics, *Wolbachia* sequence typing, as well as genomic Pool-Seq and individual genome sequencing.

**Supplementary File 2: Female outlier-scan results.** This file contains the female outlierscan table for the fixed two-latent-variable N-CPLS-DA model, based on 1,000 reshuffled five-fold outer hold-out tests. For each female sample, it reports the input-data sample index, sample identifier, BOLD class, the number of times the sample appeared in an outer test fold (n_tested), the number of times it was flagged as an outlier by held-out misclassification (n_flagged), and the resulting misassignment ratio (n_flagged / n_tested). The table is sorted by decreasing misassignment ratio and includes all female samples, including samples that were never flagged.

**Supplementary File 3: Male outlier-scan results.** This file contains the male outlier-scan table for the fixed one-latent-variable N-CPLS-DA model, based on 1,000 reshuffled five-fold outer hold-out tests. It reports the same fields as Supplementary File 2 for each male sample, is sorted by decreasing misassignment ratio, and includes all male samples, including samples that were never flagged.

**Supplementary File 4: External BOLD records.** This file contains the records identified as *Polistes dominula* held in the Barcode of Life Data System for the United States and New Zealand, retrieved on 14 September 2026 and reproduced as exported. It is the source of the record counts given in the Supplementary Results. One further US record was excluded because its submitted sequence is too short to be assigned to a BIN.

## Notes

https://github.com/oniehuis/JuChrom.jl

https://github.com/oniehuis/NCPLS.jl

https://github.com/git-hobo/popoolation2/tree/master/workflow

### Competing Interest Statement

The authors have declared no competing interest.

## References

1. Field J, Leadbeater E. 2016 Cooperation between non-relatives in a primitively eusocial paper wasp, Polistes dominula. Philosophical Transactions of the Royal Society B: Biological Sciences 371, 20150093. (doi:10.1098/rstb.2015.0093)

2. West-Eberhard M. 1969 The Social Biology of Polistine Wasps. Miscellaneous Publications Museum of Zoology 140, 1–101.

3. Reeve HK, Nonacs P. 1992 Social contracts in wasp societies. Nature 359, 823–825. (doi:10.1038/359823a0)

4. Queller DC, Strassmann JE. 1998 Kin Selection and Social Insects. BioScience 48, 165–175. (doi:10.2307/1313262)

5. Tibbetts EA, Pardo-Sanchez J, Ramirez-Matias J, Avarguès-Weber A. 2021 Individual recognition is associated with holistic face processing in Polistes paper wasps in a species-specific way. Proceedings of the Royal Society B: Biological Sciences 288, 20203010. (doi:10.1098/rspb.2020.3010)

6. Cini A, Cappa F, Pepiciello I, Platania L, Dapporto L, Cervo R. 2019 Sight in a clique, scent in society: Plasticity in the use of nestmate recognition cues along colony development in the social wasp *Polistes dominula*. Frontiers in Ecology and Evolution 7. (doi:10.3389/fevo.2019.00444)

7. Pekkarinen A, Gustafsson B. 1999 The Polistes species in northern Europe (Hymenoptera: Vespidae). Entomologica Fennica 10, 191–197. (doi:10.33338/ef.84021)

8. Höcherl N, Tautz J. 2015 Nesting behavior of the paper wasp *Polistes dominula* in Central Europe—a flexible system for expanding into new areas. Ecosphere 6, 1–11. (doi:10.1890/ES15-00254.1)

9. Kovac H, Nagy JM, Käfer H, Stabentheiner A. 2023 Relationship between Nest and Body Temperature and Microclimate in the Paper Wasp Polistes dominula. Insects 14, 886. (doi:10.3390/insects14110886)

10. Howse MWF, Haywood J, Lester PJ. 2020 Bioclimatic Modelling Identifies Suitable Habitat for the Establishment of the Invasive European Paper Wasp (Hymenoptera: Vespidae) across the Southern Hemisphere. Insects 11, 784. (doi:10.3390/insects11110784)

11. Ratnasingham S, Hebert PDN. 2013 A DNA-based registry for all animal species: The barcode index number (BIN) system. PLOS One 8, e66213. (doi:10.1371/journal.pone.0066213)

12. Schmid-Egger C, Van Achterberg K, Neumeyer R, Morinière J, Schmidt S. 2017 Revision of the West Palaearctic *Polistes* Latreille, with the descriptions of two species – an integrative approach using morphology and DNA barcodes (Hymenoptera, Vespidae). ZooKeys 713, 53–112. (doi:10.3897/zookeys.713.11335)

13. Neumeyer R, Baur H, Guex G-D, Praz C. 2014 A new species of the paper wasp genus Polistes (Hymenoptera, Vespidae, Polistinae) in Europe revealed by morphometrics and molecular analyses. ZooKeys 400, 67–118. (doi:10.3897/zookeys.400.6611)

14. Junker M et al. 2015 Three in One—Multiple Faunal Elements within an Endangered European Butterfly Species. PLOS ONE 10, e0142282. (doi:10.1371/journal.pone.0142282)

15. Martínez-Rodríguez P, Bella JL. 2018 Chorthippus parallelus and Wolbachia: Overlapping Orthopteroid and Bacterial Hybrid Zones. Frontiers in Genetics 9. (doi:10.3389/fgene.2018.00604)

16. Darnis M, Ravigné V, Sauvion N. 2025 Origin, divergence and migration routes of psyllids of the *Cacopsylla pruni* complex (Hemiptera: Psylloidea) inferred by approximate Bayesian computation methods. Journal of Biogeography 52, e15177. (doi:10.1111/jbi.15177)

17. Cutter AD. 2024 Beyond Haldane’s rule: Sex-biased hybrid dysfunction for all modes of sex determination. eLife 13, e96652. (doi:10.7554/eLife.96652)

18. Gibson JD, Niehuis O, Peirson BRE, Cash EI, Gadau J. 2013 Genetic and developmental basis of F2 hybrid breakdown in *Nasonia* parasitoid wasps. Evolution 67, 2124–2132. (doi:10.1111/evo.12080)

19. Kulmuni J, Seifert B, Pamilo P. 2010 Segregation distortion causes large-scale differences between male and female genomes in hybrid ants. Proceedings of the National Academy of Sciences 107, 7371–7376. (doi:10.1073/pnas.0912409107)

20. Werren JH. 1997 Biology of *Wolbachia*. Annual Review of Entomology 42, 587–609. (doi:10.1146/annurev.ento.42.1.587)

21. Stahlhut JK, Liebert AE, Starks PT, Dapporto L, Jaenike J. 2006 *Wolbachia* in the invasive European paper wasp *Polistes dominulus*. Insectes Sociaux 53, 269–273. (doi:10.1007/s00040-006-0868-7)

22. Graham RI, Wilson K. 2012 Male-killing *Wolbachia* and mitochondrial selective sweep in a migratory African insect. BMC Evolutionary Biology 12, 204. (doi:10.1186/1471-2148-12-204)

23. Singer TL, Espelie KE. 1992 Social wasps use nest paper hydrocarbons for nestmate recognition. Animal Behaviour 44, 63–68. (doi:10.1016/S0003-3472(05)80755-9)

24. Dani FR, Jones GR, Destri S, Spencer SH, Turillazzi S. 2001 Deciphering the recognition signature within the cuticular chemical profile of paper wasps. Animal Behaviour 62, 165–171. (doi:10.1006/anbe.2001.1714)

25. Brito JH dos S, Antonialli-Junior WF, Montagna T dos S, Mendonça A, Sguarizi-Antonio D, Súarez YR, Lima SM, Andrade LH da C, Cardoso CAL. 2017 Linear alkanes and reproductive status of Polistes versicolor (Hymenoptera: Vespidae) females in winter aggregates. Sociobiology 64, 327–333. (doi:10.13102/sociobiology.v64i3.1039)

26. Sledge MF, Dani FR, Cervo R, Dapporto L, Turillazzi S. 2001 Recognition of social parasites as nest-mates: Adoption of colony-specific host cuticular odours by the paper wasp parasite *Polistes sulcifer* . Proceedings of the Royal Society of London. Series B: Biological Sciences 268, 2253–2260. (doi:10.1098/rspb.2001.1799)

27. Beani L, Bagnères A-G, Elia M, Petrocelli I, Cappa F, Lorenzi MC. 2019 Cuticular hydrocarbons as cues of sex and health condition in Polistes dominula wasps. Insectes Sociaux 66, 543–553. (doi:10.1007/s00040-019-00721-z)

28. Da Silva RC, Van Meerbeeck L, Do Nascimento FS, Wenseleers T, Oi CA. 2021 Closerange cues used by males of *Polistes dominula* in sex discrimination. The Science of Nature 108, 15. (doi:10.1007/s00114-021-01730-4)

29. Cervo R. 2006 Polistes wasps and their social parasites: An overview. Annales Zoologici Fennici 43, 531–549.

30. Cervo R, Stemmer C, Castle WA, Queller DC, Strassmann JE. 2004 Social parasitism of *Polistes dominulus* by *Polistes nimphus* (Hymenoptera, Vespidae). Insectes Sociaux 51, 101–108. (doi:10.1007/s00040-003-0717-x)

31. Lorenzi MC, Bagnères AG. 2002 Concealing identity and mimicking hosts: A dual chemical strategy for a single social parasite? (Polistes atrimandibularis, Hymenoptera: Vespidae). Parasitology 125, 507–512. (doi:10.1017/S003118200200238X)

32. Christ JL. 1791 Die Wespen. In Naturgeschichte, Klassification und Nomenclatur der Insekten vom Bienen, Wespen und Ameisengeschlecht : Als der fünften Klasse fünfte Ordnung des Linneischen Natursystems von den Insekten, Hymenoptera : Mit häutigen Flügeln, p. 229. Frankfurt am Main: in der Hermannischen Buchhandlung. (doi:10.5962/bhl.title.87724)

33. Bezanson J, Edelman A, Karpinski S, Shah VB. 2017 Julia: A fresh approach to numerical computing. SIAM Review 59, 65–98. (doi:10.1137/141000671)

34. Niehuis O. 2026 JuChrom.jl.

35. O’Callaghan S et al. 2012 PyMS: A Python toolkit for processing of gas chromatographymass spectrometry (GC-MS) data. Application and comparative study of selected tools. BMC Bioinformatics 13, 115. (doi:10.1186/1471-2105-13-115)

36. Kováts E. 1958 Gas-chromatographische Charakterisierung organischer Verbindungen. Teil 1: Retentionsindices aliphatischer Halogenide, Alkohole, Aldehyde und Ketone. Helvetica Chimica Acta 41, 1915–1932. (doi:10.1002/hlca.19580410703)

37. Baek S-J, Park A, Ahn Y-J, Choo J. 2015 Baseline correction using asymmetrically reweighted penalized least squares smoothing. The Analyst 140, 250–257. (doi:10.1039/C4AN01061B)

38. Aitchison J. 1986 The statistical analysis of compositional data. London ; New York: Chapman and Hall.

39. Liland KH, Indahl UG, Skogholt J, Mishra P. 2022 The canonical partial least squares approach to analysing multiway datasets—N-CPLS. Journal of Chemometrics 36, e3432. (doi:10.1002/cem.3432)

40. Niehuis O. 2026 NCPLS.jl.

41. Smit S, Van Breemen MJ, Hoefsloot HCJ, Smilde AK, Aerts JMFG, De Koster CG. 2007 Assessing the statistical validity of proteomics based biomarkers. Analytica Chimica Acta 592, 210–217. (doi:10.1016/j.aca.2007.04.043)

42. Phipson B, Smyth GK. 2010 Permutation P-values should never be zero: Calculating exact P-values when permutations are randomly drawn. Statistical Applications in Genetics and Molecular Biology 9, Article39. (doi:10.2202/1544-6115.1585)

43. Danisch S, Krumbiegel J. 2021 Makie.jl: Flexible high-performance data visualization for Julia. Journal of Open Source Software 6, 3349. (doi:10.21105/joss.03349)

44. Inkscape Team. 2020 Inkscape: An open-source vector graphics editor.

45. Astrin JJ, Stüben PE. 2008 Phylogeny in cryptic weevils: Molecules, morphology and new genera of western Palaearctic Cryptorhynchinae (Coleoptera: Curculionidae). Invertebrate Systematics 22, 503–522. (doi:10.1071/IS07057)

46. Okonechnikov K, Golosova O, Fursov M, The Ugene Team. 2012 Unipro UGENE: A unified bioinformatics toolkit. Bioinformatics 28, 1166–1167. (doi:10.1093/bioinformatics/bts091)

47. Buchner D, Leese F. 2020 BOLDigger – a Python package to identify and organise sequences with the Barcode of Life Data Systems. Metabarcoding and Metagenomics 4, e53535. (doi:10.3897/mbmg.4.53535)

48. Baid G et al. 2023 DeepConsensus improves the accuracy of sequences with a gapaware sequence transformer. Nature Biotechnology 41, 232–238. (doi:10.1038/s41587-022-01435-7)

49. Martin M. 2011 Cutadapt removes adapter sequences from high-throughput sequencing reads. EMBnet.journal 17, 10. (doi:10.14806/ej.17.1.200)

50. Uliano-Silva M et al. 2023 MitoHiFi: A Python pipeline for mitochondrial genome assembly from PacBio high fidelity reads. BMC Bioinformatics 24, 288. (doi:10.1186/s12859-023-05385-y)

51. Cheng H, Concepcion GT, Feng X, Zhang H, Li H. 2021 Haplotype-resolved de novo assembly using phased assembly graphs with hifiasm. Nature Methods 18, 170–175. (doi:10.1038/s41592-020-01056-5)

52. Zhou C, McCarthy SA, Durbin R. 2023 YaHS: Yet another Hi-C scaffolding tool. Bioinformatics 39, btac808. (doi:10.1093/bioinformatics/btac808)

53. Challis R, Richards E, Rajan J, Cochrane G, Blaxter M. 2020 BlobToolKit – Interactive Quality Assessment of Genome Assemblies. G3 Genes|Genomes|Genetics 10, 1361–1374. (doi:10.1534/g3.119.400908)

54. Smit A, Hubley R. 2024 RepeatMasker.

55. Bao W, Kojima KK, Kohany O. 2015 Repbase Update, a database of repetitive elements in eukaryotic genomes. Mobile DNA 6, 11. (doi:10.1186/s13100-015-0041-9)

56. Flynn JM, Hubley R, Goubert C, Rosen J, Clark AG, Feschotte C, Smit AF. 2020 RepeatModeler2 for automated genomic discovery of transposable element families. Proceedings of the National Academy of Sciences 117, 9451–9457. (doi:10.1073/pnas.1921046117)

57. Simão FA, Waterhouse RM, Ioannidis P, Kriventseva EV, Zdobnov EM. 2015 BUSCO: Assessing genome assembly and annotation completeness with single-copy orthologs. Bioinformatics 31, 3210–3212. (doi:10.1093/bioinformatics/btv351)

58. Huang N, Li H. 2023 compleasm: A faster and more accurate reimplementation of BUSCO. Bioinformatics 39, btad595. (doi:10.1093/bioinformatics/btad595)

59. Gurevich A, Saveliev V, Vyahhi N, Tesler G. 2013 QUAST: Quality assessment tool for genome assemblies. Bioinformatics 29, 1072–1075. (doi:10.1093/bioinformatics/btt086)

60. Pfenninger M, Schönnenbeck P, Schell T. 2022 ModEst: Accurate estimation of genome size from next generation sequencing data. Molecular Ecology Resources 22, 1454–1464. (doi:10.1111/1755-0998.13570)

61. Rhie A, Walenz BP, Koren S, Phillippy AM. 2020 Merqury: Reference-free quality, completeness, and phasing assessment for genome assemblies. Genome Biology 21, 245. (doi:10.1186/s13059-020-02134-9)

62. Standage DS, Berens AJ, Glastad KM, Severin AJ, Brendel VP, Toth AL. 2016 Genome, transcriptome and methylome sequencing of a primitively eusocial wasp reveal a greatly reduced DNA methylation system in a social insect. Molecular Ecology 25, 1769–1784. (doi:10.1111/mec.13578)

63. Gilligan J et al. 2024 Rapid and cost-effective production of high-quality insect genome sequences. (doi:10.1101/2024.10.06.616902)

64. Li H. 2018 minimap2: Pairwise alignment for nucleotide sequences. Bioinformatics 34, 3094–3100. (doi:10.1093/bioinformatics/bty191)

65. Ondov BD, Treangen TJ, Melsted P, Mallonee AB, Bergman NH, Koren S, Phillippy AM. 2016 Mash: Fast genome and metagenome distance estimation using MinHash. Genome Biology 17, 132. (doi:10.1186/s13059-016-0997-x)

66. Bolger AM, Lohse M, Usadel B. 2014 Trimmomatic: A flexible trimmer for Illumina sequence data. Bioinformatics 30, 2114–2120. (doi:10.1093/bioinformatics/btu170)

67. Li H. 2013 Aligning sequence reads, clone sequences and assembly contigs with BWA-MEM. (doi:10.48550/arXiv.1303.3997)

68. Li H et al. 2009 The sequence alignment/map format and SAMtools. Bioinformatics 25, 2078–2079. (doi:10.1093/bioinformatics/btp352)

69. Kofler R, Pandey RV, Schlötterer C. 2011 PoPoolation2: Identifying differentiation between populations using sequencing of pooled DNA samples (pool-seq). Bioinformatics 27, 3435–3436. (doi:10.1093/bioinformatics/btr589)

70. Goeman JJ, Solari A. 2014 Multiple hypothesis testing in genomics. Statistics in Medicine 33, 1946–1978. (doi:10.1002/sim.6082)

71. Storey JD, Bass AJ, Dabney A. 2017 qvalue. (doi:10.18129/B9.bioc.qvalue)

72. R Core Team. 2021 R: A language and environment for statistical computing.

73. Posit Team. 2025 RStudio: Integrated development environment for R.

74. Turner S. 2018 qqman: An R package for visualizing GWAS results using Q-Q and Manhattan plots. The Journal of Open Source Software (doi:10.21105/joss.00731)

75. Li H. 2011 A statistical framework for SNP calling, mutation discovery, association mapping and population genetical parameter estimation from sequencing data. Bioinformatics 27, 2987–2993. (doi:10.1093/bioinformatics/btr509)

76. Helicon Remote Team. 2022 Helicon Remote.

77. Lightroom Team. 2022 Adobe Lightroom Classic.

78. Photoshop Team. 2022 Adobe Photoshop.

79. Rohlf FJ. 2010 tpsUtil, file utility program.

80. Rohlf FJ. 2006 tpsDig2, digitize landmarks and outlines.

81. Rohlf FJ. 2015 The tps series of software. Hystrix, the Italian Journal of Mammalogy 26, 1–4. (doi:10.4404/hystrix-26.1-11264)

82. Bustamante T, Fuchs S, Grünewald B, Ellis JD. 2021 A geometric morphometric method and web application for identifying honey bee species (Apis spp.) using only forewings. Apidologie 52, 697–706. (doi:10.1007/s13592-021-00857-7)

83. Francoy TM, Wittmann D, Drauschke M, Müller S, Steinhage V, Bezerra-Laure MAF, Jong DD, Gonçalves LS. 2008 Identification of Africanized honey bees through wing morphometrics: Two fast and efficient procedures. Apidologie 39, 488–494. (doi:10.1051/apido:2008028)

84. Klingenberg CP. 2011 MorphoJ: An integrated software package for geometric morphometrics. Molecular Ecology Resources 11, 353–357. (doi:10.1111/j.1755-0998.2010.02924.x)

85. Katzke J et al. 2026 High-throughput phenomics of global ant biodiversity. Nature Methods 23, 663–672. (doi:10.1038/s41592-026-03005-0)

86. Cecilia A et al. 2025 The IMAGE beamline at the KIT Light Source. Journal of Synchrotron Radiation 32, 1036–1051. (doi:10.1107/S1600577525003777)

87. Vogelgesang M, Faragó T, Morgeneyer TF, Helfen L, dos Santos Rolo T, Myagotin A, Baumbach T. 2016 Real-time image-content-based beamline control for smart 4D X-ray imaging. Journal of Synchrotron Radiation 23, 1254–1263. (doi:10.1107/S1600577516010195)

88. Faragó T, Gasilov S, Emslie I, Zuber M, Helfen L, Vogelgesang M, Baumbach T. 2022 tofu: A fast, versatile and user-friendly image processing toolkit for computed tomography. Journal of Synchrotron Radiation 29, 916–927. (doi:10.1107/S160057752200282X)

89. Amira Team. 2022 Amira.

90. Comet Technologies Canada Inc. 2022 Dragonfly 3D world.

91. Wurdack M, Herbertz S, Dowling D, Kroiss J, Strohm E, Baur H, Niehuis O, Schmitt T. 2015 Striking cuticular hydrocarbon dimorphism in the mason wasp *Odynerus spinipes* and its possible evolutionary cause (Hymenoptera: Chrysididae, Vespidae). Proceedings of the Royal Society B: Biological Sciences 282, 20151777. (doi:10.1098/rspb.2015.1777)

92. Baur H, Leuenberger C. 2020 Multivariate Ratio Analysis (MRA): R-scripts and tutorials for calculating Shape PCA, Ratio Spectra, and LDA Ratio Extractor. (doi:10.5281/zenodo.3890195)

93. Baldo L et al. 2006 Multilocus sequence typing system for the endosymbiont *Wolbachia pipientis*. Applied and Environmental Microbiology 72, 7098–7110. (doi:10.1128/AEM.00731-06)

94. Jolley KA, Bray JE, Maiden MCJ. 2018 Open-access bacterial population genomics: BIGSdb software, the PubMLST.org website and their applications. Wellcome OPEN Research 3, 124. (doi:10.12688/wellcomeopenres.14826.1)

95. International Commission on Zoological Nomenclature. 1999 International code of zoological nomenclature. 4th edn. London: International Trust for Zoological Nomenclature.

96. Herrich-Schäffer GAW. 1841 Synopsis Generis POLISTES. In Faunae insectorum Germanicae initia, oder Deutschlands Insecten, p. 39. Regensburg: G. J. Manz.

97. Mayr E. 1949 Systematics and the origin of species from the viewpoint of a zoologist. New York: Columbia University Press.

98. Menzel F, Zumbusch M, Feldmeyer B. 2018 How ants acclimate: Impact of climatic conditions on the cuticular hydrocarbon profile. Functional Ecology 32, 657–666. (doi:10.1111/1365-2435.13008)

99. Holze H, Schrader L, Buellesbach J. 2021 Advances in deciphering the genetic basis of insect cuticular hydrocarbon biosynthesis and variation. Heredity 126, 219–234. (doi:10.1038/s41437-020-00380-y)

100. Servedio MR, Noor MAF. 2003 The role of reinforcement in speciation: Theory and data. Annual Review of Ecology, Evolution, and Systematics 34, 339–364. (doi:10.1146/an-nurev.ecolsys.34.011802.132412)

101. Dietz L et al. 2023 Standardized nuclear markers improve and homogenize species delimitation in Metazoa. Methods in Ecology and Evolution 14, 543–555. (doi:10.1111/2041-210X.14041)

102. Moris VC et al. 2021 Cuticular hydrocarbons on old museum specimens of the spiny mason wasp, Odynerus spinipes (Hymenoptera: Vespidae: Eumeninae), shed light on the distribution and on regional frequencies of distinct chemotypes. Chemoecology 31, 311–322. (doi:10.1007/s00049-021-00350-8)

103. Masaquiza D, Ferrán MO, Guamán S, Naranjo E, Vaca M, Curbelo LM, Arenal A. 2023 Geometric Morphometric Analysis of Wing Shape to Identify Populations of Apis mellifera in Camagüey, Cuba. Insects 14, 306. (doi:10.3390/insects14030306)

104. Ratnasingham S, Hebert PDN. 2007 BOLD: The Barcode of Life Data System (http://www.barcodinglife.org). Molecular Ecology Notes 7, 355–364. (doi:10.1111/j.1471-8286.2007.01678.x)

105. Saito F, Kojima J, Nguyen LTP, Kanuka M. 2007 Polistes formosanus Sonan, 1927 (Hymenoptera: Vespidae), a Good Species Supported by both Morphological and Molecular Phylogenetic Analyses, and a Key Social Wasp in Understanding the Historical Biogeography of the Nansei Islands. Zoological Science 24, 927–939. (doi:10.2108/zsj.24.927)

